# Improving imputation quality in BEAGLE for crop and livestock data

**DOI:** 10.1101/577338

**Authors:** T. Pook, M. Mayer, J. Geibel, S. Weigend, D. Cavero, C.C. Schoen, H. Simianer

**Author notes:** T. Pook; University of Goettingen, Department of Animal Sciences, Center for Integrated Breeding Research, Animal Breeding and Genetics Group, Albrecht-Thaer-Weg 3, 37075 Goettingen, Germany.

## Abstract

Imputation is one of the key steps in the preprocessing and quality control protocol of any genetic study. Most imputation algorithms were originally developed for the use in human genetics and thus are optimized for a high level of genetic diversity. Different versions of BEAGLE were evaluated on genetic datasets of doubled haploids of two European maize landraces, a commercial breeding line and a diversity panel in chicken, respectively, with different levels of genetic diversity and structure which can be taken into account in BEAGLE by parameter tuning. Especially for phasing BEAGLE 5.0 outperformed the newest version (5.1) which in turn also lead to improved imputation. Earlier versions were far more dependent on the adaption of parameters in all our tests. For all versions, the parameter ne (effective population size) had a major effect on the error rate for imputation of ungenotyped markers, reducing error rates by up to 98.5%. Further improvement was obtained by tuning of the parameters affecting the structure of the haplotype cluster that is used to initialize the underlying Hidden Markov Model of BEAGLE. The number of markers with extremely high error rates for the maize datasets were more than halved by the use of a flint reference genome (F7, PE0075 etc.) instead of the commonly used B73. On average, error rates for imputation of ungenotyped markers were reduced by 8.5% by excluding genetically distant individuals from the reference panel for the chicken diversity panel. To optimize imputation accuracy one has to find a balance between representing as much of the genetic diversity as possible while avoiding the introduction of noise by including genetically distant individuals.

## INTRODUCTION

Imputation is one of the key steps in preprocessing genetic data generated by SNP-chips or DNA sequencing, as follow-up applications like genomic prediction (Meuwissen *et al*. 2001) often do not allow for missing values. In some applications the use of a higher marker density can lead to better results even though individuals were not genotyped for most markers (e.g. in genome-wide association studies previously not identified regions can be detected (Yan *et al*. 2017)).

The imputation of genotype data was first introduced by Li and Stephens (2003). The basic idea of the algorithm is the fitting of a Hidden Markov Model (HMM, (Baum and Petrie 1966; Rabiner 1989)) to the sequence of alleles of a haplotype. Over the years, a wide variety of tools with similar basic frameworks, but improvements to the computational efficiency for larger datasets (Howie *et al*. 2009), reference panels (Browning *et al*. 2018) or modifications for improved modeling have been developed. Among others, improvements to the modeling include the use of coalescent trees (Marchini *et al*. 2007), haplotype clusters (Scheet and Stephens 2006) and pre-phasing steps (Scott *et al*. 2007; Howie *et al*. 2012; Loh *et al*. 2016).

To account for the specific structure of livestock and crop datasets, special tools for both cases have been developed. As fully homozygous lines are commonly present in crops, the software TASSEL (Bradbury *et al*. 2007) was developed to work well on this data structure (Swarts *et al*. 2014). Since pedigrees in animal breeding can be much denser than in human populations (both w.r.t. depth and family size), tools like FImpute (Sargolzaei *et al*. 2014) and AlphaImpute (Hickey *et al*. 2011) have been developed to fully utilize this information.

In the imputation process all those methods use the fact that physically close markers are likely inherited together, resulting in nonrandom associations of alleles. These methods thereby rely on the knowledge of the physical position or at least the order of markers for modeling linkage and thus the resulting linkage disequilibrium (LD). In contrast, the software LinkImpute (Money *et al*. 2015) accounts for LD between pairs of markers and not their physical positions. This can be particularly relevant for species in which no reference sequence is available or whose genomes are known for a high amount of translocations and inversions.

In contrast to other methods using a HMM, the Markov chain in BEAGLE is not initialized by the genotypes or haplotypes themselves, but instead the genetic dataset is used to initialize a haplotype cluster (Browning and Browning 2007), which subsequently initializes the HMM. In essence, imputation is then performed by identifying the most likely path through the haplotype cluster based on the non-missing genotypes. As BEAGLE was originally developed for application in human genetics, default settings are chosen to work well for imputation in outbred human populations. Nevertheless, the user still has considerable flexibility to tune the algorithm to the specific genetic structure of the respective dataset. As imputation is usually just a step in the preprocessing and quality control protocol, authors tend to use the default settings of a recent version of some imputation software.

To increase the operational marker density via imputation an additional dataset (reference panel) that is genotyped under a higher density can be used. With increasing computational power and more efficient methods available the common advice here is to use as many individuals as possible to get a good representation of the population (Zhang *et al*. 2013; Browning *et al*. 2018).

In this paper, we compare different BEAGLE versions (4.0 / 4.1 / 5.0 / 5.1) and perform bench-marking tests in regard to imputation quality on virtually all parameters in BEAGLE for a variety of livestock and crop datasets, as it is one of the most frequently used tools in both animal and plant breeding and a new version of the tool has been recently published (Browning *et al*. 2018). We further evaluate which individuals to include in a reference panel when aiming at increasing the marker density of a dataset.

Since imputation algorithms like BEAGLE rely on the assumed physical order of markers, the used reference genome influences the imputation quality. Recently, a variety of new maize reference genomes have been made public (Unterseer *et al*. 2017). We here compare the imputation performance of the commonly used B73v4 (Schnable *et al*. 2009; Jiao *et al*. 2017) and new reference genomes from flint lines in maize that should be genetically closer to our material. To this day, all reference genomes derived in chicken were generated based on an inbred Red Jungle Fowl (*Gallus gallus gallus*; (International Chicken Genome Sequencing Consortium 2004; Bellott *et al*. 2010).

## MATERIALS AND METHODS

### Genotype data used

In the following, we will consider genotypic data of 910 doubled haploid (DH) lines of two European maize (*Zea mays*) landraces (*n* = 501 Kemater Landmais Gelb (KE) and *n* = 409 Petkuser Ferdinand Rot (PE), (Hölker *et al*. 2019)) genotyped using the 600k Affymetrix® Axiom® Maize Array (Unterseer *et al*. 2014). Markers were filtered for being assigned to the highest quality class (Poly High Resolution (Pirani *et al*. 2013)), having a callrate of at least 90%, and for having at most 5% heterozygous calls, as no heterozygous calls are expected for DH lines. The remaining heterozygous calls were set to NA and subsequently imputed using BEAGLE 4.0 with *nsamples* = 50, resulting in a dataset of 501,124 markers with known haplotype phases.

We further considered two chicken (*Gallus gallus*) datasets genotyped with the 580k SNP Affymetrix® Axiom® Genome-Wide Chicken Genotyping Array (Kranis *et al*. 2013). Firstly, a chicken diversity panel containing 1,810 chicken of 82 breeds including Asian types, European types, wild types, commercial broilers and layers (Weigend *et al*. 2014; Malomane *et al*. 2019). Secondly, a dataset containing 888 chicken of a commercial breeding program from Lohmann Tierzucht GmbH. For quality control SNPs / animals with less than 99% / 95% callrate were removed. We will here focus on chromosome 1, 7 and 20 with 56,773 / 65,177, 12,585 / 13,533 and 5,539 / 5,940 SNPs representing cases for large, medium and small size chromosomes. Remaining missing genotypes for both chicken panels were imputed using BEAGLE 4.1 default.

For tests regarding imputation of ungenotyped markers in maize we used the overlapping markers (45,655 SNPs) of the Illumina® MaizeSNP50 BeadChip chip (Ganal *et al*. 2011) as a smaller SNP array. As there is no similar public smaller array with a majority of overlapping markers for the chicken panels, we simply used a subset of every tenth marker. All tests regarding imputation quality were performed on imputed datasets. This might favor the respective method used for the imputation. As the missingness in the maize data (1.20%), diversity panel (0.27%) and commercial chicken breeding line (0.32%) were low in the raw data, this effect should only be minor and is neglected here.

To assess the genetic diversity of the three datasets, we derived the LD decay (Figure 1) resulting in the highest rates of association for the European maize landraces, followed by the commercial chicken dataset and the chicken diversity panel. The overall genetic diversity in all used datasets should be far smaller than in an outbred human population, which is the data structure BEAGLE was originally developed for. It should be noted that this comparison does not account for possible differences in ascertainment bias (Albrechtsen *et al*. 2010) between the arrays or the genetic diversity of species and their genomes. Since BEAGLE (and other HMM based imputation methods) are relying on local associations between markers this should still be a good indication for potential imputation performance.

**Figure 1.**
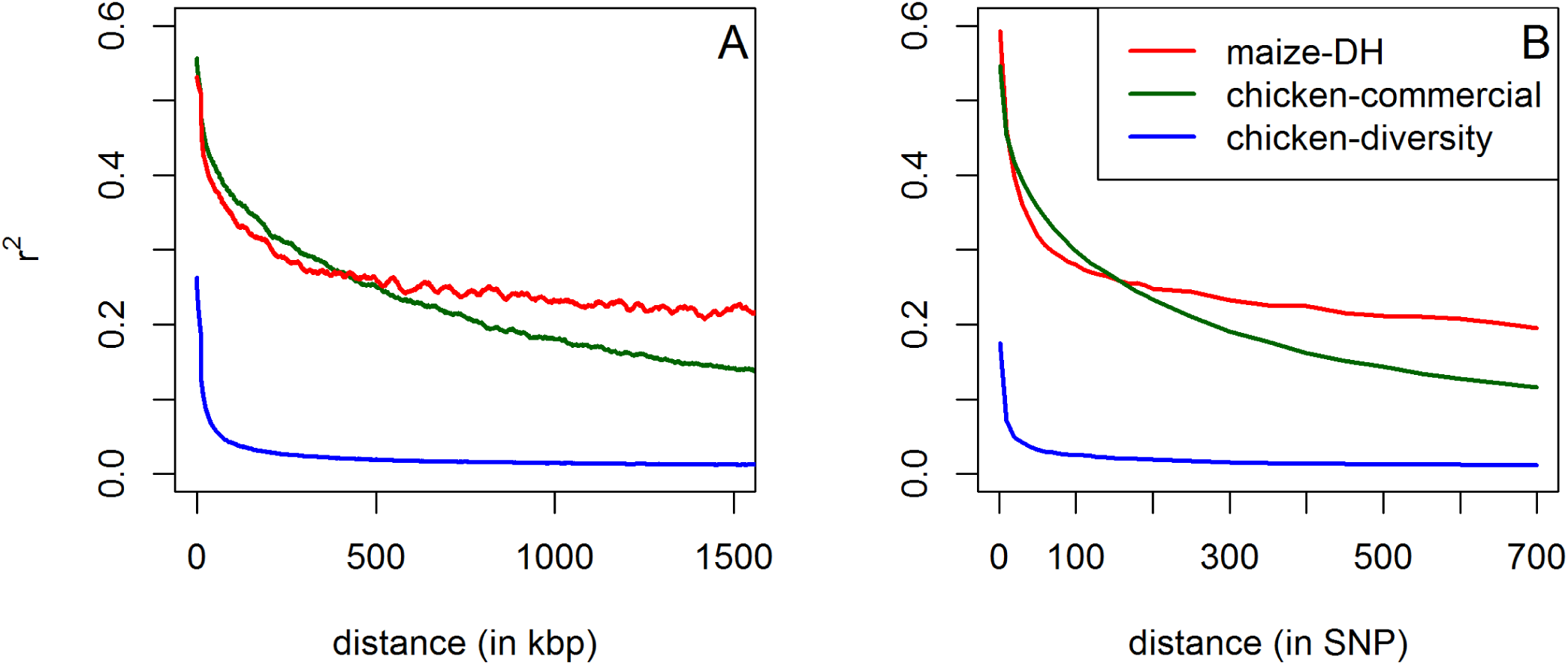
LD decay based on physical length (A) and marker distance (B) for chromosome 1 for all considered datasets. Outliers in (A) are corrected for by using a Nadaraya-Watson-estimator (Nadaraya 1964), using a Gaussian kernel and a bandwidth of 50 kb. (B) is using averaged values for each SNP distance.

### Evaluation Pipeline

The imputation process itself can be split up into three internally linked steps which can be of different importance based on the data at hand and, in the following, will be analyzed separately:

1. Inference: All partly or fully missing individual genotypes in the actual dataset are completed, but no additional markers are added.
2. Imputation of ungenotyped markers (UM imputation): Additional markers are added to the genetic data based on information provided by a second dataset (reference panel) with higher marker density.
3. Phasing: The two haplotypes of diploid individuals, i.e. their gametic phases, are estimated from genotype data.

To assess the quality of inference and UM imputation we used the following testing pipeline and repeated the procedure 100 times for each test. We start from a completed dataset in which missing genotypes have been imputed, and consider this as the “true” genotype dataset:

1. Randomly generate missing values (NAs) in the “true” genotype dataset.

- In case of inference set randomly chosen alleles of all genotypes to NA (in our case: 1% of all alleles with no partly missing genotypes).
- In case of UM imputation additionally set all entries in a particular marker to NA (maize: according to existing low density array (Ganal *et al*. 2011); chicken: 90% of all markers).
2. Perform the imputation procedure under a given parameter setting, software and potential use of a reference panel.
3. Evaluation of performance by comparison to the “true” dataset For more on this we refer to the following subsections.

### Evaluation of inference and UM imputation quality

To evaluate the quality of inference and UM imputation we count the total number of entries in the genotype matrix that are different to the “true” dataset (allelic error rate). In this procedure, markers with a low minor allele frequency have a lower impact on the overall quality than in the commonly used practice of calculating the correlation between imputed and “true” dataset (Hickey *et al.* 2012). To account for this, we will provide error rates depending on the allele frequency as well. A disadvantage of using a correlation is that it does not account for fixed markers as correlation is not defined for those markers, leading to them being excluded from the analysis. As rare variants tend to be more difficult to impute and those variants tend to be fixed at a higher rate, this leads to lower average correlations for methods imputing a rare allele (instead of just imputing the same variant everywhere). Therefore, a fair comparison should only consider those markers that are not fixed over all settings and different software. Especially for UM imputation this would lead to a much smaller set of markers to be considered.

### Evaluation of phasing quality

To evaluate phasing quality we use the switch error rate as defined in Lin *et al*. (2002), which evaluates the number of switches between neighboring heterozygous sites to recover the true haplotype phase compared to the total number of heterozygous markers. Since the true haplotype phase is usually not known the assessment of phasing quality is usually not as straight forward. As we are working with doubled haploid lines in the maize dataset, the true gametic phase is known and a “true” dataset for testing was generated by randomly combining two doubled haploid lines to a Pseudo *S*_0_. The rest of the pipeline can be performed in the same way as for the inference testing. For this analysis, we considered datasets with no missing genotypes to remove any potential noise caused by inference errors.

### Choice of reference panel in UM imputation

A common first question when planning to generate genetic data is how many individuals need to be genotyped with high marker density to obtain sufficient imputation quality for individuals genotyped with lower marker density. To evaluate this, we performed imputation on datasets containing 50 individuals as the “true” dataset in our pipeline and generated reference panels containing 25, 50, 100, 150, 200, 250, 300, and 350 individuals, respectively. Furthermore, we investigate how to chose the individuals to include in a reference panel. This is especially relevant when potential candidates for the reference panel vary in their relationship to the dataset itself. For this, we split the chicken diversity panel into ten subpopulations by iteratively minimizing the total sum of squared genetic distances between breeds within the subpopulations. Distances between the breeds were calculated as Nei standard genetic distances (Nei 1972). In a first step, the custom made algorithm randomly assigned the breeds to ten equal sized subpopulations. The contribution of each breed to the sum of squared distances was calculated and the algorithm started iteratively exchanging the most noisy breeds to other subpopulations. If there was a reduction of the total sum of squared distances within the subpopulations, the exchange was accepted and the contributions were calculated again. The process was repeated until no exchange could improve the fit. To overcome results depending on specific starting positions, the process was repeated for 60 random starting points. Nei standard genetic distances for evaluation of UM imputation quality of BEAGLE were calculated based on the subpopulation assignment of individuals and UM imputation was performed using the following reference panels:

(A) All other individuals of the same subpopulation
(B) All individuals of one other subpopulation
(C) All individuals of all other subpopulations
(D) All individuals of subpopulations with below-average Nei standard genetic distance to the dataset
(E) All individuals of those subpopulations with reduced error rates when testing A + B compared to A as the reference panel

Additionally combinations of panels A + B, A + C, A + D and A + E were tested. Tests were repeated 20 times for each subpopulation with datasets containing 50 randomly sampled individuals. For each dataset, all different reference panels were tested. The interested reader is referred to Supplementary Table S8 for a detailed list of the used subpopulation assignments and Supplementary Figure S1 for the resulting neighbor-joining-tree.

### Data Availability

Genetic data for chromosome 1 for all three panels used are available at https://github.com/tpook92/HaploBlocker. Table S1 and S2 contains error rates of UM imputation for the commercial breeding line and the diversity panel in chicken. Table S3 provides phasing error rates for the set of Pseudo *S_0_* with no missing data. Table S4 contains inference error rates for the PE DH-lines using different reference genomes. Table S5 and S6 contain lists of “critical” markers for KE and PE. Table S7 gives error rates of UM imputation using different reference panels for the subpopulations. Table S8 contains the subpopulation assignments for all chicken from the diversity panel. Table S9 contains the minimal error rates and used parameter settings for all performed tests.

Figure S1 provides the neighbor-joining-tree for the ten subpopulations of the chicken diversity panel. Figure S2 displays the relation between local LD and error rate for chromosome 9 in maize. Figure S3 displays the change in the number of errors in each marker by using low values of *buildwindow*. Figure S4 and S5 display the relation between DR2 and the number of errors per marker. Figure S6 - S24 display the relation between input parameters and error rates for inference in the maize data. Figure S25 - S45 display the relation between input parameters and error rates for inference and phasing for the set of Pseudo *S_0_*. Figure S46 - S74 display the relation between input parameters and error rates for UM imputation for the maize data, the commercial chicken line and the chicken diversity panel.

Supplemental files are available at FigShare:

## RESULTS

In the following, obtained error rates of the imputation under a variety of tuning option in BEAGLE are discussed. Here, we consider virtually all available parameters in BEAGLE, the size of the reference panel, and the underlying genetic map. The effect on the error rate of different tuning options are somewhat independent from each other as they commonly affect different parts of the imputation algorithm. Therefore, we will first consider each tuning option individually and later discuss suggested imputation pipelines for the different use cases.

Unless otherwise mentioned, we will report for maize the error rates in the landrace KE averaged over all chromosomes. Results for PE were similar with, on average, slightly increased error rates.

### Inference quality

On default, BEAGLE 5.0 (error rate: 0.0142%) and BEAGLE 5.1 (0.0127%) both clearly outperform BEAGLE 4.1 (0.255%) and BEAGLE 4.0 (0.201%) for the maize data. For all four versions the error rates are significantly higher for alleles with low frequency (Figure 2). In regard to the location of inference errors one can observe a high volatility with a tendency to have increased error rates in telomeric regions (Figure 3). Additionally, error rates in regions of high LD tend to be lower (Supplementary Figure S2).

**Figure 2.**
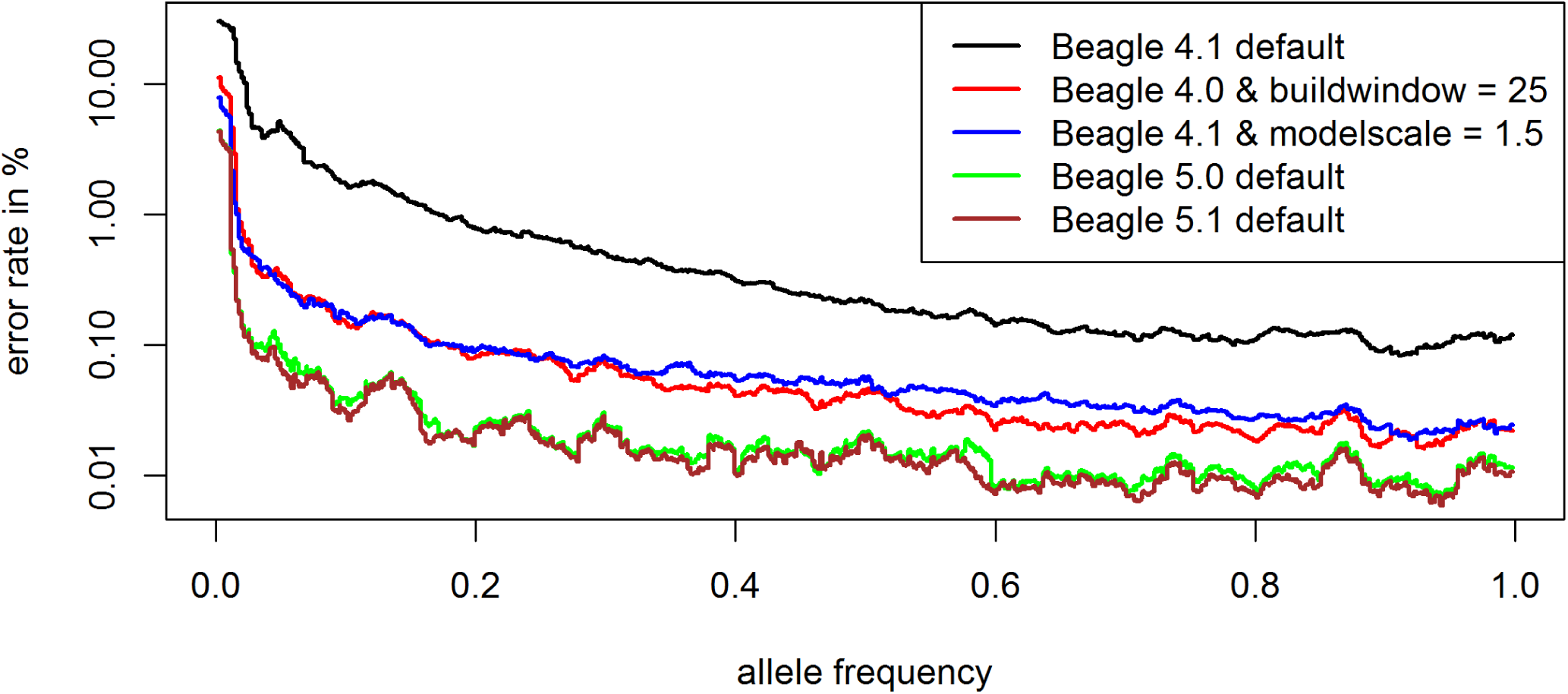
Allele specific error rate depending on the allele frequency under different BEAGLE settings for the maize data. Only those dataset entries with the respective allele in the “true” dataset are considered when deriving the allele specific error rate. Y-axis is log-scaled.

**Figure 3.**
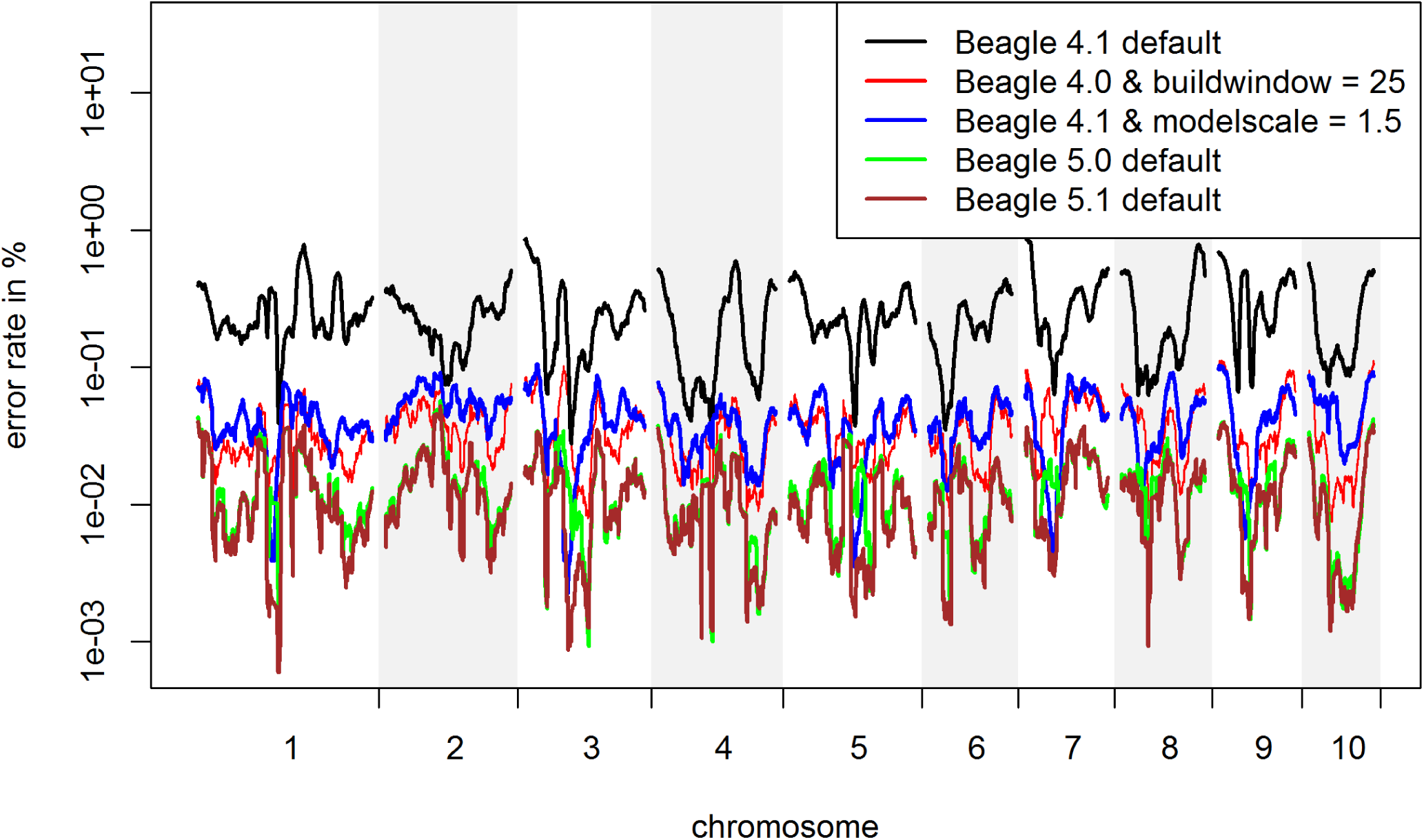
Inference error rate based on the location of the genome. Outliers are corrected for by using a Nadaraya-Watson-estimator (Nadaraya 1964), using a Gaussian kernel and a bandwidth of 3,000 markers for the maize data. Y-axis is log-scaled.

For all four versions the biggest improvement was obtained by tuning parameters that are affecting the structure of the haplotype cluster. The optimal parameter values (Table 1) for *buildwindow* (4.0), *singlescale* (4.0), *modelscale* (4.1) lead to less similar haplotypes being clustered jointly. *Phase-segment* (5.0), *phase-states* (5.0 / 5.1) affect the minimum length and number of different haplotypes in the haplotype cluster. Overall, all these settings lead to longer and/or less related haplotypes to be considered jointly. The gains by fitting those parameters are much higher in BEAGLE 4.0 and 4.1 but overall error rates are still higher than in BEAGLE 5.0 and 5.1 (Table 1) with BEAGLE 5.1 performing best. Improvements in overall inference quality can be observed for all allele frequency classes and regions in the genome (Figures 2 & 3). It should be noted that in contrast to later tests in UM imputation the use of low (and probably more realistic) values for *ne* (effective population size) can lead to substantially increased error rates (Figure 4). The interested reader is referred to Supplementary Figures S6 - S24 for the effect on the inference error rate for different parameters. For the maize data the inference error rates were basically unaffected by the number of iterations performed in any of the imputation steps in BEAGLE (Table 1). Since the haplotype phase in DH-lines is known and the main purpose of further iterations in BEAGLE is to improve that haplotype phase, this should not be that surprising. After parameter tuning error rates are still lowest in BEAGLE 5.1 with 0.0122% but differences are considerably reduced (BEAGLE 4.0: 0.0307%, BEAGLE 4.1: 0.0436%, BEAGLE 5.0: 0.0132%, Supplementary Table S9). Tuning of both *singlescale* and *buildwindow* in BEAGLE 4.0 jointly did not further improve performance with *buildwindow* overall performing better for inference. Even though error rates for extremely low values for *buildwindow* are lowest, this change is not recommended as some markers do show massively increased error rates (Supplementary Figure S3).

**Figure 4.**
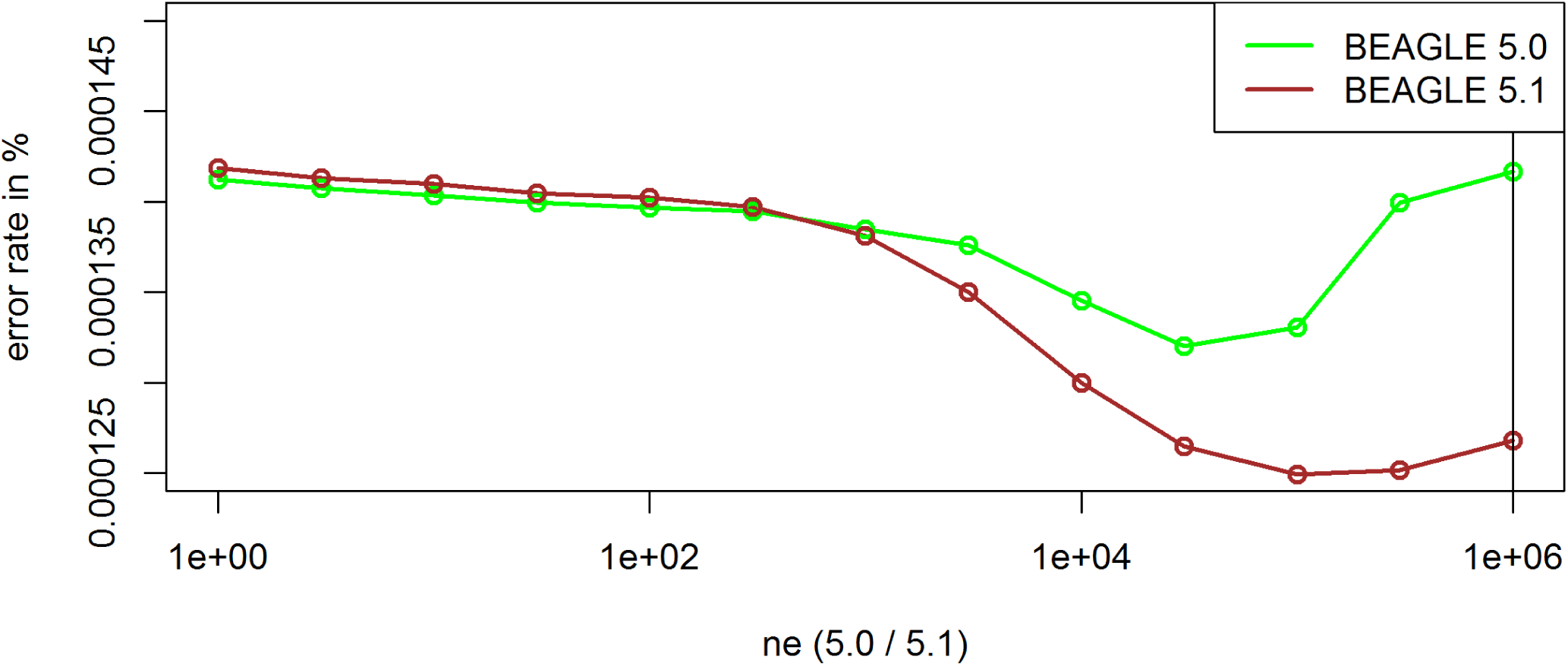
Effect of the parameter *ne* on the inference error rates for the maize data in BEAGLE 5.0 and 5.1. Default settings are indicated by the vertical line.

**Table 1.**
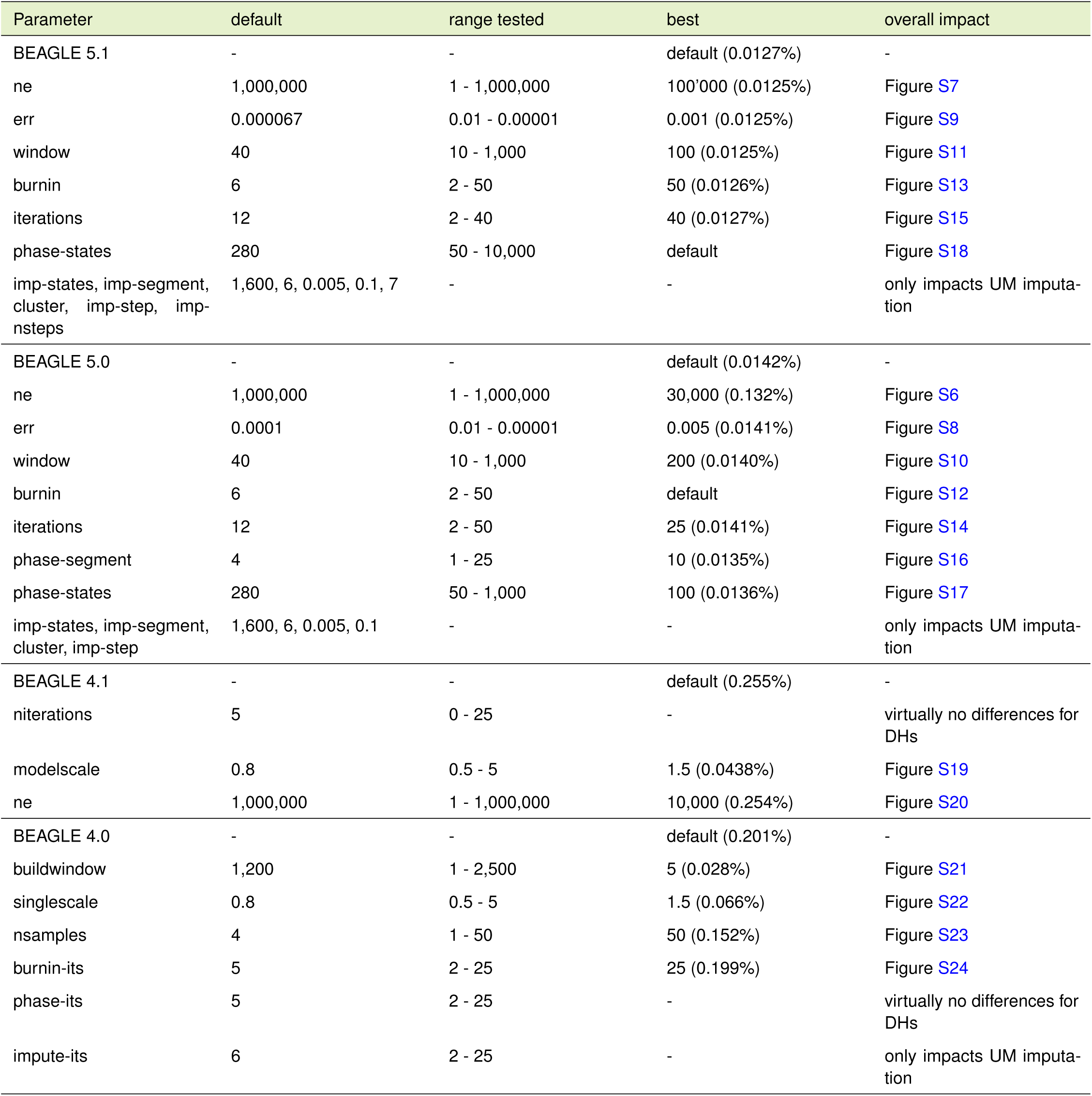
Inference error for the KE DH-lines by changing a single imputing parameter.

The inference error rates for the chicken diversity panel are much higher for all versions (∼1%) and the relative improvement obtained by adapting parameter settings is lower. As the chicken diversity panel contains more variation and is structurally more similar to outbred human data than the European landraces in maize, this should not be that surprising. With the exception of the parameter *err* the change from the default was always in the same direction as for the maize data. As *err* is controlling the allele mismatch probability of known alleles when identifying the most likely path through the haplotype cluster (Browning *et al*. 2018) this can be seen as an indicator for a higher overall data quality for the maize data. Lowest obtained error rates are 1.01% for BEAGLE 4, 0.80% for BEAGLE 4.1, 0.81% for BEAGLE 5.0 and 0.82% for BEAGLE 5.1 (Supplementary Table S9).

Inference error rates for the datasets from the commercial chicken breeding program are between 0.20% and 0.23% for basically all tested settings, leading us to conclude that for inference on this dataset there is not much potential to decrease error rates. A potential reason for this is that other error sources like SNP calling errors may be higher than inference error rates.

When working with the Pseudo *S*_0_ in maize instead, ideal parameter settings are very similar with the key difference of additional gains by increasing the number of iterations performed (Table 2). As the algorithm starts with randomly phased genotypes and improves the phase in each iteration, this should again not be surprising. However, excessive *burnin* iterations prior to the actual algorithm only worsened results. The interested reader is referred to Supplementary Figure S25 - S45 for parameter influences on both inference and phasing quality for the Pseudo *S*_0_. Inference accuracies after parameter tuning are again similar with BEAGLE 5.0 performing best (BEAGLE 4.0: 0.0193%, BEAGLE 4.1: 0.0168%, BEAGLE 5.0: 0.0109%, BEAGLE 5.1: 0.0148%). Note that error rates given in Table 2 are just for chromosome 10, as not all tests were performed in sufficient sample size for all chromosomes but effect of parameters results should be very similar for all chromosomes. For all three datasets containing heterozygous individuals BEAGLE 5.0 outperformed BEAGLE 5.1, with differences being highest for the set of Pseudo *S*_0_.

**Table 2.** Inference and phasing error for the 250 Pseudo S_0_ lines based on the KE DH-lines for chromosome 10. * BEAGLE crashed for this dataset when using *phase-segment* > 10, *phase-states* < 100 or *phase-states* > 10,000.

| Parameter | default | range tested | best inference | best phasing | overall impact |
| --- | --- | --- | --- | --- | --- |
| BEAGLE 5.1 | - | - | default (0.0239%) | default (2,540) | - |
| ne | 1,000,000 | 1 - 1,000,000 | 30 (0.0168%) | 10 (3,206) | Figure S26 |
| err | 0.00015 | 0.05 - 0.00001 | 0.0005 (0.0229%) | 0.05 (2,556) | Figure S28 |
| window | 40 | 10 - 1,000 | 200 (0.0179%) | 200 (2,581) | Figure S30 |
| burnin | 6 | 2 - 25 | 25 (0.0229%) | 2 (2,555) | Figure S32 |
| iterations | 12 | 2 - 40 | default | 40 (2,638) | Figure S34 |
| phase-states | 280 | 100* - 10,000* | 10,000 (0.0168%) | 5,000 (2,675) | Figure S37 |
| imp-states, imp-segment, cluster, imp-step, imp-nsteps | 1,600, 6, 0.005, 0.1, 7 | - | - | - | only impacts UM imputation |
| BEAGLE 5.0 | - | - | default (0.0138%) | default (2,716) | - |
| ne | 1,000,000 | 1 - 1,000,000 | 1 (0.0111%) | 30,000 (3,136) | Figure S25 |
| err | 0.0001 | 0.05 - 0.00001 | 0.005 (0.133%) | 0.001 (2,747) | Figure S27 |
| window | 40 | 10 - 200 | 100 (0.0139%) | 200 (2,737) | Figure S29 |
| burnin | 6 | 2 - 25 | 2 (0.0136%) | 2 (2,748) | Figure S31 |
| iterations | 12 | 2 - 40 | 20 (0.0135%) | 40 (2,760) | Figure S33 |
| phase-segment | 4 | 1 - 10* | 10 (0.132%) | 10 (2,758) | Figure S35 |
| phase-states | 280 | 100* - 10,000* | 10,000 (0.0130%) | 5,000 (2,815) | Figure S36 |
| imp-states, imp-segment, cluster | 1,600, 6, 0.005 | - | - | - | only impacts UM imputation |
| BEAGLE 4.1 | - | - | default (0.0345%) | default (2,617) | - |
| niterations | 5 | 0 - 25 | 25 (0.0249%) | 25 (3,392) | Figure S38 |
| modelscale | 0.8 | 0.5 - 5 | 1 (0.0198%) | 1 (3,223) | Figure S39 |
| ne | 1,000,000 | 1 - 1,000,000 | 30 (0.0325%) | 30,000 (2,999) | Figure S40 |
| BEAGLE 4.0 | - | - | default (0.119%) | default (1,240) | - |
| buildwindow | 1,200 | 1 - 5,000 | 50 (0.0316%) | 5,000 (1,618) | Figure S41 |
| singlescale | 0.8 | 0.5 - 5 | 1.0 (0.0626%) | 1.25 (1,955) | Figure S42 |
| nsamples | 4 | 1 - 50 | 50 (0.0780%) | 50 (2,308) | Figure S43 |
| burnin-its | 5 | 2 - 50 | 50 (0.108%) | 50 (1,599) | Figure S44 |
| phase-its | 5 | 2 - 50 | 50 (0.0944%) | 50 (2,320) | Figure S45 |
| impute-its | 5 | 2 - 50 | - | - | only impacts UM imputation |

### Phasing quality

The number of phasing errors for the set of Pseudo *S*_0_ in maize is extremely low with just one phasing error per 2,540 heterozygous markers in BEAGLE 5.1, which should be sufficient for most applications, and the obtainable improvements by parameter tuning were relatively low (Table 2). Error rates in BEAGLE 5.0 were about 10% lower (2’716). Biggest improvements in both BEAGLE 5.0 and 5.1 were obtained by adaptation of *ne*. For *S*_0_ the ideal parametrization in BEAGLE 5.0 for phasing is much higher than for inference (Figure S25). Especially for BEAGLE 4.0 and 4.1 parameters influencing the structure of the haplotype library had substantial impact on the error rates. In contrast to inference and UM imputation the ideal parametrization for *buildwindow* (4.0) and *phase-states* (5.0 / 5.1) are higher than the default settings (Table S3). This in turn leads to only highly related haplotypes to be considered jointly.

To further isolate the structure of phasing errors the same tests were performed on a set of Pseudo *S*_0_ without missing alleles. The interested reader is referred to the Supplementary Table S3 for detailed results on this. As phasing is not affected by potential inference errors in this case, error rates are even lower (BEAGLE 5.1 default: one error per 5’756 heterozygous markers, BEAGLE 5.0: 6,141) but the direction of improvement for all parameters stays the same. It should be noted that the maize dataset considered in this study contains highly related individuals and a substantial ascertainment bias towards markers with medium allele frequency (Albrechtsen *et al*. 2010) which both should improve phasing accuracy. For datasets containing less related individuals and sequence data, phasing accuracies can be substantially worse.

### UM Imputation quality

The algorithm used for UM imputation in BEAGLE 5.0 and 5.1 is the same, thereby differences are only caused because of slightly different techniques for phasing (B. Browning, personal communication). As no phasing is required for the DH-lines error rates never differed by more than 0.001% and are here reported jointly. When performing UM imputation, error rates were much higher than in the inference case. For all considered datasets tuning of *ne* was absolutely essential (Table 3, Figure 5), because individuals in the considered livestock and crop datasets are far more related than in an outbred human population with an effective population size of 1,000,000 that is assumed in BEAGLE as default. In the imputation algorithm a low value for *ne* is leading to a reduced probability to switch to a random node in the haplotype cluster and should therefore be beneficial for highly related individuals (Browning and Browning 2016; Browning *et al*. 2018). BEAGLE 4.0 does not provide a parameter for the effective population size and is just assuming equidistant markers and fixed switch rates.

**Figure 5.**
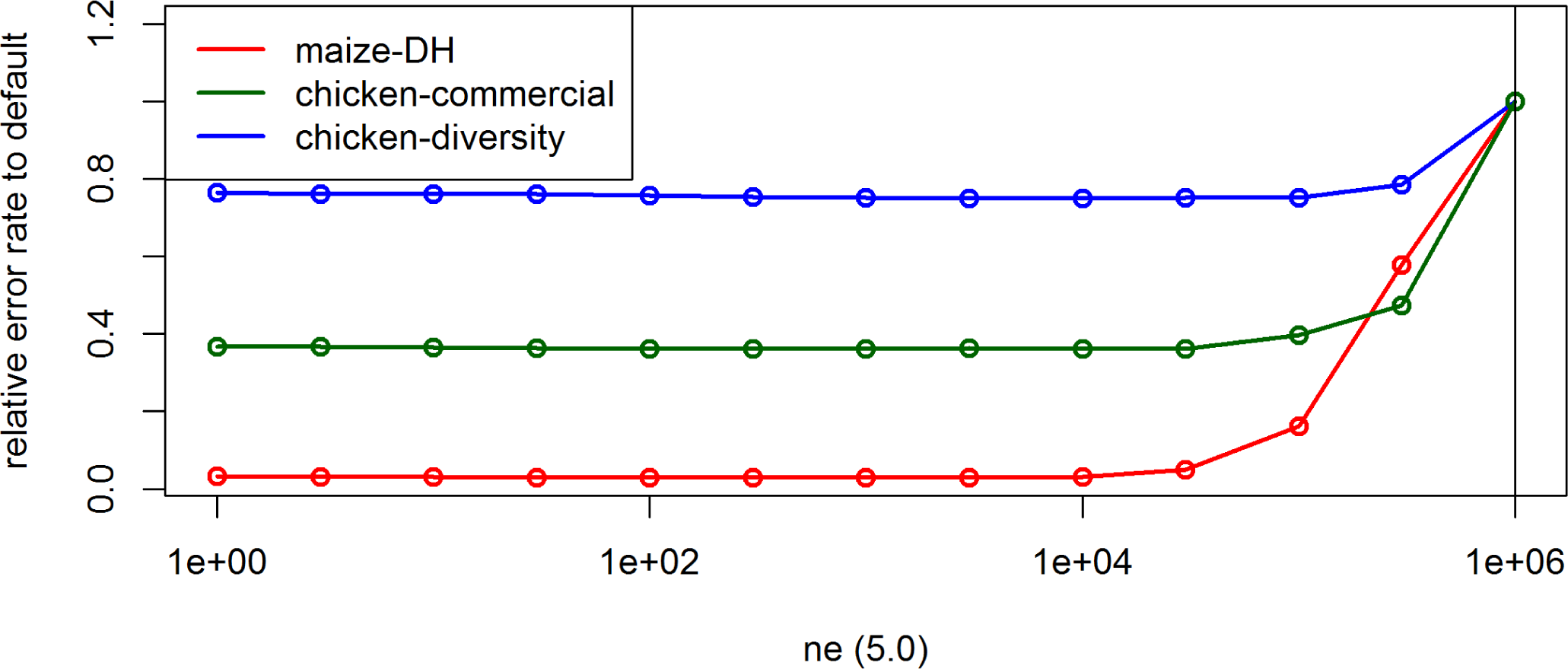
Effect of the parameter *ne* on the UM imputation error rate for the maize data, the commercial chicken line and the chicken diversity panel in BEAGLE 5.0. Default settings are indicated by the vertical line.

**Table 3.**
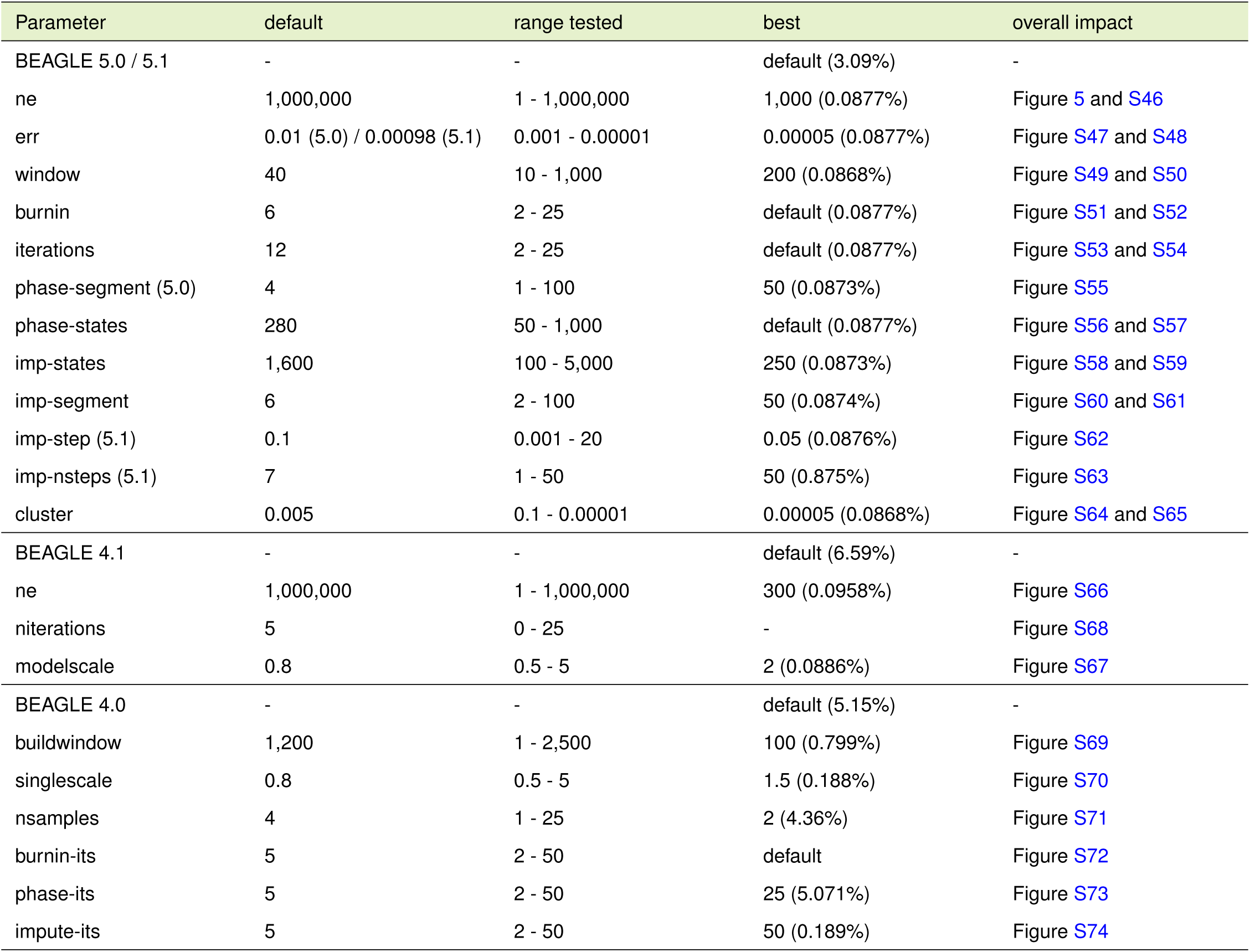
UM imputation error for the KE DH-lines by changing a single imputing parameter with *ne* = 1,000 for BEAGLE 5.0 / 5.1 and *ne* = 300 for BEAGLE 4.1.

| Parameter | default | range tested | best | overall impact |
| --- | --- | --- | --- | --- |
| BEAGLE 5.0 / 5.1 | - | - | default (3.09%) | - |
| ne | 1,000,000 | 1 - 1,000,000 | 1,000 (0.0877%) | Figure 5 and S46 |
| err | 0.01 (5.0) / 0.00098 (5.1) | 0.001 - 0.00001 | 0.00005 (0.0877%) | Figure S47 and S48 |
| window | 40 | 10 - 1,000 | 200 (0.0868%) | Figure S49 and S50 |
| burnin | 6 | 2 - 25 | default (0.0877%) | Figure S51 and S52 |
| iterations | 12 | 2 - 25 | default (0.0877%) | Figure S53 and S54 |
| phase-segment (5.0) | 4 | 1 - 100 | 50 (0.0873%) | Figure S55 |
| phase-states | 280 | 50 - 1,000 | default (0.0877%) | Figure S56 and S57 |
| imp-states | 1,600 | 100 - 5,000 | 250 (0.0873%) | Figure S58 and S59 |
| imp-segment | 6 | 2 - 100 | 50 (0.0874%) | Figure S60 and S61 |
| imp-step (5.1) | 0.1 | 0.001 - 20 | 0.05 (0.0876%) | Figure S62 |
| imp-nsteps (5.1) | 7 | 1 - 50 | 50 (0.875%) | Figure S63 |
| cluster | 0.005 | 0.1 - 0.00001 | 0.00005 (0.0868%) | Figure S64 and S65 |
| BEAGLE 4.1 | - | - | default (6.59%) | - |
| ne | 1,000,000 | 1 - 1,000,000 | 300 (0.0958%) | Figure S66 |
| niterations | 5 | 0 - 25 | - | Figure S68 |
| modelscale | 0.8 | 0.5 - 5 | 2 (0.0886%) | Figure S67 |
| BEAGLE 4.0 | - | - | default (5.15%) | - |
| buildwindow | 1,200 | 1 - 2,500 | 100 (0.799%) | Figure S69 |
| singlescale | 0.8 | 0.5 - 5 | 1.5 (0.188%) | Figure S70 |
| nsamples | 4 | 1 - 25 | 2 (4.36%) | Figure S71 |
| burnin-its | 5 | 2 - 50 | default | Figure S72 |
| phase-its | 5 | 2 - 50 | 25 (5.071%) | Figure S73 |
| impute-its | 5 | 2 - 50 | 50 (0.189%) | Figure S74 |

All other parameter settings were tested with adapted *ne*, as relative effects were virtually zero otherwise. Appropriate parameter settings for the other parameters were similar to the inference case (Table 3) but the overall deviations from the default for *buildwindow, singlescale* and *modelscale* were slightly lower. As the number of informative markers in a window with a set number of markers is lower than in the inference case this also makes sense from a modeling perspective. In BEAGLE 5.0 and 5.1 there are additional parameters to control the structure of the haplotype cluster that are only available for UM imputation (*imp-segment, imp-states, cluster*). Similar to inference, the optimized parameter settings lead to longer and less related haplotypes to be considered jointly. Furthermore, a method to detect identity-by-state (IBS) segments (*imp-step, imp-nsteps*) has been added in BEAGLE 5.1 but defaults are already adequately chosen for the maize data. After parameter adaptation error rates in BEAGLE 5.0 and 5.1 were lowest (0.0856% / 0.0857%), followed by BEAGLE 4.1 (0.0887%) and BEAGLE 4.0 (0.139%) (Supplementary Table S9).

For both chicken datasets similar results were obtained with BEAGLE 5.0 slightly outperforming BEAGLE 5.1 in for these sets. The interested reader is referred to Supplementary Table S1 and S2 for detailed results for UM imputation for the chicken panels. Overall, the relative gains by adaptation of *ne* for both the commercial breeding line (0.774% to 0.280% in BEAGLE 5.0) and the diversity chicken panel (3.313% to 2.484% in BEAGLE 5.0) were lower than for the maize data. The optimal parametrization for the effective population for the diversity panel was highest (*ne* = 3,000). With this, the smaller gains by tuning the effective population size nicely support our expectation of the effective population sizes of the underlying populations. It should still be noted that especially BEAGLE 5.0 and 5.1 were very robust to changes in the effective population size (Figure 5 and S46) and overall error rates differ by only 0.013% for an effective population size between *ne* = 1 and *ne* = 10,000 for the maize dataset, indicating that the use of any reasonable value should work here. As the default of 1,000,000 is not realistic for most livestock and crop datasets, adaptation is necessary and critical when performing UM imputation. For BEA442 4.1 there were usually no statistically significant differences between reasonable ne values and overall variance in error rates between runs was slightly higher.

As one would expect a larger reference panel leads to smaller error rates for UM imputation (Figure 6). Overall, the effect of a larger reference panel in BEAGLE 5.0 was higher than for BEAGLE 4.1. It should still be noted that even for a reference panel with 20 individuals error rates after parameter tuning were below 1% for the maize data and overall error rates only reduce slightly after reaching a size of 150. With higher amounts of overall genetic diversity, the required size of the reference panel should be increasing (Zhang *et al*. 2013).

**Figure 6.**
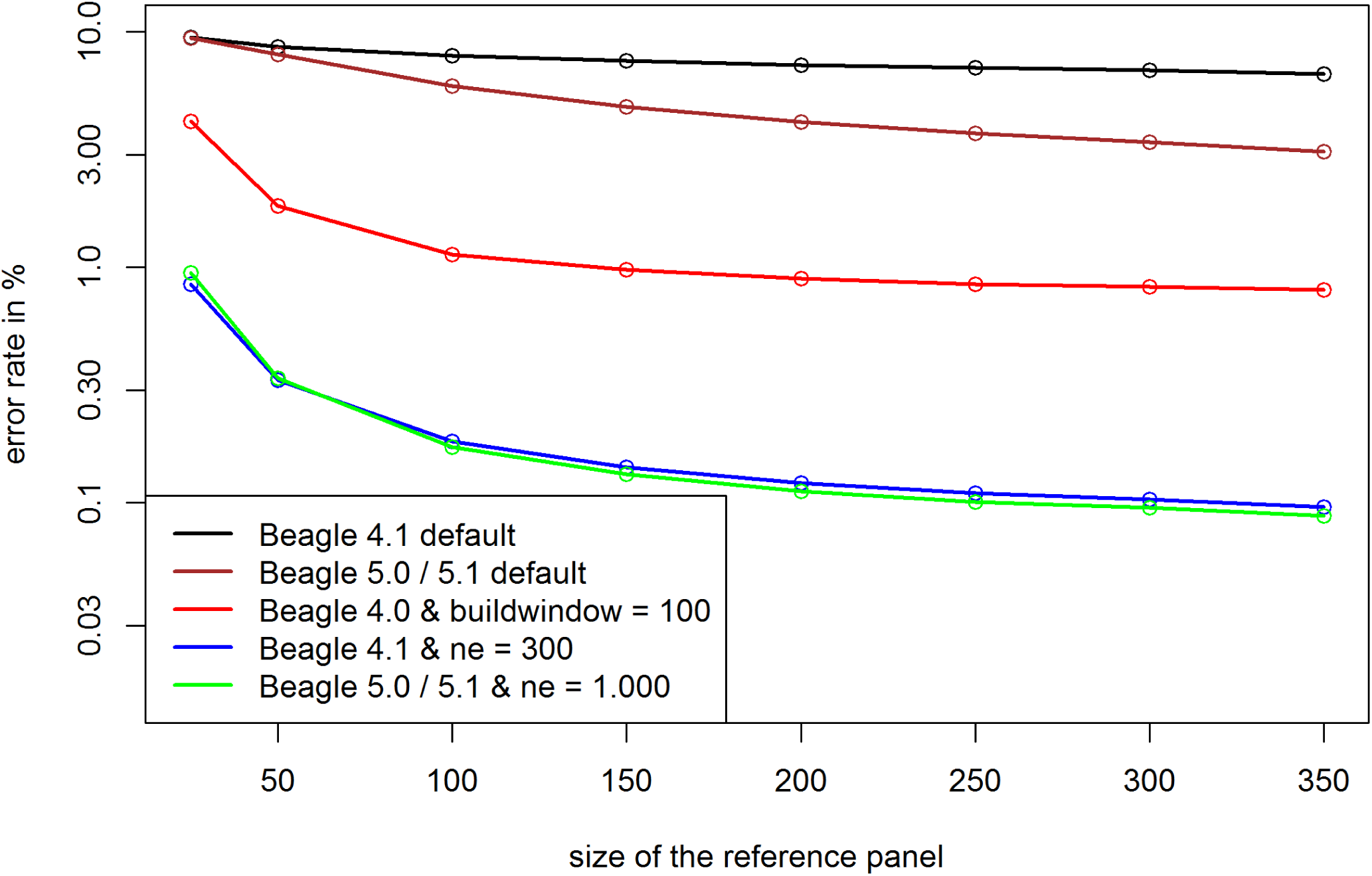
Error rates for UM imputation depending on the size of the reference panel in the maize data. Y-axis is log-scaled.

### Comparison of reference genomes

The most commonly used reference genome in maize genetics is the dent line B73 (Schnable *et al*. 2009; Jiao *et al*. 2017). The European maize landraces tested here are considered as flint germplasm with potential major differences in the physical map (Unterseer *et al*. 2016). After reducing error rates of inference by choosing appropriate parameter settings, markers with high error rates tend to be clustered (Figure 7). Markers and regions with high inference error rate can be considered as candidates for misalignment in the genetic map. We compared our results obtained with B73v4 (Jiao *et al*. 2017) to those obtained with reference genomes of the flint lines F7, EP1, DK105 and PE0075 (Unterseer *et al*. 2017). Since the array itself was constructed using B73 as a reference (Unterseer *et al*. 2014) more markers can be mapped to the B73 reference than to the other reference genomes. For those markers mapped to both B73 and the respective flint reference genomes average error rates for inference are reduced by 3-5% (Table 4). This improvement is mainly caused by a much reduced number of markers with extremely high error rates. On average, the overall number of markers with error rates above 10% (here referred to as: “critical” markers) is reduced by 57%. For a detailed list of the “critical” markers for all reference genomes mapped on the 600k array (Unterseer *et al*. 2014), we refer to Supplementary Table S5 and S6. No notable difference in inference quality for PE when using PE0075 as the reference genome compared to other flint references (Supplementary Table S4) was found.

**Figure 7.**
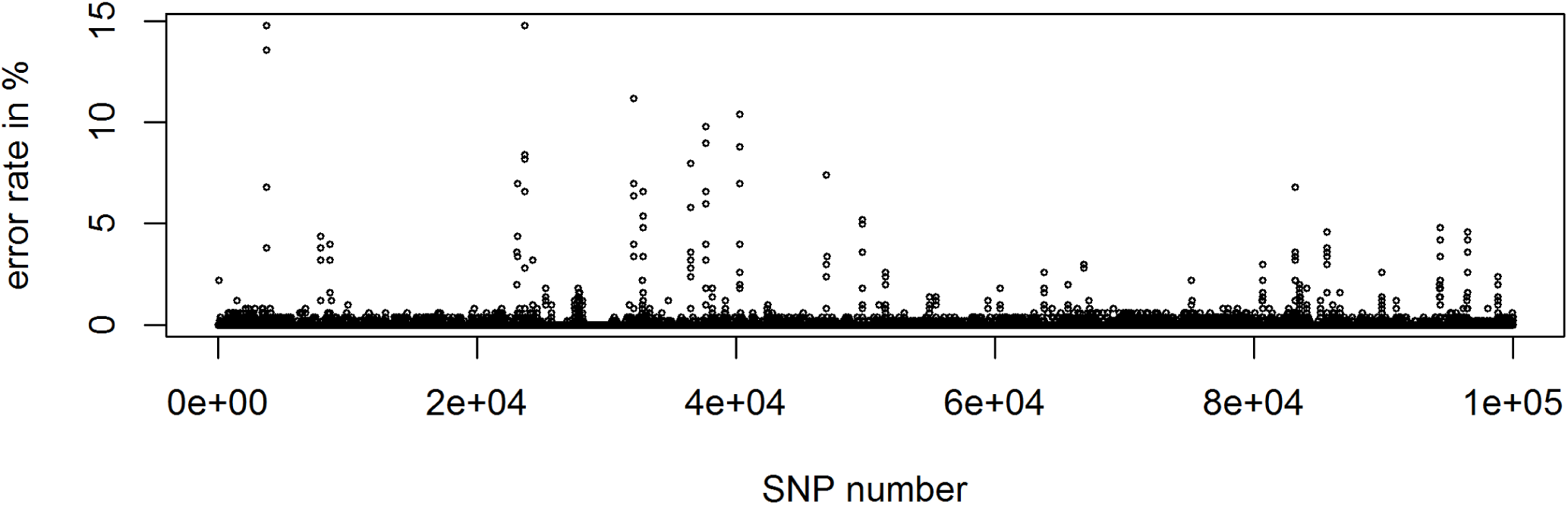
Error rate per marker for the first 100,000 SNPs according to physical position (starting with chromosome 1) using BEAGLE 5.0

**Table 4.** Inference error rates using different reference genomes compared to B73 for KE DH-lines. Only markers mapped on both the flint reference genome & B73v4 (Jiao et al. 2017) are considered for “critical” markers (error rate > 10%).

| Reference genome | F7 | EP1 | DK105 | PE0075 |
| --- | --- | --- | --- | --- |
| Overlapping markers to B73v4 | 352,326 | 342,037 | 338,882 | 338,244 |
| "Critical" markers when using this map | 109 | 113 | 115 | 114 |
| "Critical" markers when using B73v4 | 271 | 264 | 262 | 262 |
| Relative change in error rate | - 5.11% | -3.87% | -4.68% | -3.32% |

### Use of a genetic map

Up to BEAGLE 4.0 all markers are assumed to be equidistant, whereas in BEAGLE 4.1, 5.0 and 5.1 the genetic distance between markers can be provided. On default, the position in base pairs is converted by a ration of 100,000,000 base pairs per Morgan. This might be realistic for human genetics but for chicken a ratio of 41,203,130 / 33,955,860 / 26,631,160 base pairs per Morgan for chromosomes 1 / 7 / 20 is more realistic (Groenen *et al*. 2009). However, the use of those genetic maps without any further parameter adaptation leads to massively increased error rates. Error rates for UM imputation increased to 3.23% for the commercial line and 15.8% for the diversity panel compared to the 0.774% and 3.313% without a provided genetic map in BEAGLE 5.0. A potential reason for this is that other parameters like *ne* and *imp-segment* are implicitly affected by the higher distance between markers, leading to smaller segments being considered jointly in the haplotype cluster. After additional fitting of *ne* error rates reduced to values (0.276% / 2.50%) which were very similar to those obtained without use of a genetic map (0.280% / 2.48%; Supplementary Table S1 and S2).

### Quality control using Dosage R-Squared

When performing UM imputation BEAGLE is providing the measurement Dosage R-Squared (DR2; (Browning and Browning 2009)) as an estimate for the uncertainty for the imputation quality in each respective marker. When using BEAGLE 5.0 with adapted *ne*, only some markers have low DR2 values and the observed error rates in those markers are highly increased (Figure 8.A). Markers with DR2 values below 0.8 on average had 140 times as many imputing errors for UM imputation. Note that no scaling for the allele frequency was performed here and no apparent correlation between DR2 values and minor allele frequencies could be observed. In case of no adaption of the effective population size, the number of markers with low DR2 values is massively increased. Even though error rates are still a higher for markers with low DR2, the relative differences are much lower (18 times as many errors for markers with DR2 < 0.8). Even more problematic for filtering is that in contrast to the 44 problematic markers after parameter adaptation, a total of 31,635 of the 62,986 markers in the panel have DR2 values below 0.8 (Figure 8.B). Results for the commercial chicken line (Supplementary Figure S4) and the diversity panel (Figure Supplementary Figure S5) are similar even though differences in DR2 are not as distinct for adapted parameter settings.

**Figure 8.**
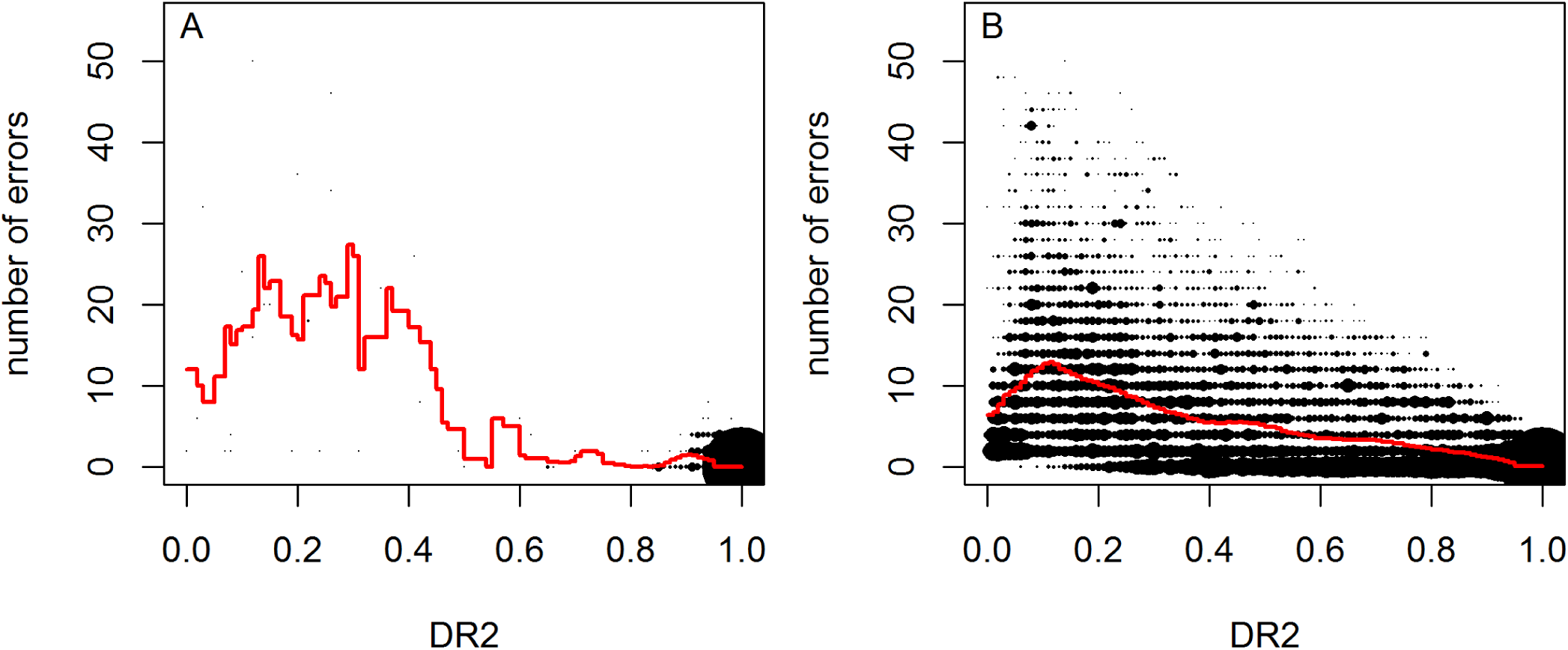
DR2 values in relation to the obtained number of error per marker after fitting of *ne* (A) and on default (B) in BEAGLE 5.0 for the maize data. 50 / 350 DH-lines were used for study / reference sample.

### Choice of the reference panel

In case the reference population has a lot of stratification, the design of a good reference panel for UM imputation is more difficult, as genetically distant individuals may introduce more noise than relevant information to the model. When comparing results for all considered reference datasets for UM imputation of a single subpopulation it becomes apparent that UM imputation without other individuals from the same subpopulation leads to extremely high error rates (>15%) and thus should in practice only be performed with extreme caution. In contrast, the decision to include other subpopulations in the reference panel is not as clear. When including single other subpopulations in the reference panel we observe significant effects on the overall error rate of UM imputation. Absolute differences of UM imputation error rates are between −0.307% and +0.604% with overall error rates between 1% and 4%. For a detailed list containing all changes in error rates when including a single other subpopulation in the reference panel, we refer to Supplementary Table S7. It should be noted that subpopulations with lower genetic distance to the dataset tend to reduce the error rate and a less related subpopulation leads to an increased error rate (Figure 9). For all ten subpopulations the slope of the error rate in regard to distance to the subgroup is statistically significantly positive with the main difference between the subpopulations being the intercept. The most extreme case for this is subpopulation 6 (turquoise Δ in Figure 9; including all wild types). For this group the inclusion of any other subpopulation in the reference panel decreases the imputation quality and is ignored for all averages and statistics in this subsection. Even though SNP-based genetic distances to other subgroups are relatively low, the time to the last common ancestor of any other subpopulation is most likely relatively high. Overall imputation quality when using a reference panel containing all subpopulations is worse than when using a reference panel with only those subpopulation with below average genetic distance (Nei 1972) to the dataset (2.25% vs. 2.18% - Figure 10).

**Figure 9.**
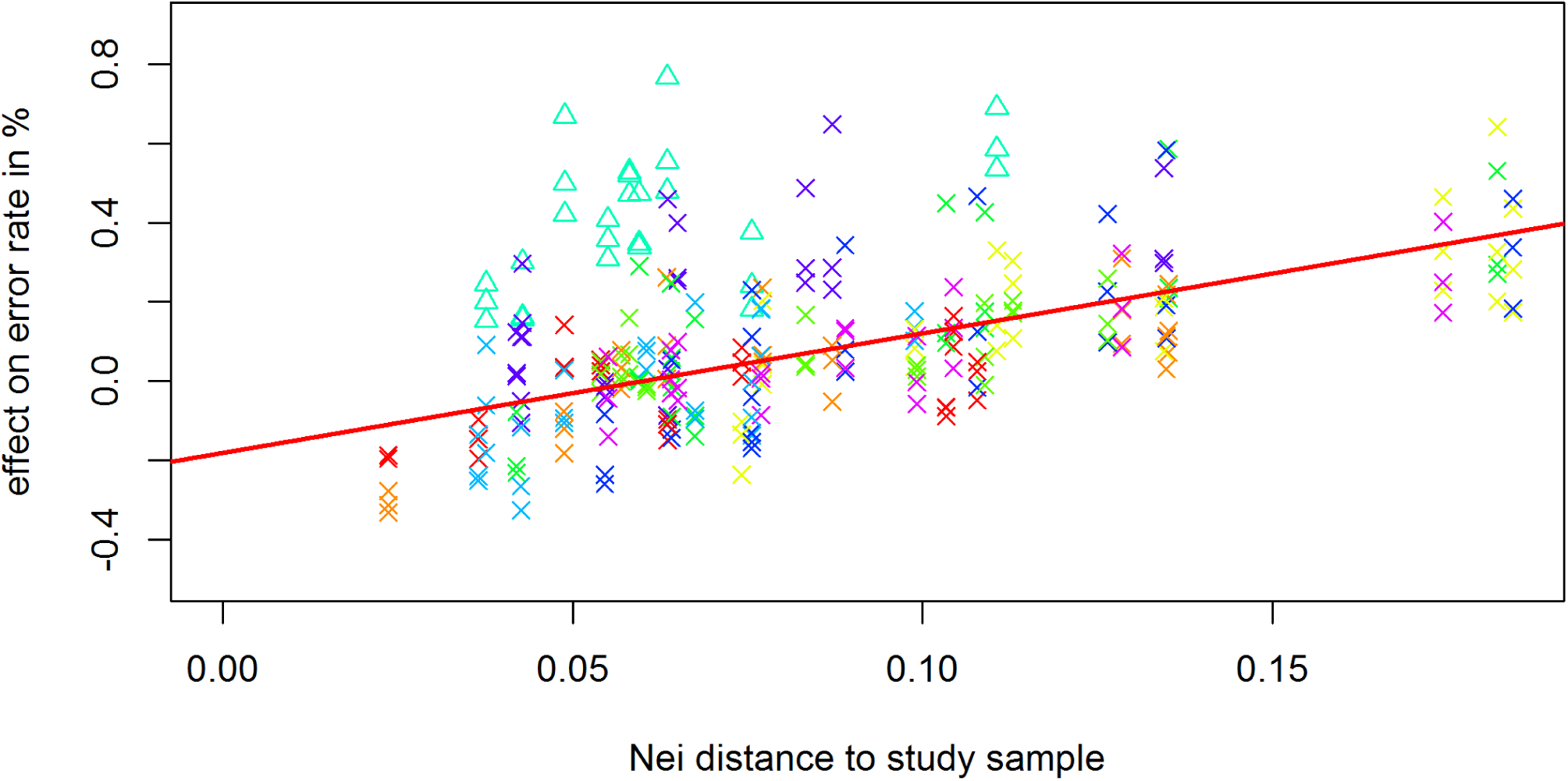
Effect of the inclusion of a single subpopulation in the reference panel based on their genetic distance to the dataset for the chicken diversity panel. Colors according to the subpopulation used as the real dataset in Supplementary Figure S1. For a detailed list of subpopulation assignment we refer to Supplementary Table S8. Subpopulation 6 (including wild types - turquoise Δ) is ignored in the regression.

**Figure 10.**
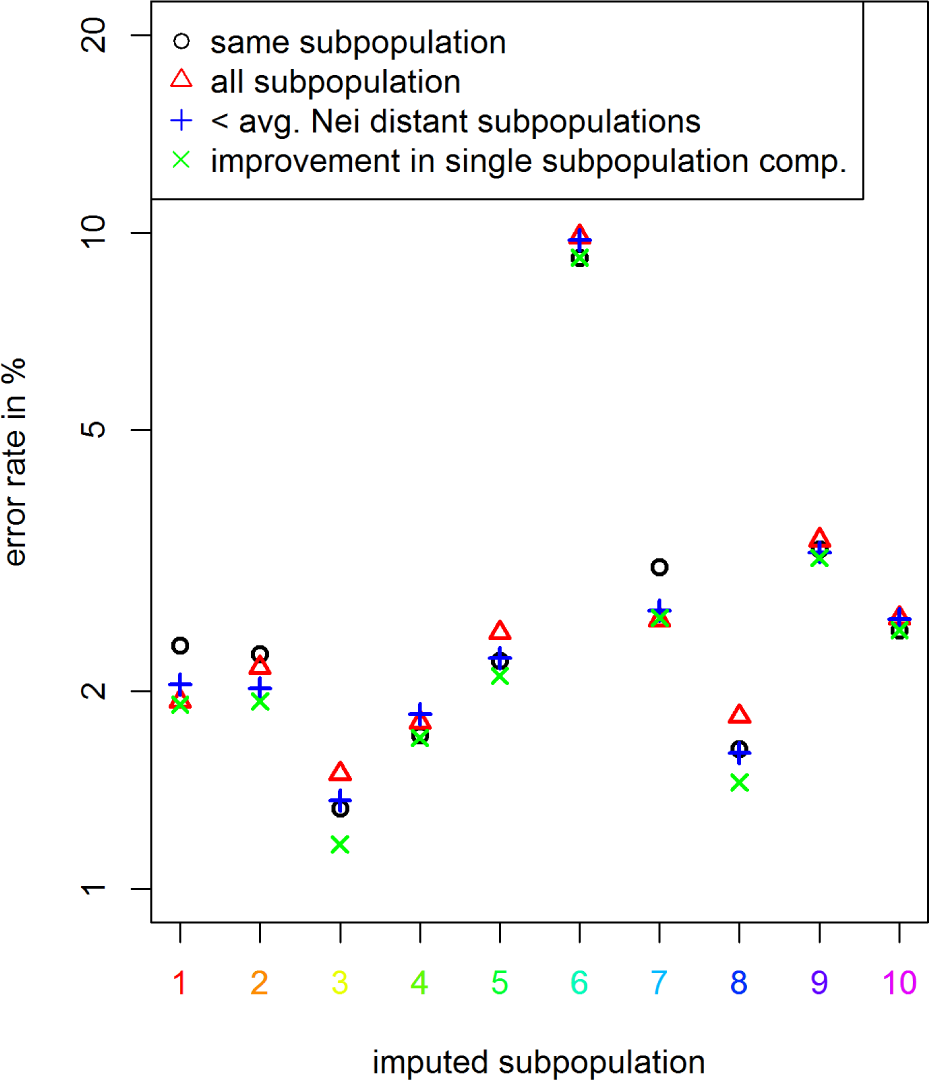
Comparison of error rates of UM imputation for different reference panels for the different subpopulations in the chicken diversity panel. Y-axis is log-scaled. For a detailed list on which individual is assigned to which subpopulation we refer to Supplementary Table S8.

Even though results are statistically significant (two-sample t-test: p-value: 0.0117), differences are only minor and of limited practical relevance for most applications. In our analysis a reference panel containing only the individuals of the same subpopulation on average lead to an UM imputation error of 2.26% with no statistically significant difference to reference panels containing all subpopulations. When performing in-depth analysis for which regions of the dataset UM imputation quality is improved, we observed that especially those individuals with rare variants and overall higher error rates benefited from including more samples in the reference. On the contrary, already well imputed individuals usually had similar or slightly increased error rates. When using a reference panel containing all those subpopulations that individually lead to reduced error rates, average error rates are reduced to 2.06%. It should be noted that a selection based on error rates in UM imputation is usually not possible in practice. Nevertheless, this result demonstrates that there is potential in the use of more sophisticated approaches than just selecting all subpopulation with below average Nei distance (Nei 1972) as the reference panel. For a detailed list containing error rates for all four different structures of reference panels, we refer to Supplementary Table S7.

## DISCUSSION AND CONCLUSIONS

### Significance of improvement

When comparing error rates under different parameter settings one has to keep in mind the relevance of that optimization. A difference in error rates of 1% in a dataset containing 1% missing genotypes will only result in an improved overall data quality of 0.01% and thus might be negligible compared to other error sources like calling errors (Unterseer *et al*. 2014). If those improvements would mainly occur in the markers of interest (e.g. markers with low minor allele frequency) or the overall share of missing positions is high (as in UM imputation), this improvement could still be significant for later steps of the analysis.

It should be noted that positions set to NA in this study are chosen at random whereas in a real dataset there might be causal reasons like deletions, leading to some markers with much higher missing rates. When performing imputation on the actual NAs, we observed a higher variance in the imputed allele under different random seeds. As all considered methods always input one of the two allelic variants, this is ignored here but it should be noted that actual error rates are probably a bit higher than reported in this study.

### Genetic map and DR2

The used reference genome only mildly affected overall error rates in maize. As the number of markers with extremely high error rates is reduced, we still recommend the use of a reference genome of a more related individual. This of course requires its existence and similar overall quality. The overall gains should not be high enough to justify the generation of a new reference genome just for imputation. Instead one could consider either removing critical markers from the set or use imputation methods like LinkImpute (Money *et al*. 2015) that do not rely on a genetic map.

We highly recommend the use of DR2 to check validity of results obtained in BEAGLE 5.0 and 5.1. Firstly, observation of a high number of low values of DR2 can be seen as an indicator of overall poor imputation quality. Secondly, one should consider removing markers with low values for DR2 as error rates of UM imputation are typically massively increased. Here, one has to find a balance between removing potentially informative high quality markers and working with low quality markers that could potentially lead to false positive results in later steps of an analysis. In any case, markers that tend to have large effects (e.g. in a genome-wide association study) should be checked for their DR2 value.

### Reference panel

Without any knowledge of the genetic structure or excessive testing of genetic relatedness, we recommend to use all available individuals genotyped under high marker density for the reference panel, as the BEAGLE algorithm seems to be quite good at filtering out irrelevant information. However, in case most of the genetic diversity of the study sample can be represented in a subset of the default with B73v4 (Jiao *et al*. 2017) as a reference genome. individuals in a reference panel (e.g. a reference panel containing all founder individuals), significant improvements to UM imputation performance can be made by excluding genetically distant individuals. Representing a high share of the genetic diversity of a dataset however is far more important as error rates increase massively if no genomic data of highly related individuals is available in the reference.

### Parameter adaptation

Overall, we can conclude that the quality for inference, UM imputation and phasing in BEAGLE 5.0 and 5.1 was better or at least as good as in BEAGLE 4.0 and 4.1 and less tuning of parameters is necessary to obtain good performance for livestock and crop datasets. However, even in BEAGLE 5.0 and 5.1 the adaptation of the parameter *ne* is absolutely necessary when working with genetic datasets with less diversity than a human outbred population. Especially when no parameter tuning in BEAGLE 4.0 and 4.1 was done, one should consider re-running previous preprocessing and quality control protocols. However, a switch from BEAGLE 5.0 to 5.1 is not necessary, nor even recommended as error rates for phasing (and thereby inference and UM imputation) were lower in BEAGLE 5.0. It should be noted that all datasets in this study contain less genetic diversity than an outbred human population and for datasets with higher genetic diversity like those of UK Biobank (http://www.ukbiobank.ac.uk/) BEAGLE 5.1 is supposed to have around 25% lower error rates (B. Browning, personal communications).

Especially for UM imputation and in case of heterozygous individuals an increase of the number of iterations improved results slightly. As long as computing time is no issue we suggest to increase the number of iterations. As the gains by a higher number of iterations is relatively low one can also consider reducing the number of iterations to 4 (or in case of DHs to 2) for large datasets which will dramatically reduce computing time.

Other than in the case of *ne* for UM imputation, improvements in BEAGLE 5.0 and 5.1 by parameter tuning are relatively small, leading us to conclude that the use of default settings should be enough for most applications. Especially for datasets with relatively low genetic diversity one should consider increasing the parameters *phase-segments*, *imp-segments* and *window* while reducing *imp-states* and *ne*. For substantial changes of the imputing parameters and for maximizing the imputing accuracy we strongly suggest to apply a testing pipeline similar to the one suggested in the methods section. As potential gains should not be much higher than 5-10% one has to decide based on the application if this additional effort is worth it. Obtainable improvements in BEAGLE 4.0 and 4.1 are high but we do not recommend to use these versions anymore. Additional benefits of the use of BEAGLE 5.0 and 5.1 are massively reduced computing times and memory requirements. Two potential exceptions to this are if high quality pedigree is available, as only BEAGLE 4.0 is able to incorporate pedigree data in its imputation algorithm and in case only genotype likelihoods are available as input as BEAGLE 5.0 and 5.1 only allow for genotypes as input.

## ACKNOWLEDGEMENTS

The authors thank the German Federal Ministry of Education and Research (BMBF) for the funding of our project (MAZE - “Accessing the genomic and functional diversity of maize to improve quantitative traits”; Funding ID: 031B0195). The “Synbreed - Synergistic Plant and Animal Breeding” project was funded by the German Federal Ministry of Education and Research (FKZ 0315528E).

We also thank Brian Browning for providing quick and thoughtful replies to all our questions regarding insights into BEAGLE and providing us with personalized software updates for BEAGLE 5.1.

## 1. SUPPLEMENTARY TABLES/FIGUES

**Table S1** UM imputation error for the commercial breeding line in chicken by changing a single imputing parameter with *ne* =100 for BEAGLE 5.1, *ne* = 300 for BEAGLE 5.0 and *ne* = 10,000 for BEAGLE 4.1. * BEAGLE crashed for this dataset when using *phase-segment* > 10, *phase-states* < 100 or *imp-step* > 5.

**Table S2** UM imputation error for the diversity panel in chicken by changing a single imputing parameter with *ne* = 300 for BEAGLE 5.1, *ne* = 3,000 for BEAGLE 5.0 and *ne* = 10,000 for BEAGLE 4.1. * BEAGLE crashed for this dataset when using *phase-segment* > 10 or *phase-states* < 100.

**Table S3** Phasing error, as number of heterozygous markers per switch error, for Pseudo *S*_0_ generated based on the KE DH-lines by changing a single imputing parameter.

**Table S4** Inference error rates using different reference genomes compared to B73 for PE DH-lines. Only markers mapped on both the flint reference genome & B73v4 (Jiao *et al*. 2017) are considered for “critical” markers (error rate > 10%).

**Table S5** List of “critical” markers for Kemater Landmais Gelb using different reference genomes.

**Table S6** List of “critical” markers for Petkuser Ferdinand Rot using different reference genomes.

**Table S7** Error rates for UM imputation for different reference panels, including A (same subpopulation), C (all other subpopulation), D (below average Nei distant subpopulations), E (All subpopulation with reduced error rate when testing A + B compared to A as the reference panel).

**Table S8** Assignments to subpopulations for the chicken diversity panel based on Nei standard genetic distances (Nei 1972).

**Table S9** Minimal obtained error rates and used parameter settings for inference and UM imputation. Deviations from the ideal single parameter settings are caused by BEAGLE crashing when changing parameters jointly.

**Figure S1.**
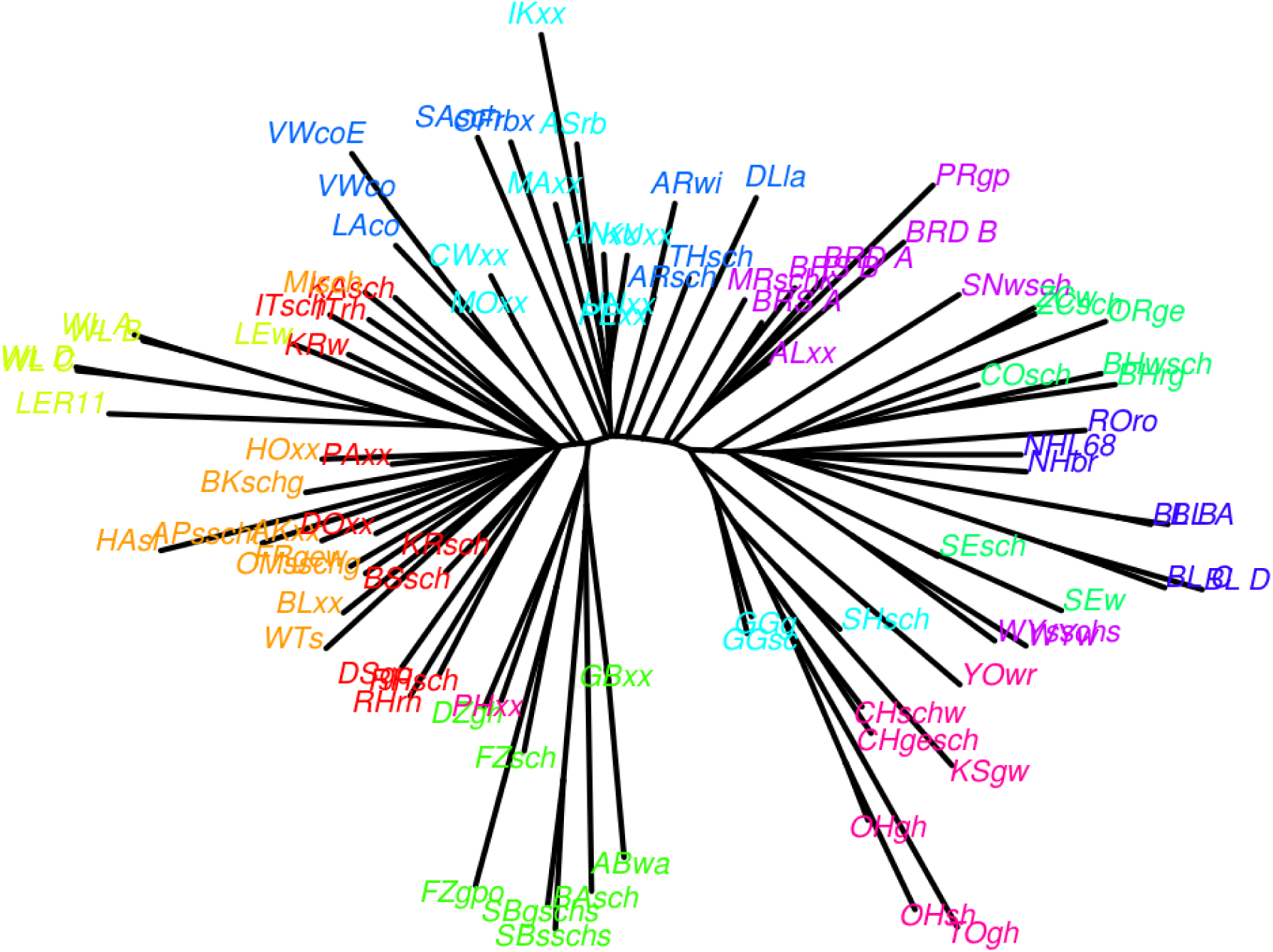
Neighbor-joining-tree for ten subpopulations in the chicken diversity panel. For a detailed list on which individual is assigned to which subpopulation we refer to Supplementary Table S8.

**Figure S2.**
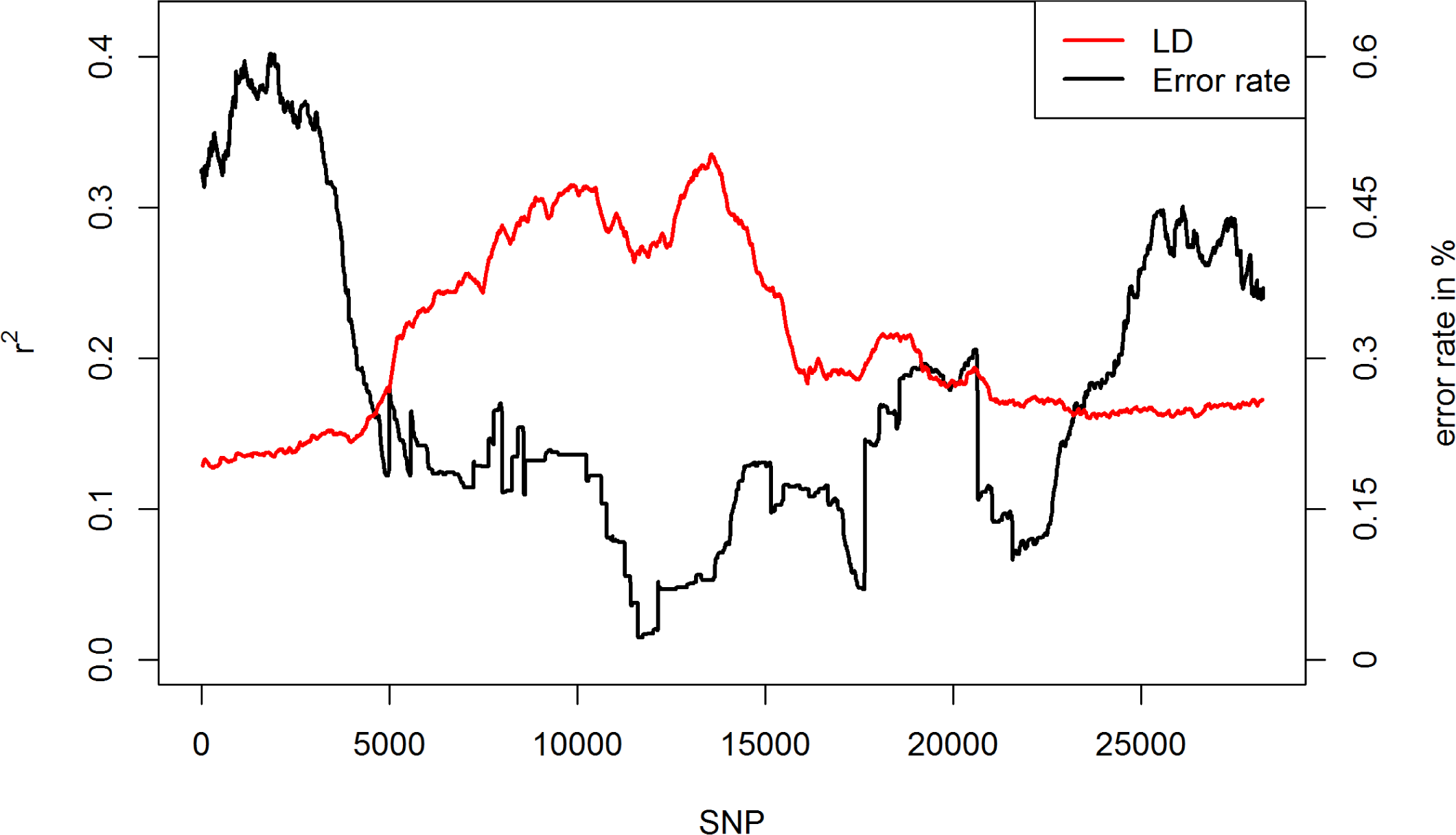
Relationship between region error rate and LD (*r*^2^) on chromosome 9 in the maize data. Outliers are corrected for by using a Nadaraya-Watson-estimator (Nadaraya 1964), using a Gaussian kernel and a bandwidth of 3,000 markers in both cases.

**Figure S3.**
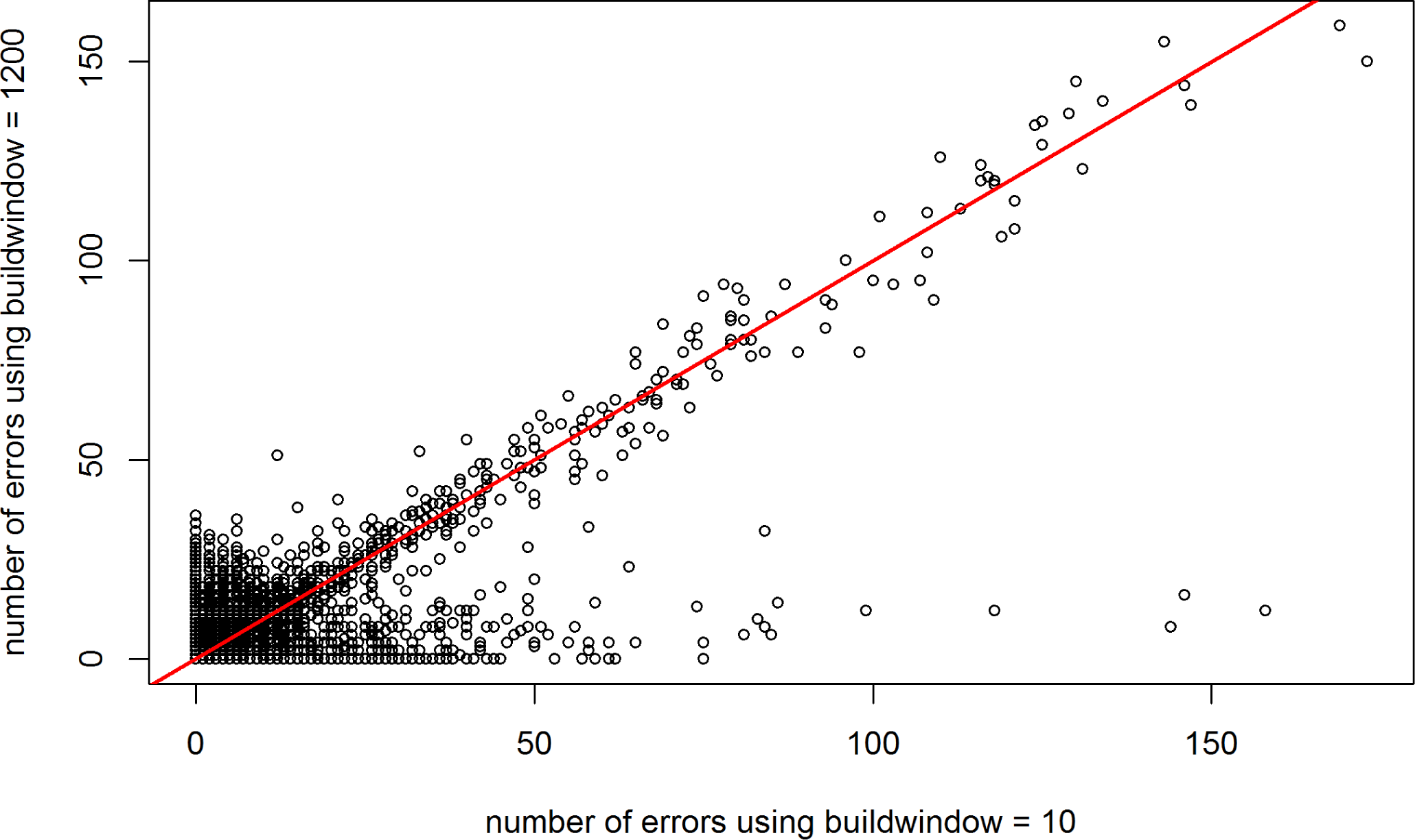
Total number of errors per marker (50 repetitions) for BEAGLE 4.0 using *buildwindow* of 10 and 1200 (default) in the maize data.

**Figure S4.**
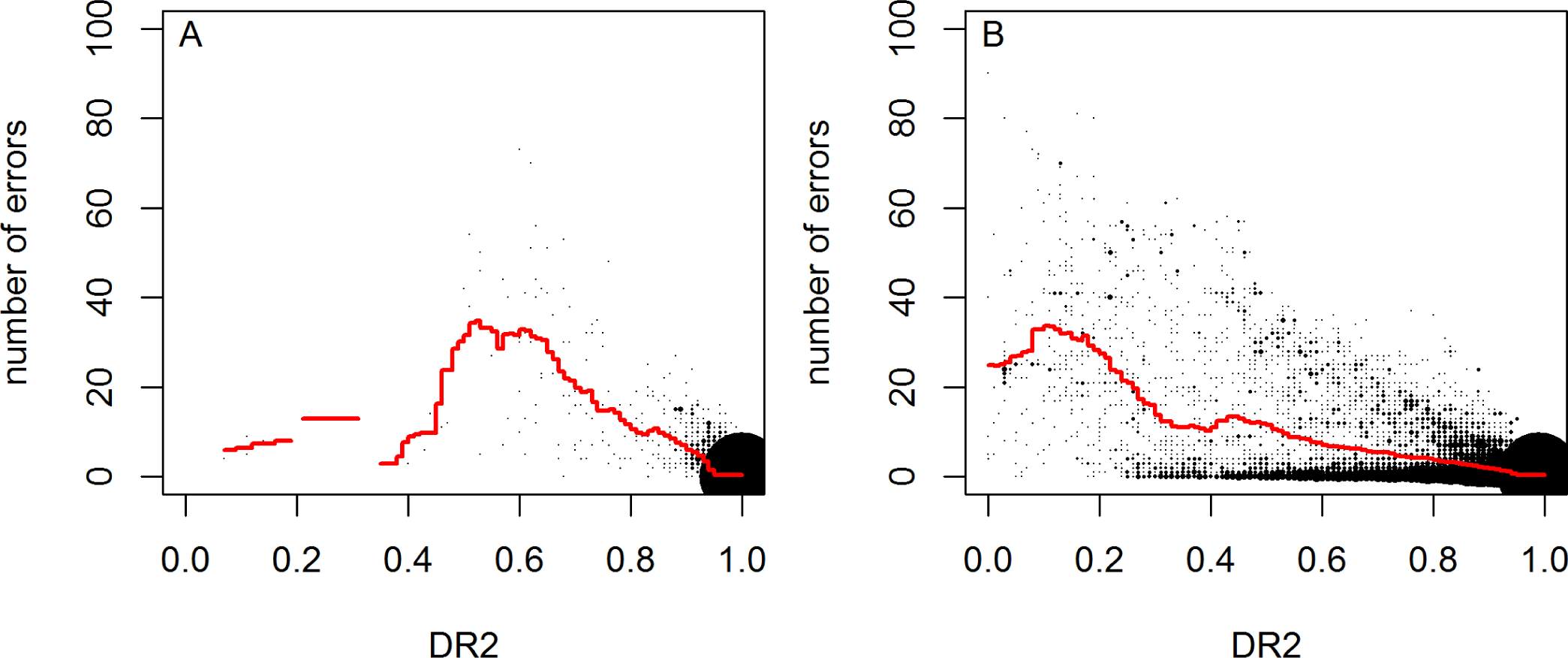
DR2 values in relation to the obtained number of error per marker after fitting of *ne* (A) and on default (B) in BEAGLE 5.0 for the commercial chicken line. 100 / 788 lines were used for study / reference sample.

**Figure S5.**
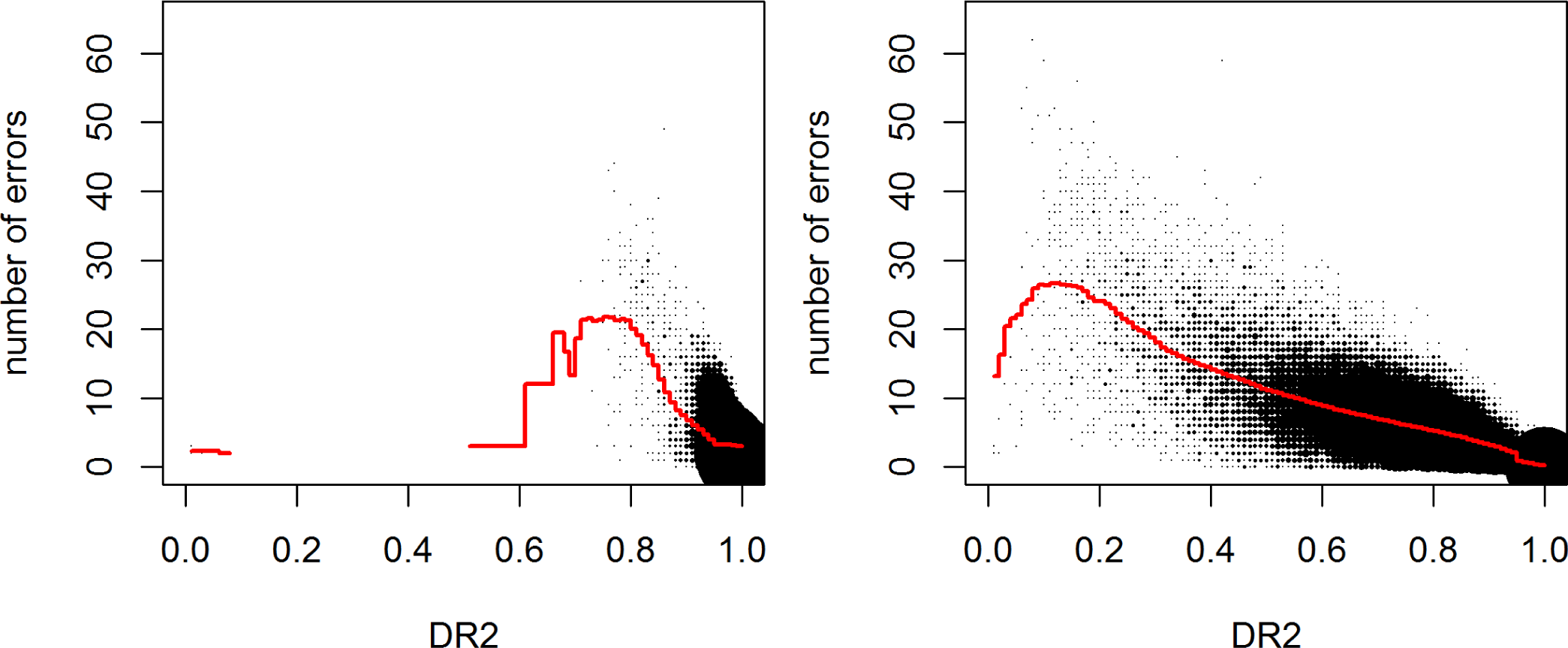
DR2 values in relation to the obtained number of error per marker after fitting of *ne* (A) and on default (B) in BEAGLE 5.0 for the chicken diversity panel. 100 / 1710 lines were used for study / reference sample.

**Figure S6.**
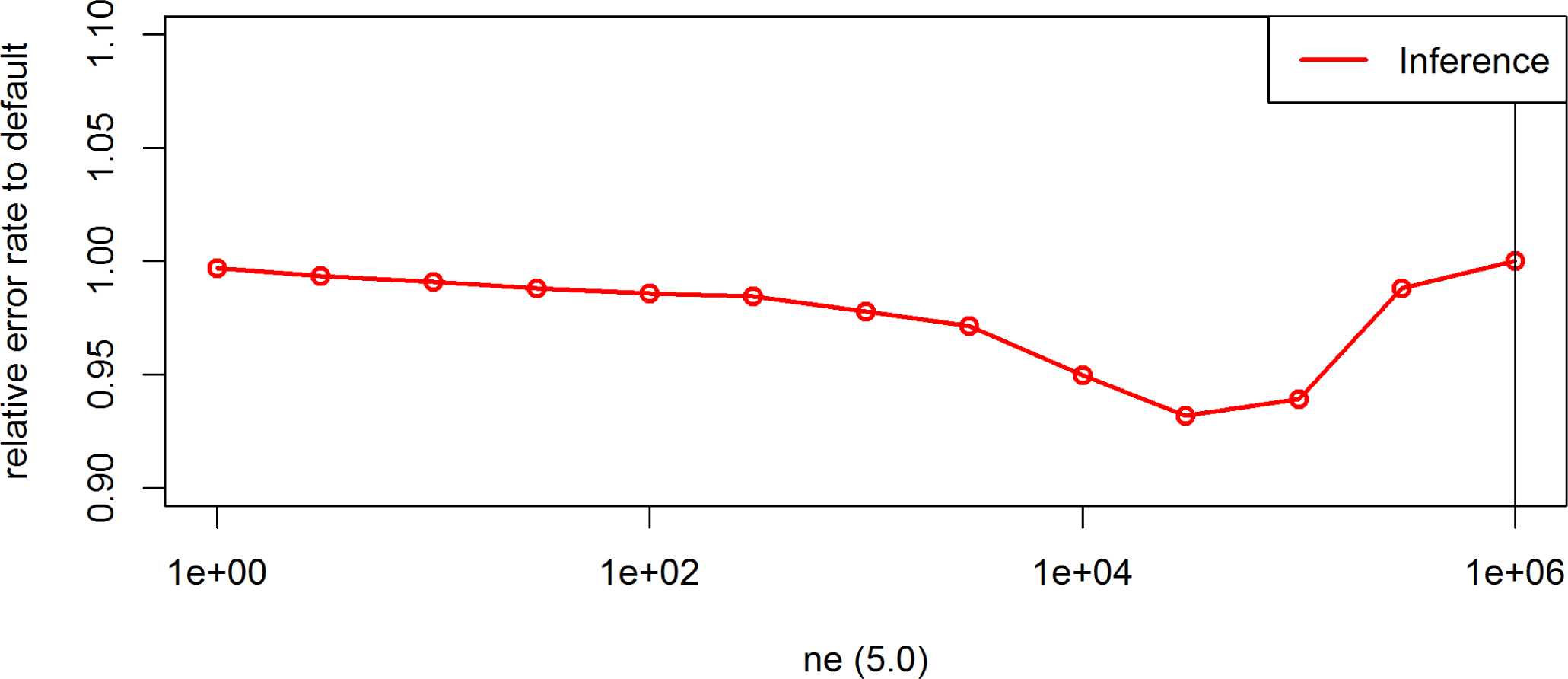
Effect of the parameter *ne* on the inference error rates for the maize data in BEAGLE 5.0. Default settings are indicated by the vertical line.

**Figure S7.**
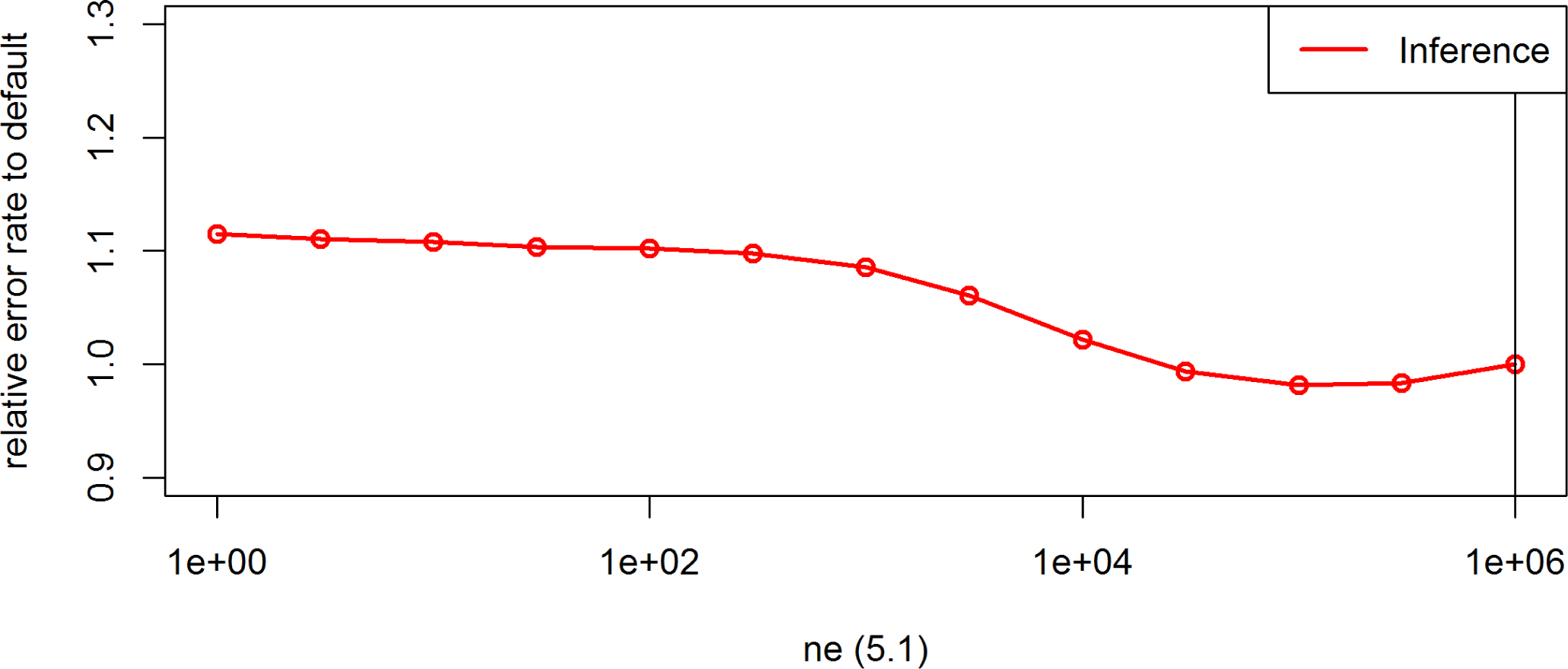
Effect of the parameter *ne* on the inference error rates for the maize data in BEAGLE 5.1. Default settings are indicated by the vertical line.

**Figure S8.**
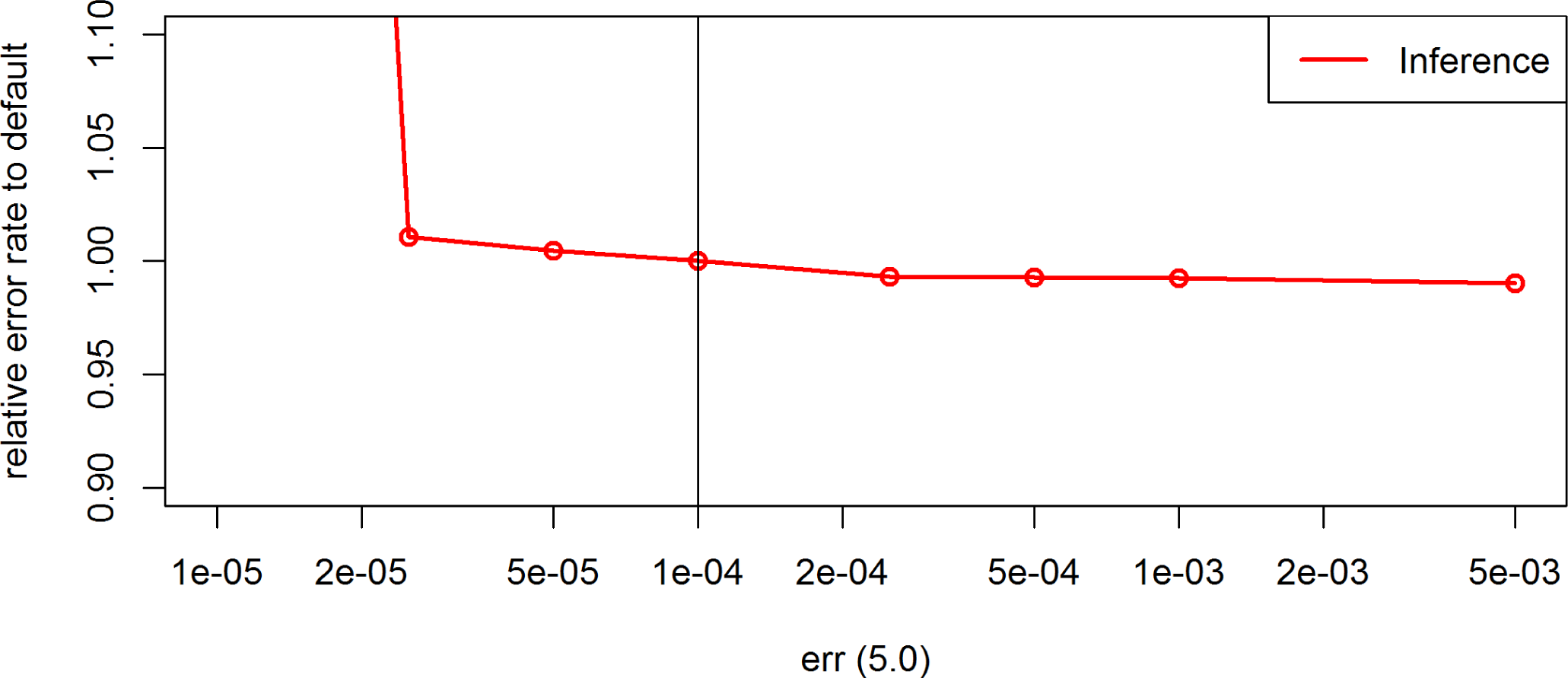
Effect of the parameter *err* on the inference error rates for the maize data in BEAGLE 5.0. Default settings are indicated by the vertical line.

**Figure S9.**
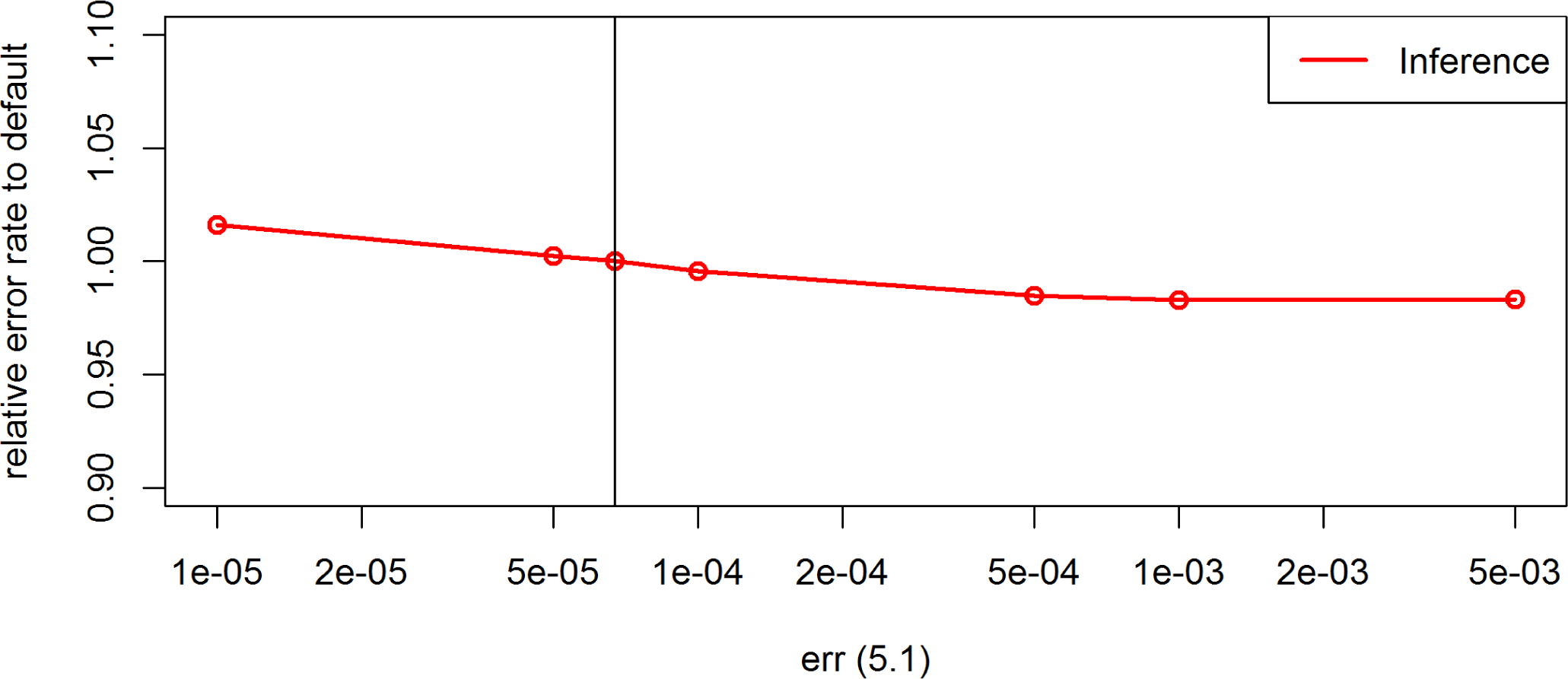
Effect of the parameter *err* on the inference error rates for the maize data in BEAGLE 5.1. Default settings are indicated by the vertical line.

**Figure S10.**
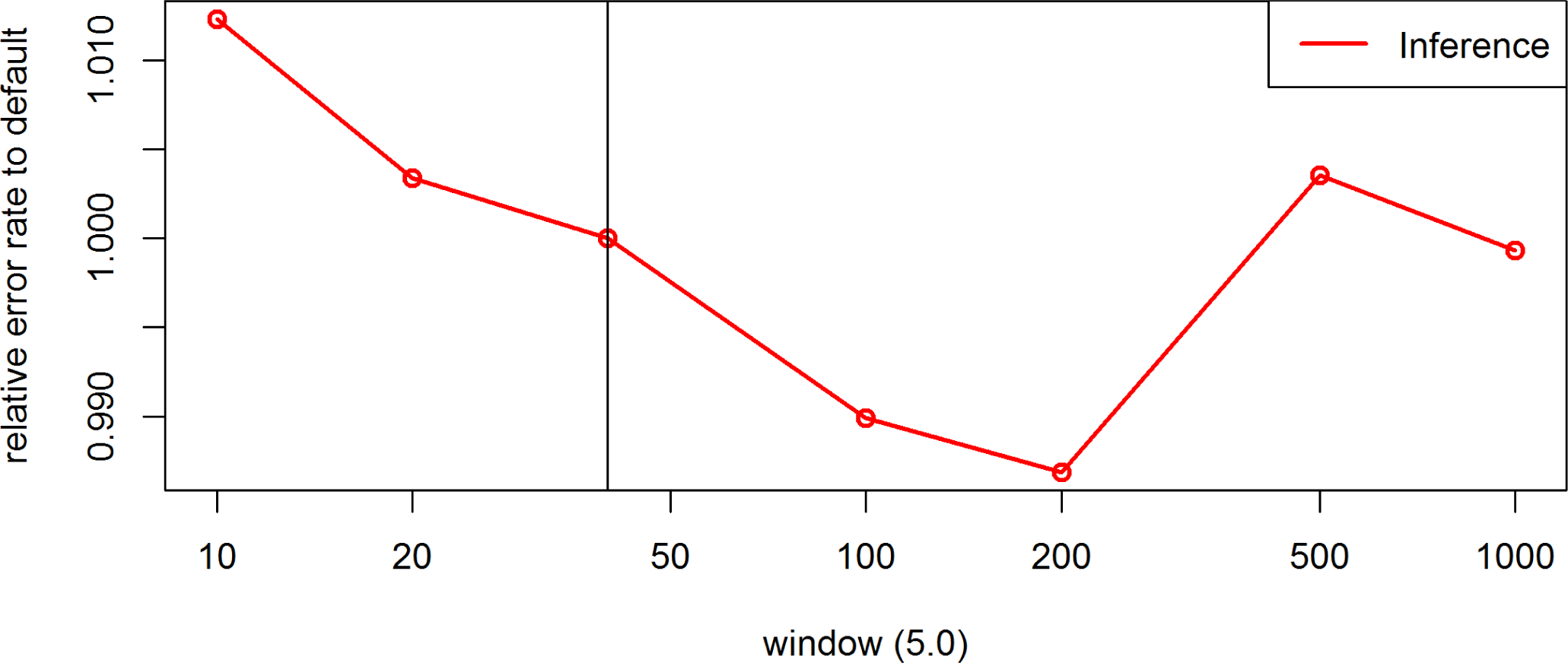
Effect of the parameter *window* on the inference error rates for the maize data in BEAGLE 5.0. Default settings are indicated by the vertical line.

**Figure S11.**
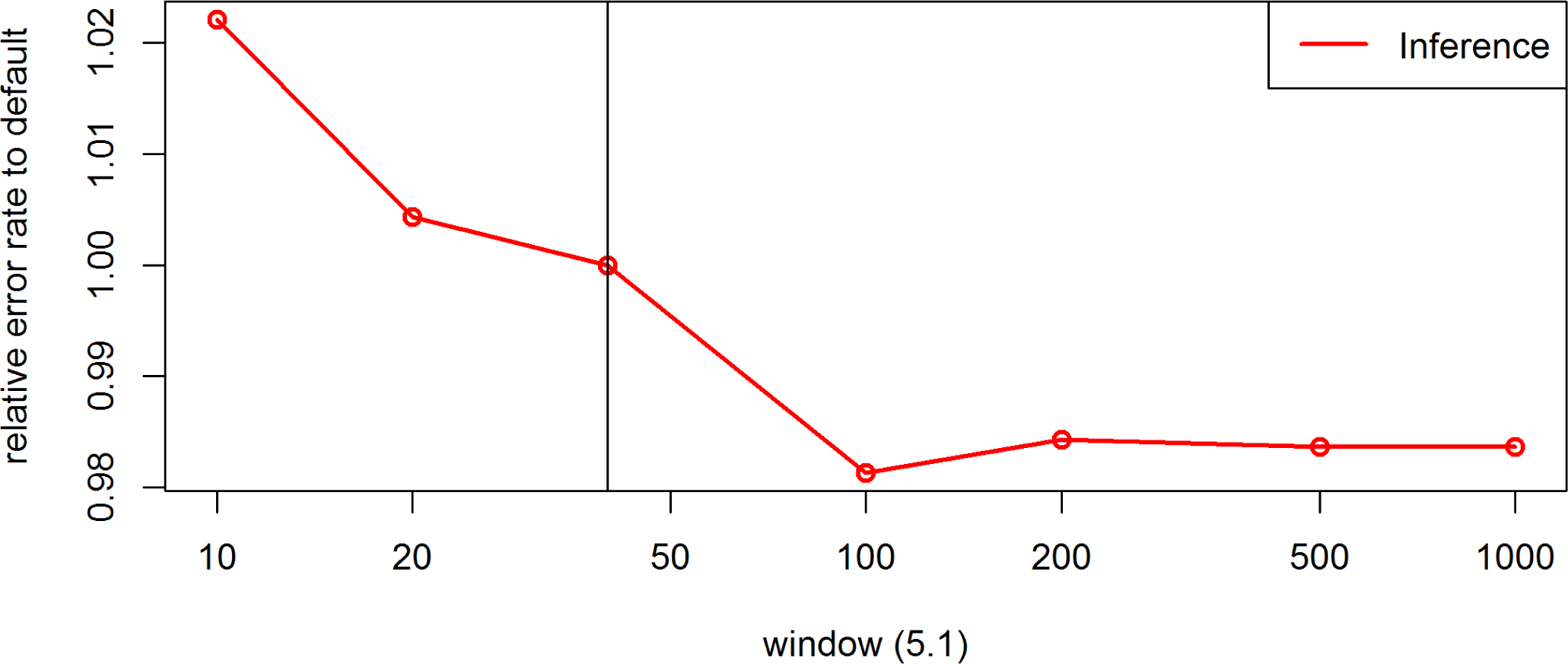
Effect of the parameter *window* on the inference error rates for the maize data in BEAGLE 5.1. Default settings are indicated by the vertical line.

**Figure S12.**
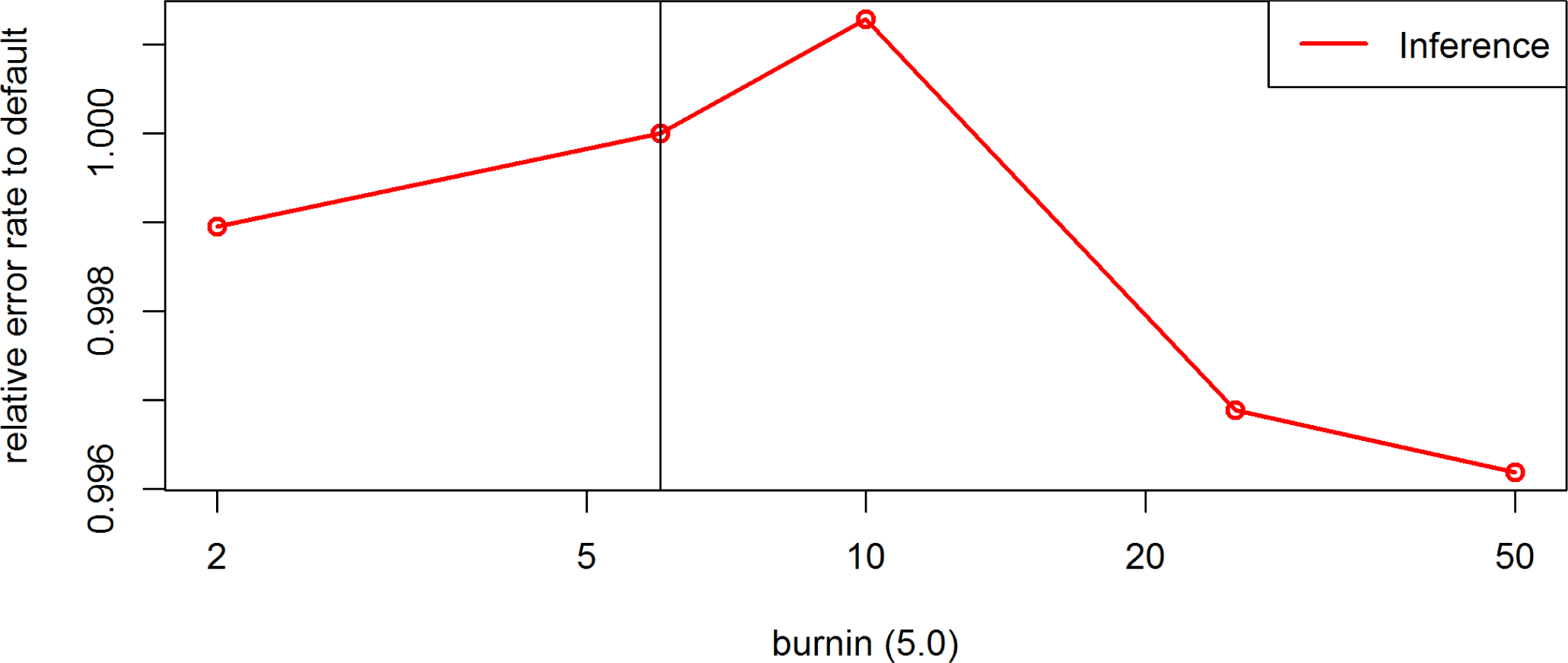
Effect of the parameter *phase-stages* on the inference error rates for the maize data in BEAGLE 5.0. Default settings are indicated by the vertical line.

**Figure S13.**
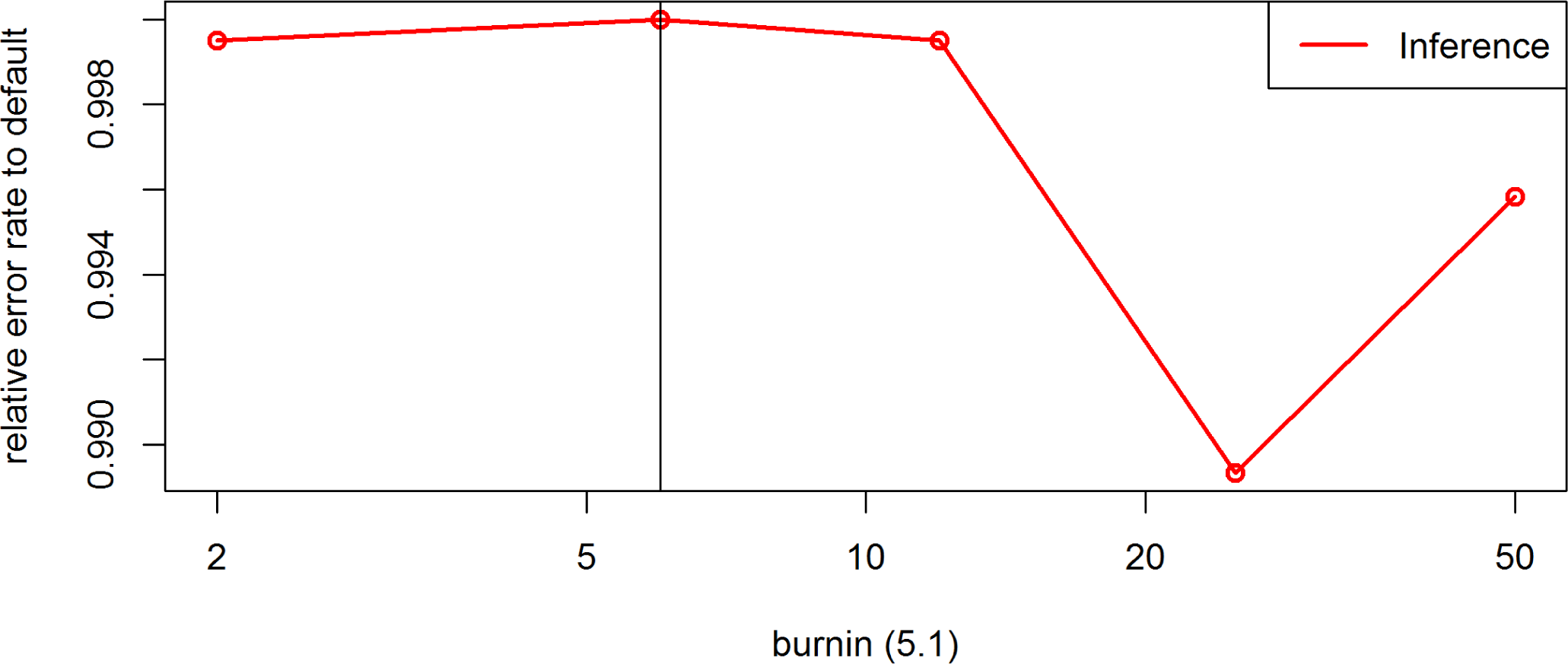
Effect of the parameter *phase-stages* on the inference error rates for the maize data in BEAGLE 5.1. Default settings are indicated by the vertical line.

**Figure S14.**
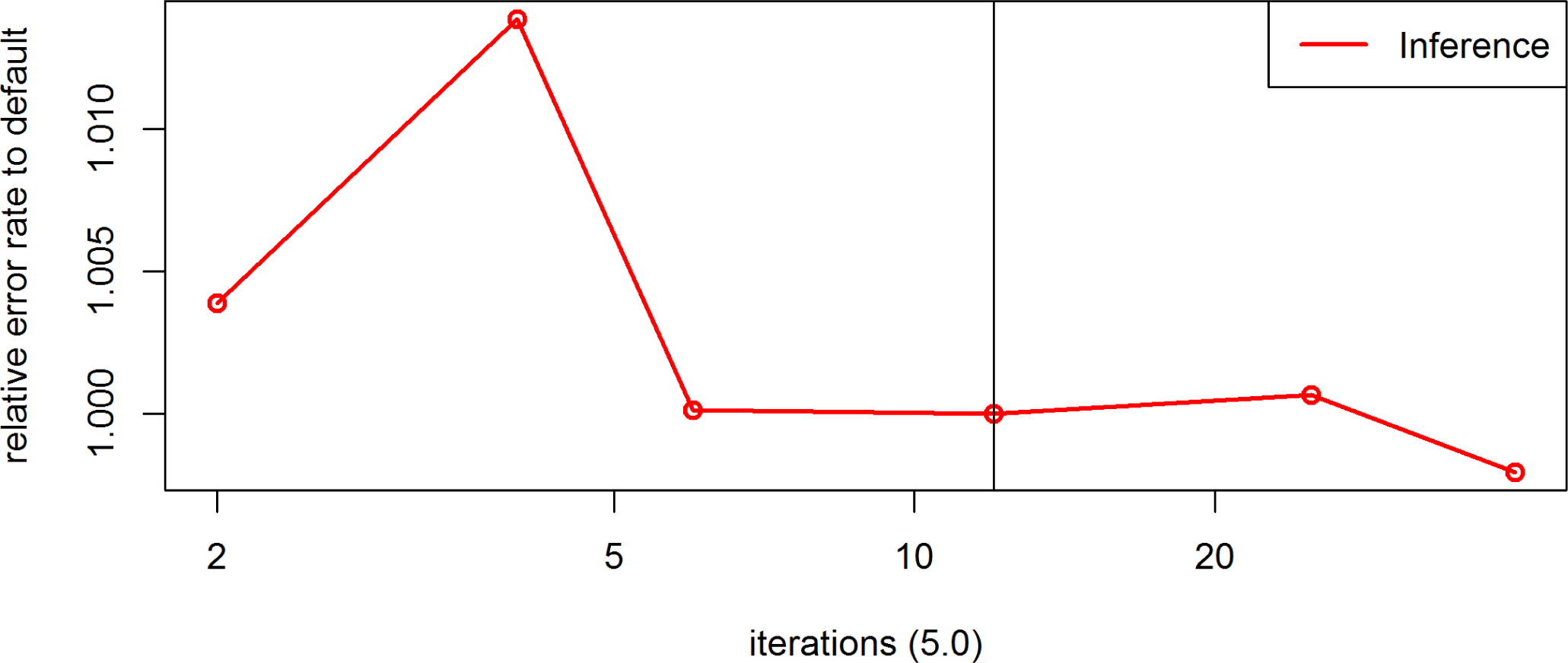
Effect of the parameter *phase-stages* on the inference error rates for the maize data in BEAGLE 5.0. Default settings are indicated by the vertical line.

**Figure S15.**
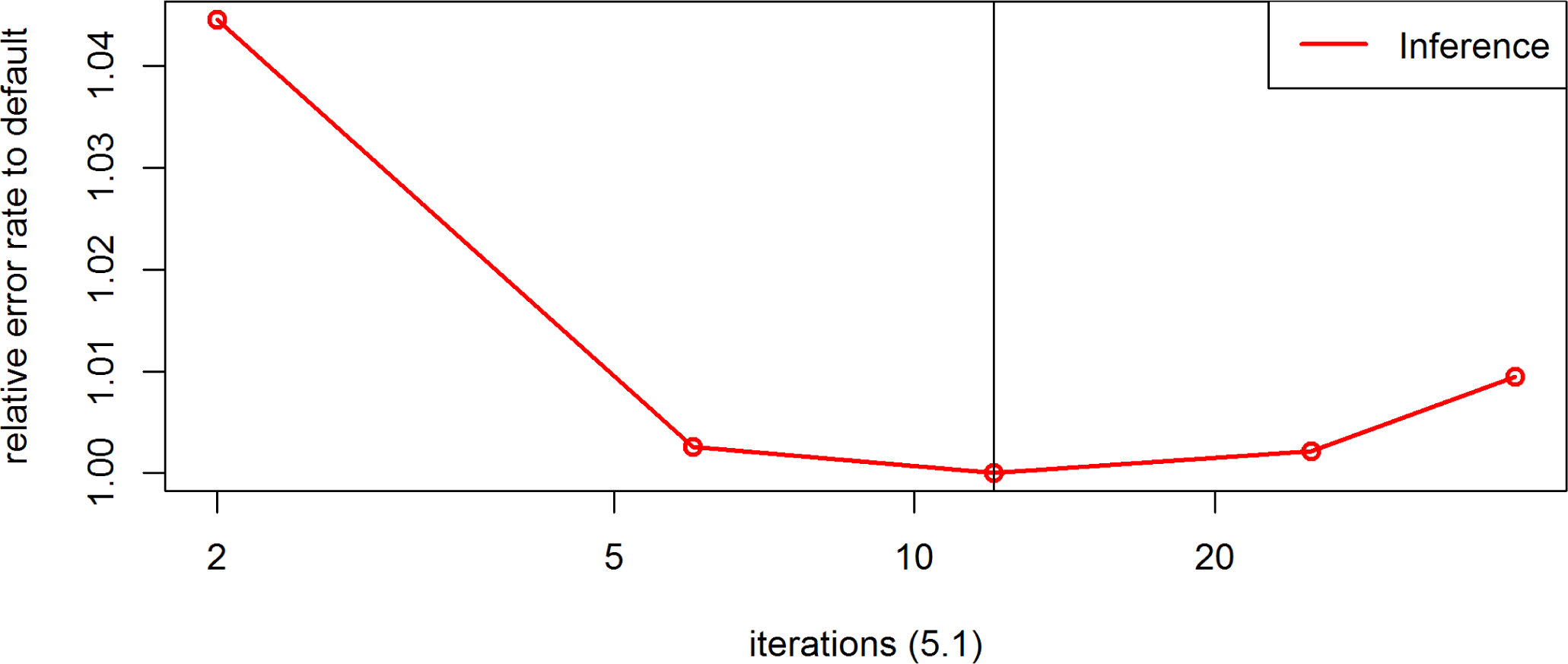
Effect of the parameter *phase-stages* on the inference error rates for the maize data in BEAGLE 5.1. Default settings are indicated by the vertical line.

**Figure S16.**
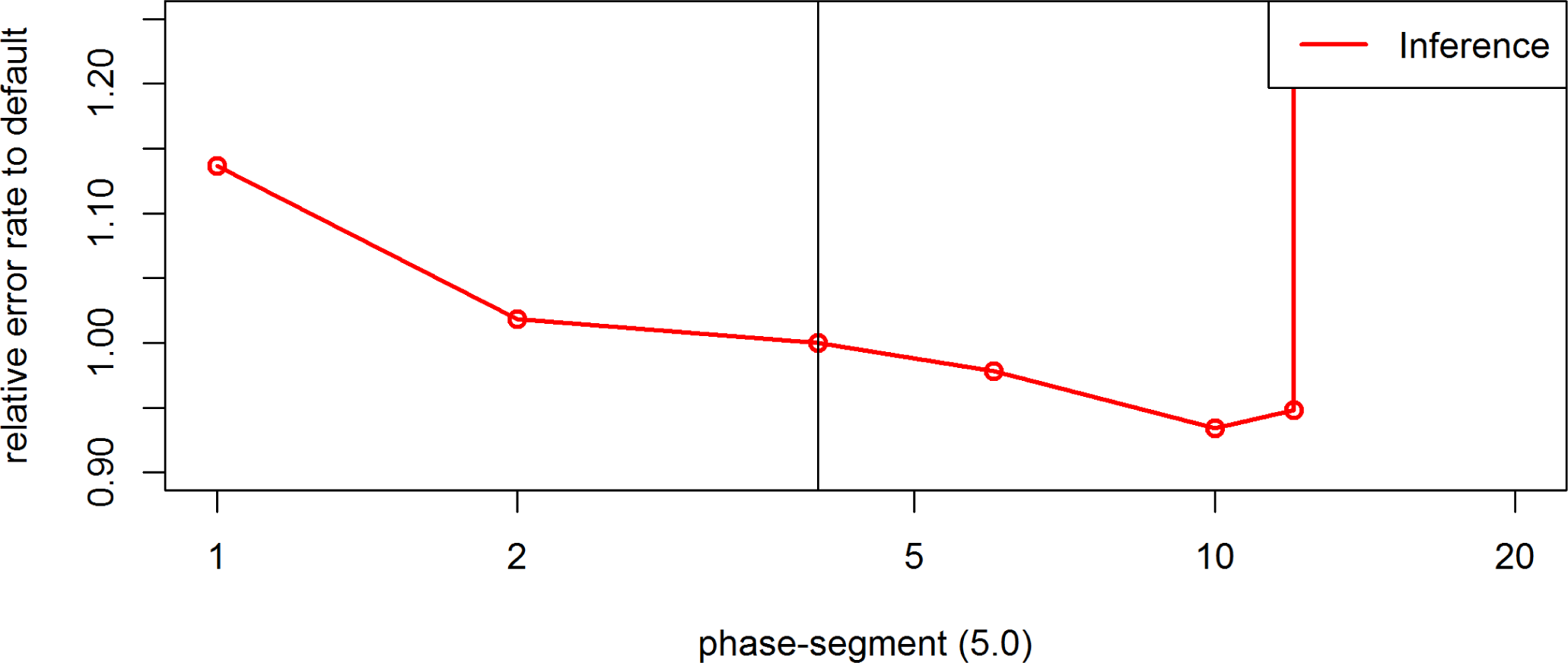
Effect of the parameter *phase-segment* on the inference error rates for the maize data in BEAGLE 5.0. Default settings are indicated by the vertical line.

**Figure S17.**
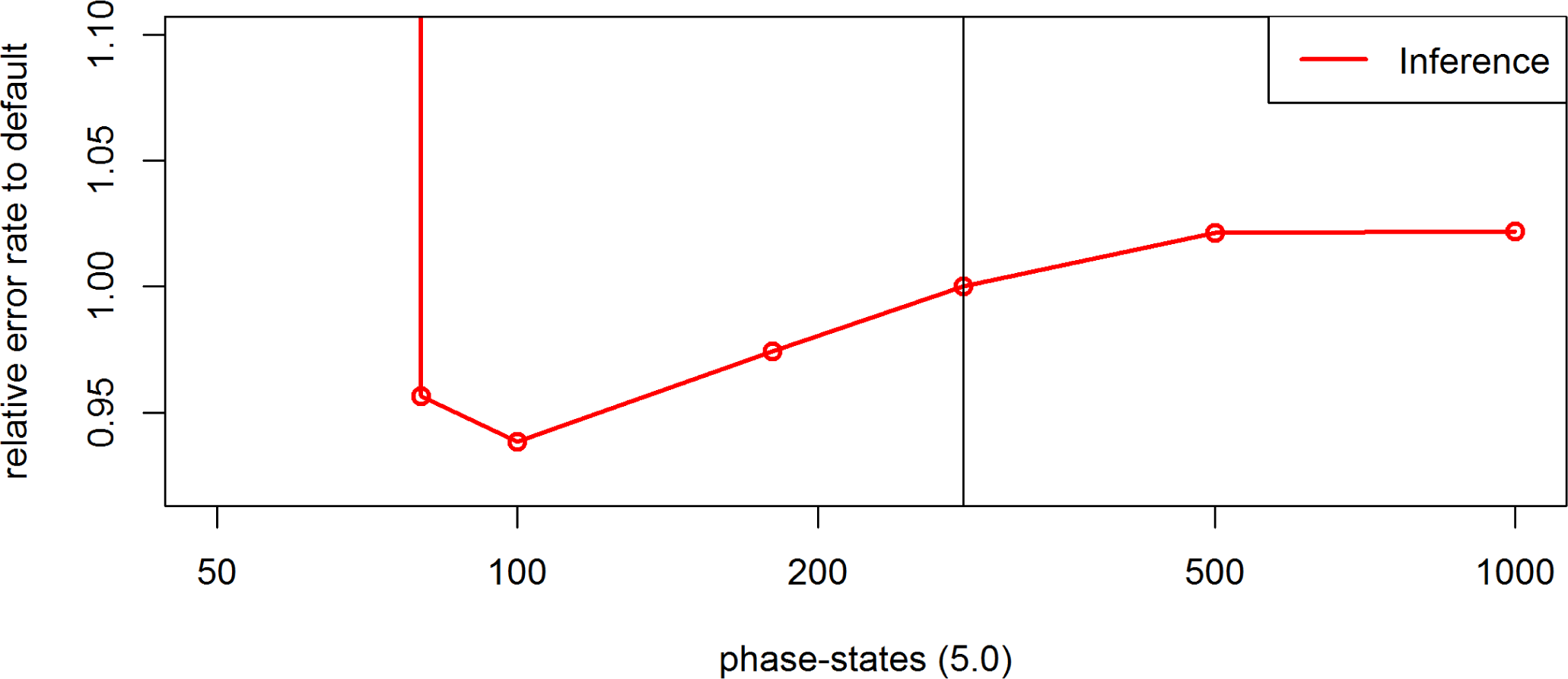
Effect of the parameter *phase-stages* on the inference error rates for the maize data in BEAGLE 5.0. Default settings are indicated by the vertical line.

**Figure S18.**
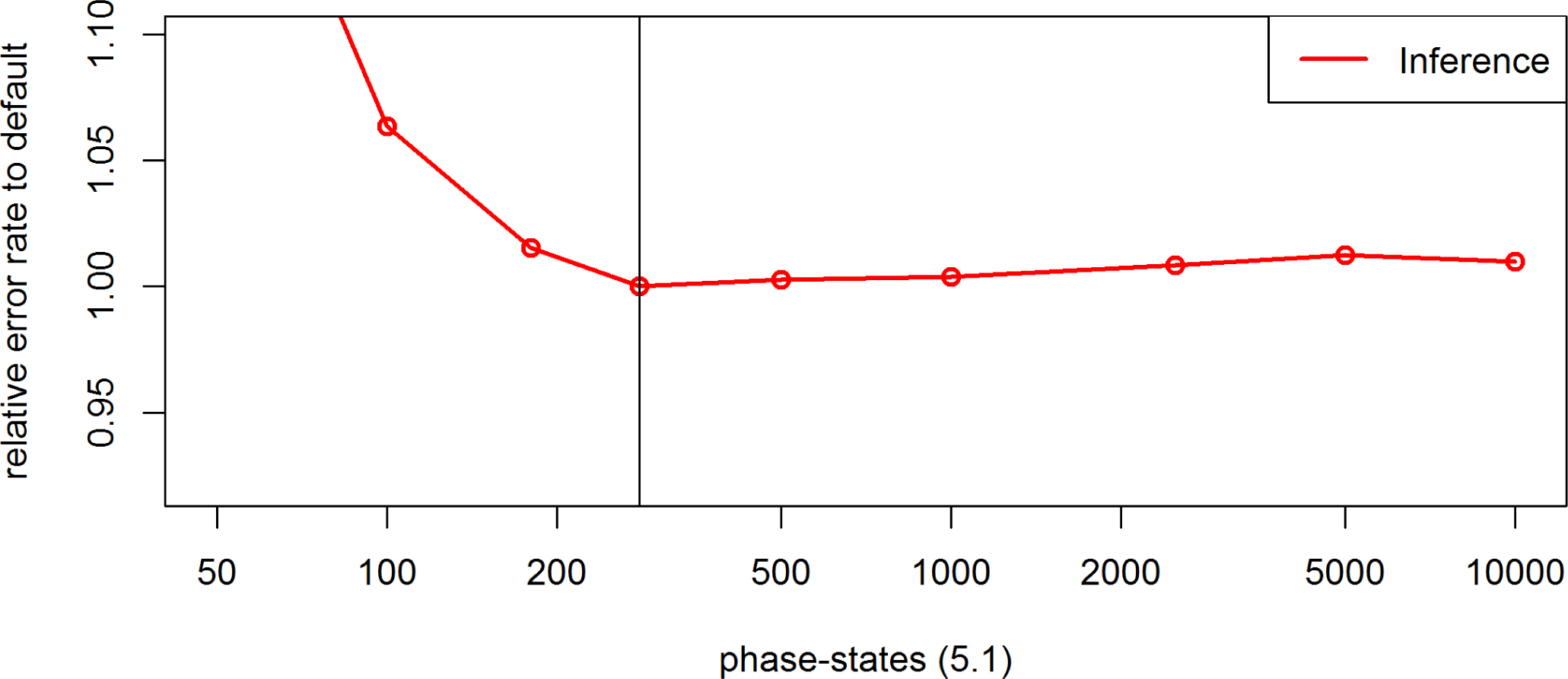
Effect of the parameter *phase-stages* on the inference error rates for the maize data in BEAGLE 5.1. Default settings are indicated by the vertical line.

**Figure S19.**
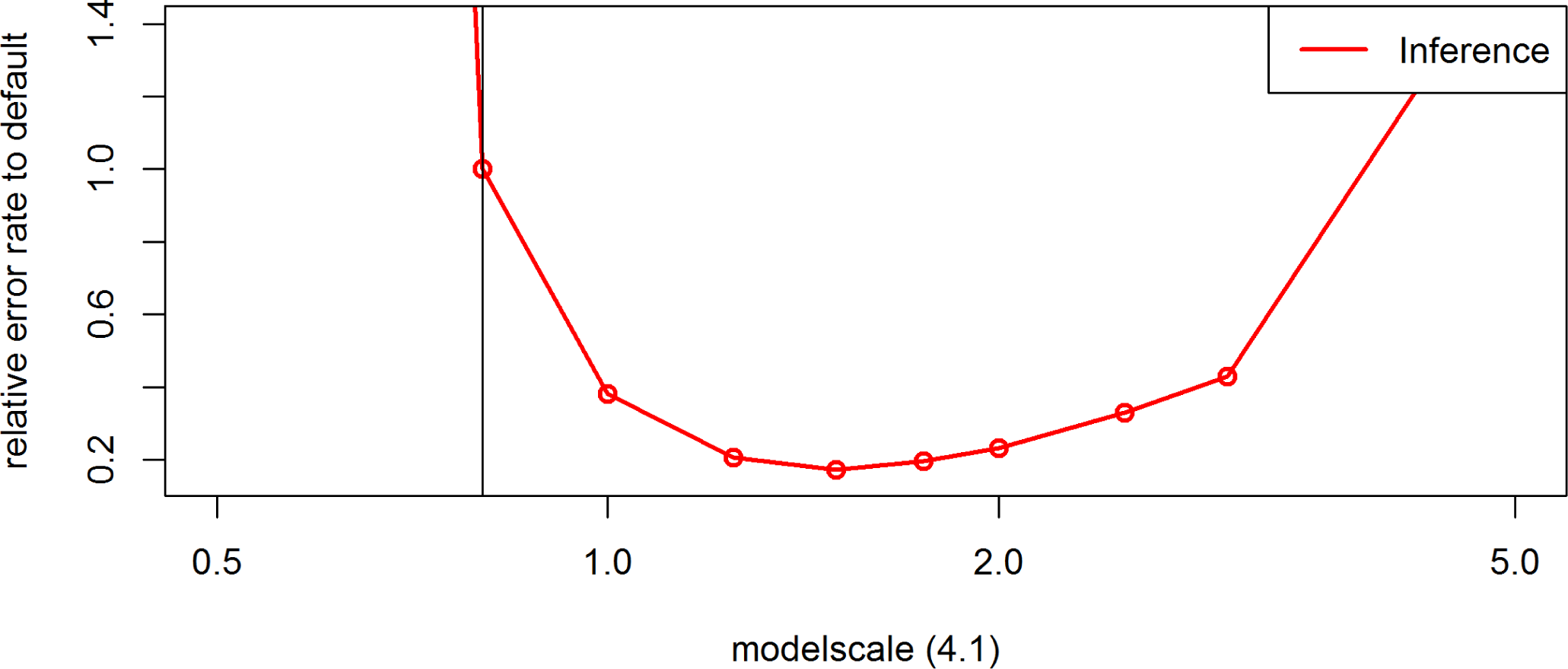
Effect of the parameter *modelscale* on the inference error rates for the maize data in BEAGLE 4.1. Default settings are indicated by the vertical line.

**Figure S20.**
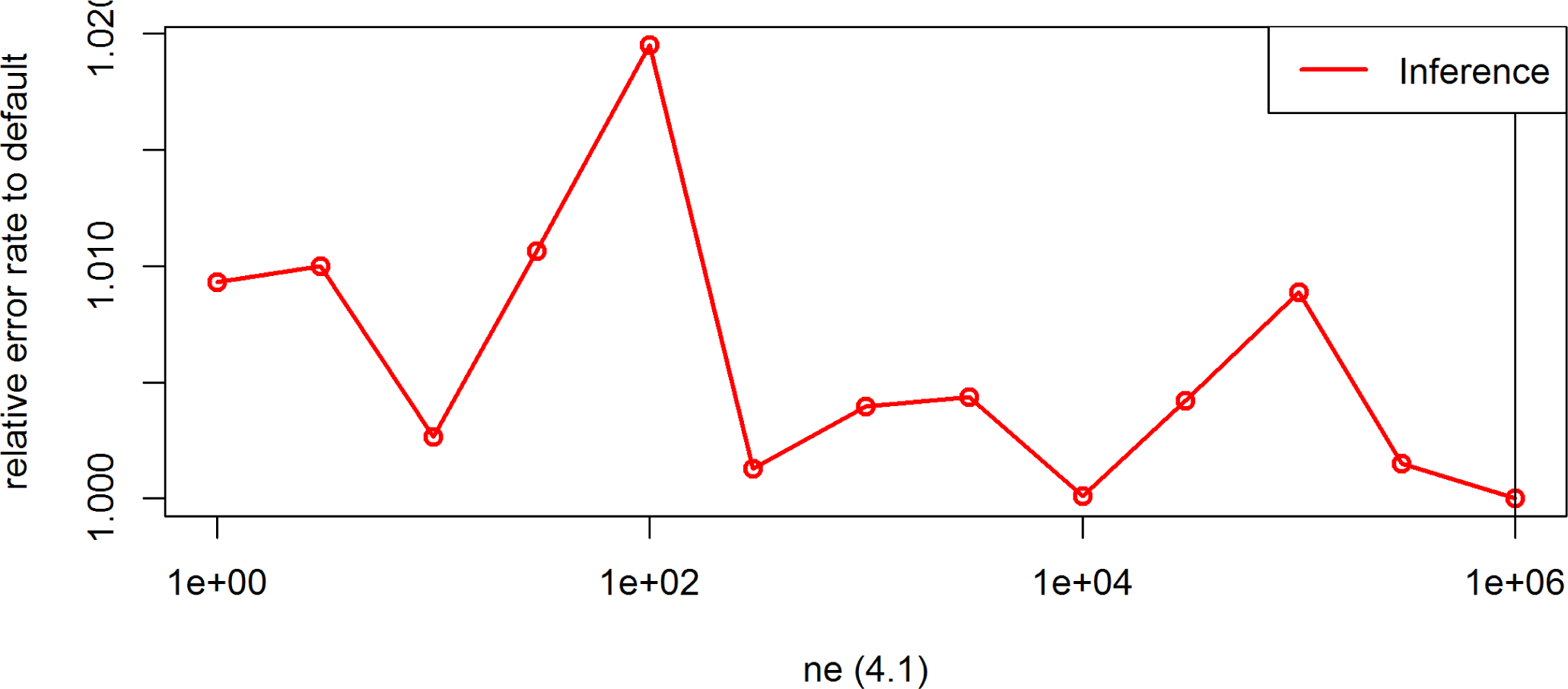
Effect of the parameter *ne* on the inference error rates for the maize data in BEAGLE 4.1. Default settings are indicated by the vertical line.

**Figure S21.**
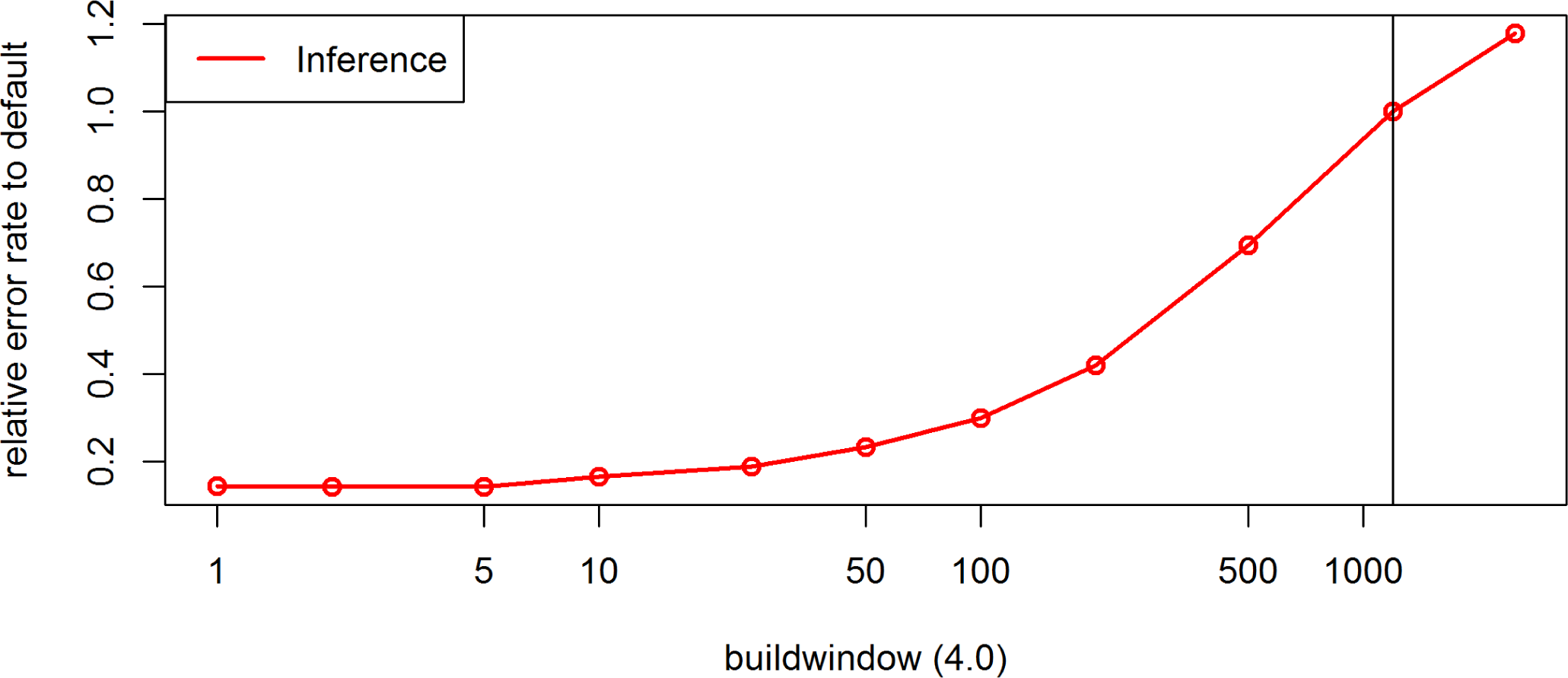
Effect of the parameter *buildwindow* on the inference error rates for the maize data in BEAGLE 4.0. Default settings are indicated by the vertical line.

**Figure S22.**
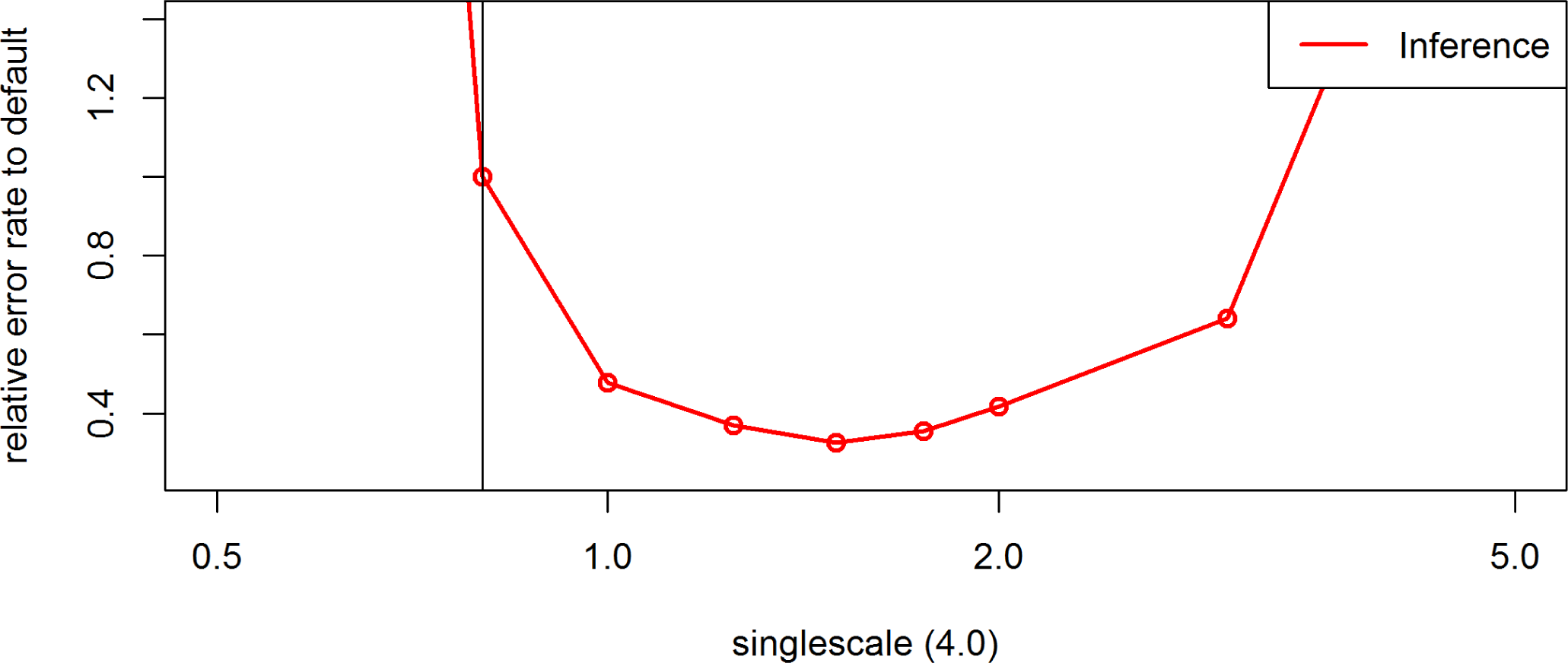
Effect of the parameter *singlescale* on the inference error rates for the maize data in BEAGLE 4.0. Default settings are indicated by the vertical line.

**Figure S23.**
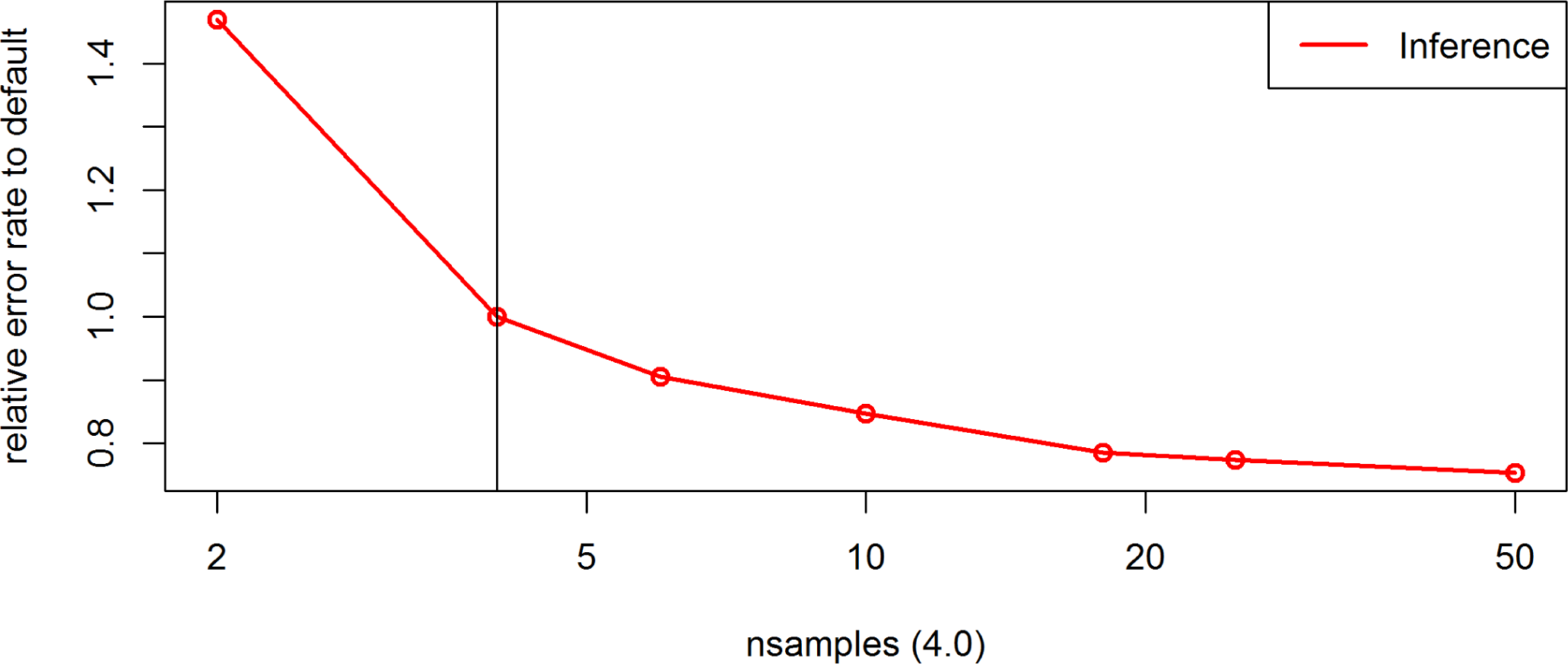
Effect of the parameter *nsamples* on the inference error rates for the maize data in BEAGLE 4.0. Default settings are indicated by the vertical line.

**Figure S24.**
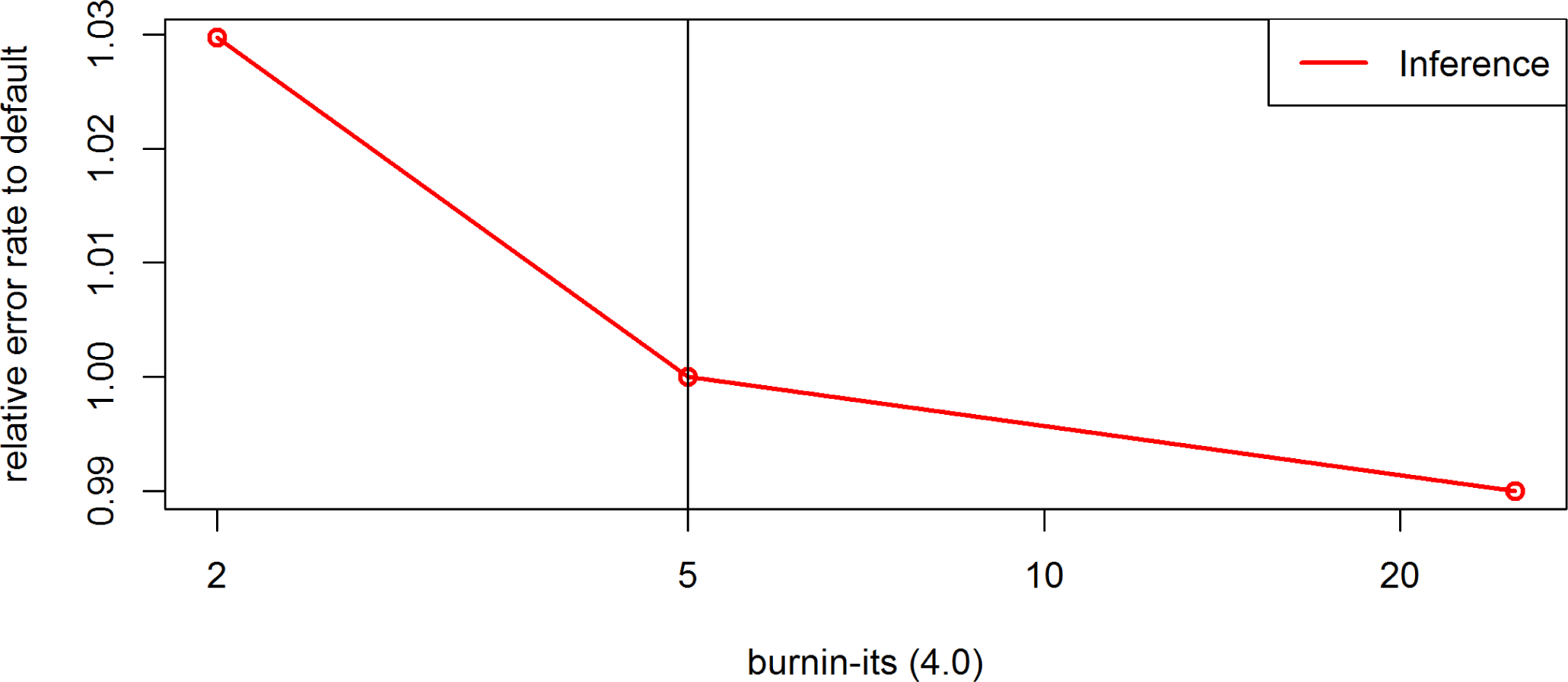
Effect of the parameter *burnin-its* on the inference error rates for the maize data in BEAGLE 4.0. Default settings are indicated by the vertical line.

**Figure S25.**
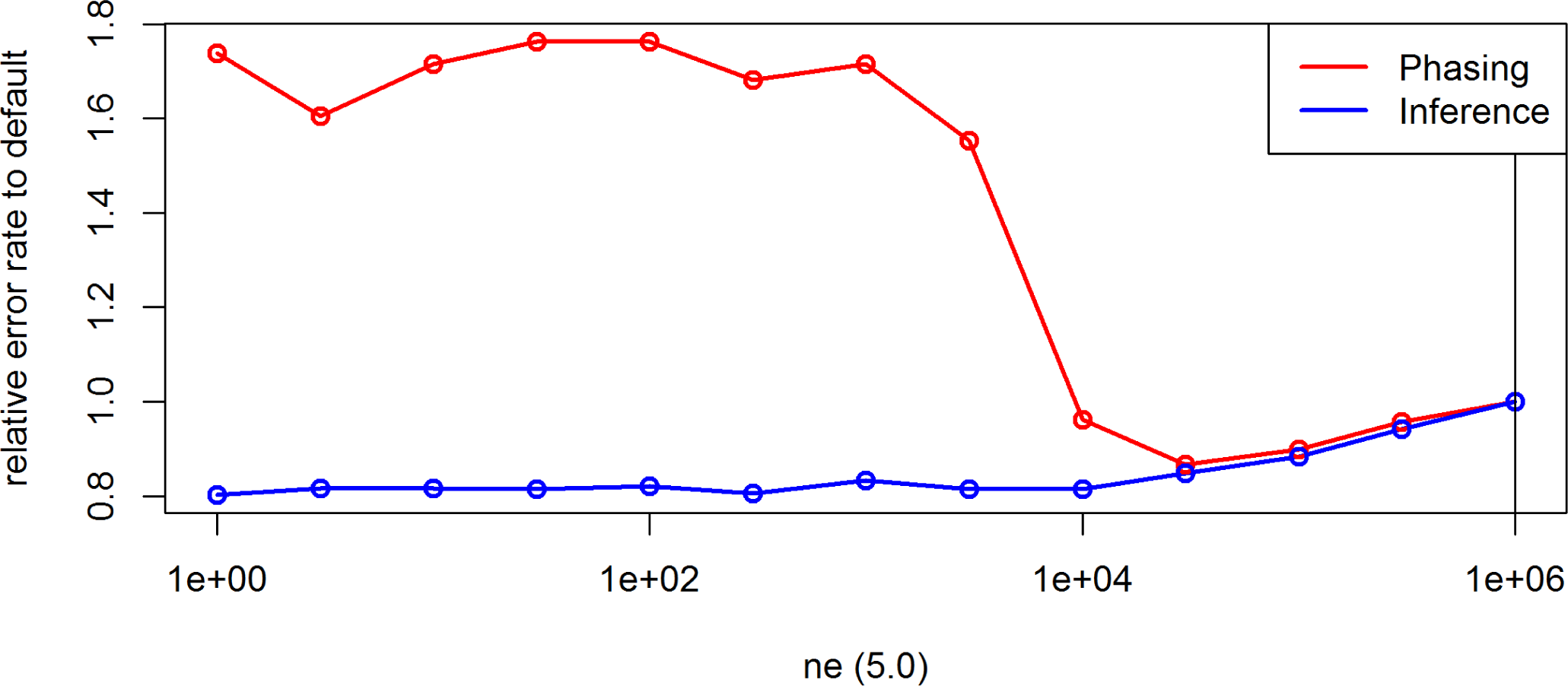
Effect of the parameter *ne* on the inference and phasing error rates for chromosome 10 of 250 Pseudo *S*_0_ generated based on the maize data in BEAGLE 5.0. Default settings are indicated by the vertical line.

**Figure S26.**
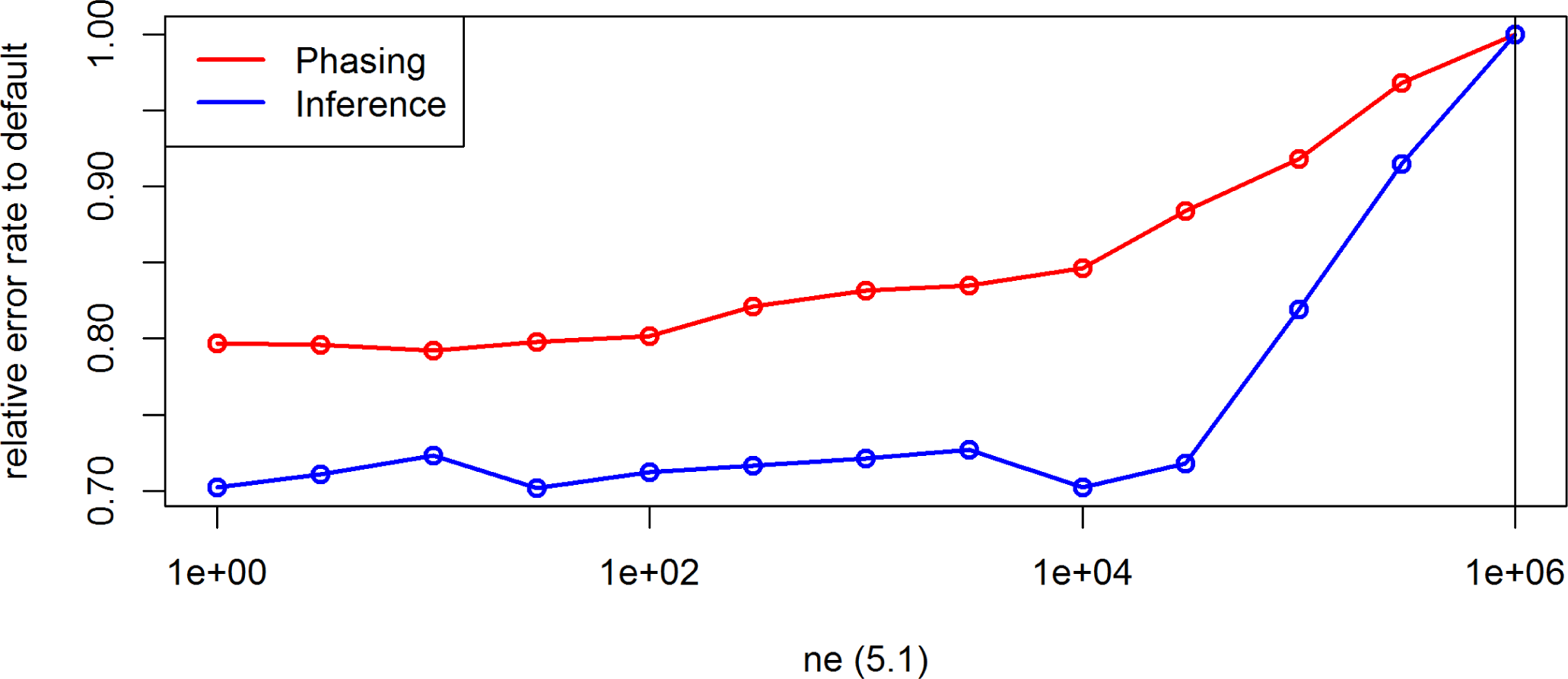
Effect of the parameter *ne* on the inference and phasing error rates for chromosome 10 of 250 Pseudo *S*_0_ generated based on the maize data in BEAGLE 5.1. Default settings are indicated by the vertical line.

**Figure S27.**
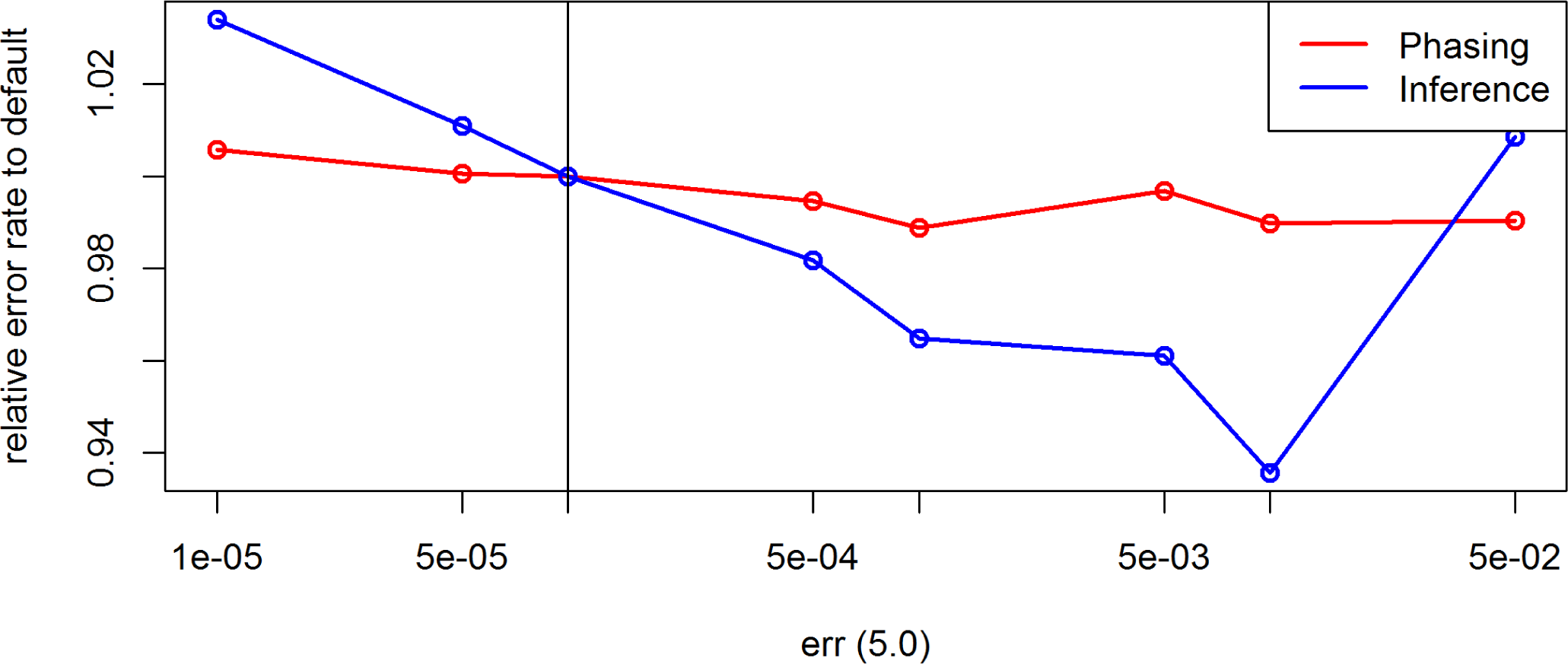
Effect of the parameter *err* on the inference and phasing error rates for chromosome 10 of 250 Pseudo *S*_0_ generated based on the maize data in BEAGLE 5.0. Default settings are indicated by the vertical line.

**Figure S28.**
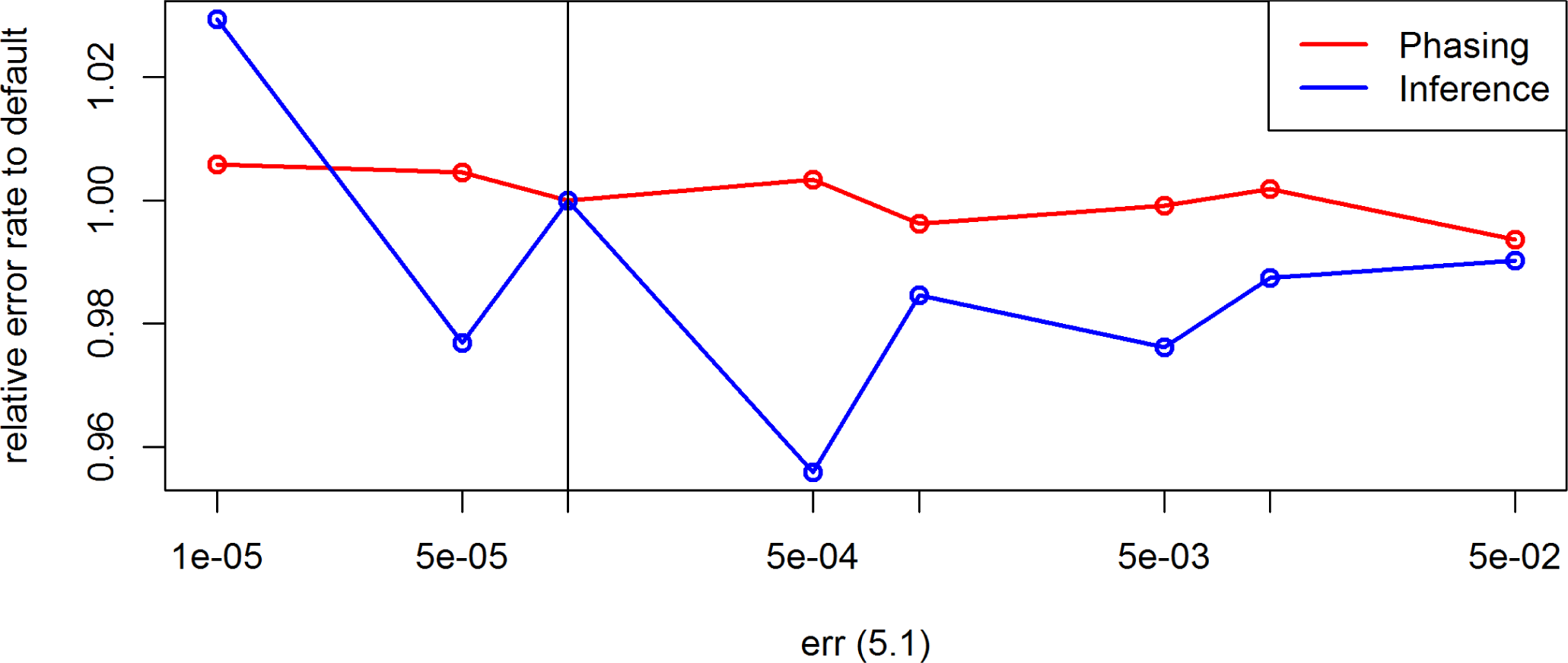
Effect of the parameter *err* on the inference and phasing error rates for chromosome 10 of 250 Pseudo *S*_0_ generated based on the maize data in BEAGLE 5.1. Default settings are indicated by the vertical line.

**Figure S29.**
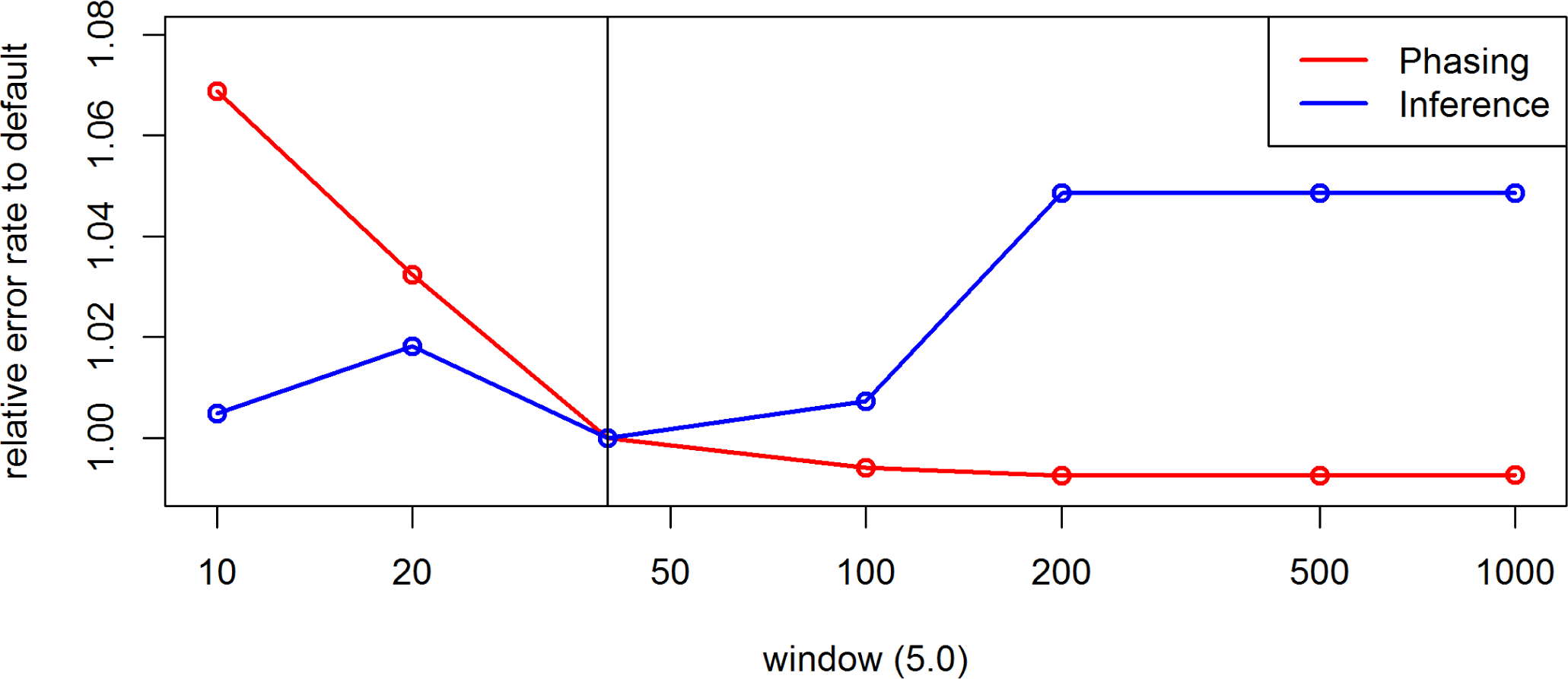
Effect of the parameter *window* on the inference and phasing error rates for chromosome 10 of 250 Pseudo *S*_0_ generated based on the maize data in BEAGLE 5.0. Default settings are indicated by the vertical line.

**Figure S30.**
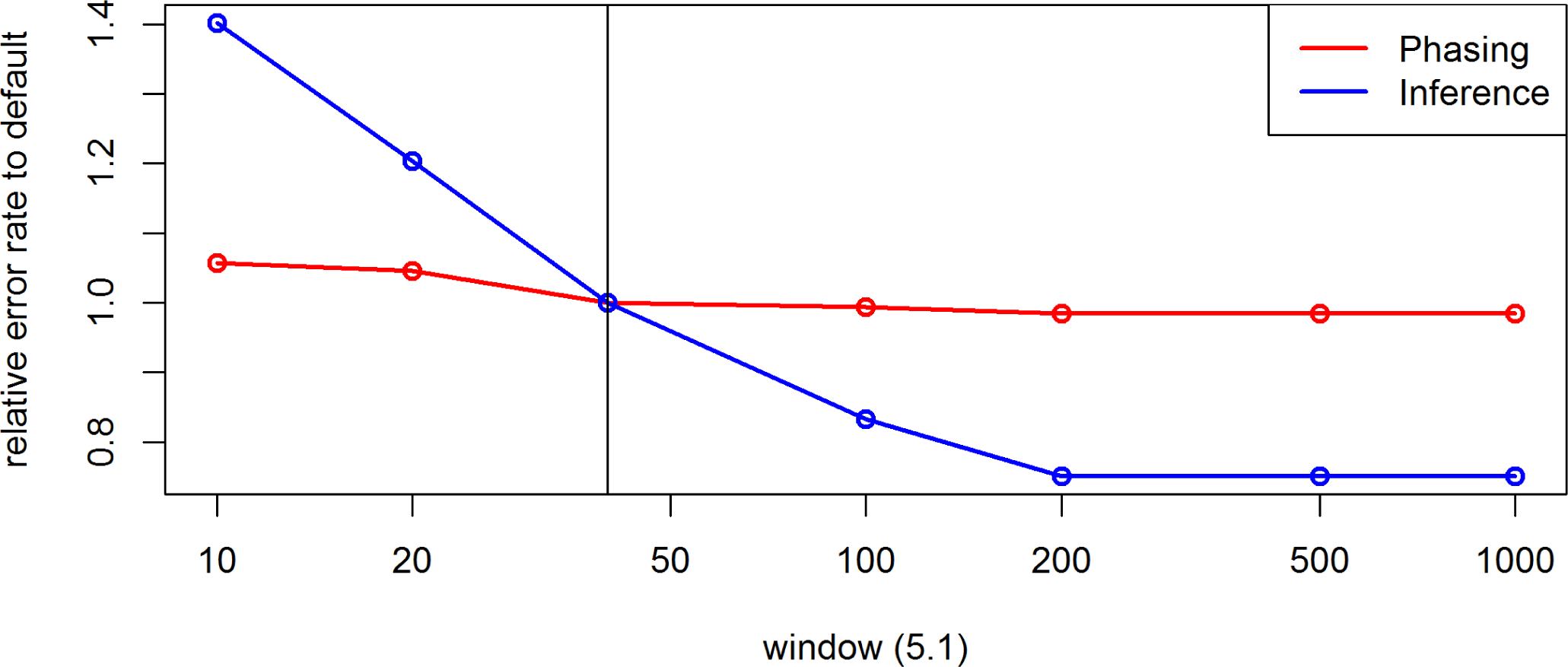
Effect of the parameter *window* on the inference and phasing error rates for chromosome 10 of 250 Pseudo *S*_0_ generated based on the maize data in BEAGLE 5.1. Default settings are indicated by the vertical line.

**Figure S31.**
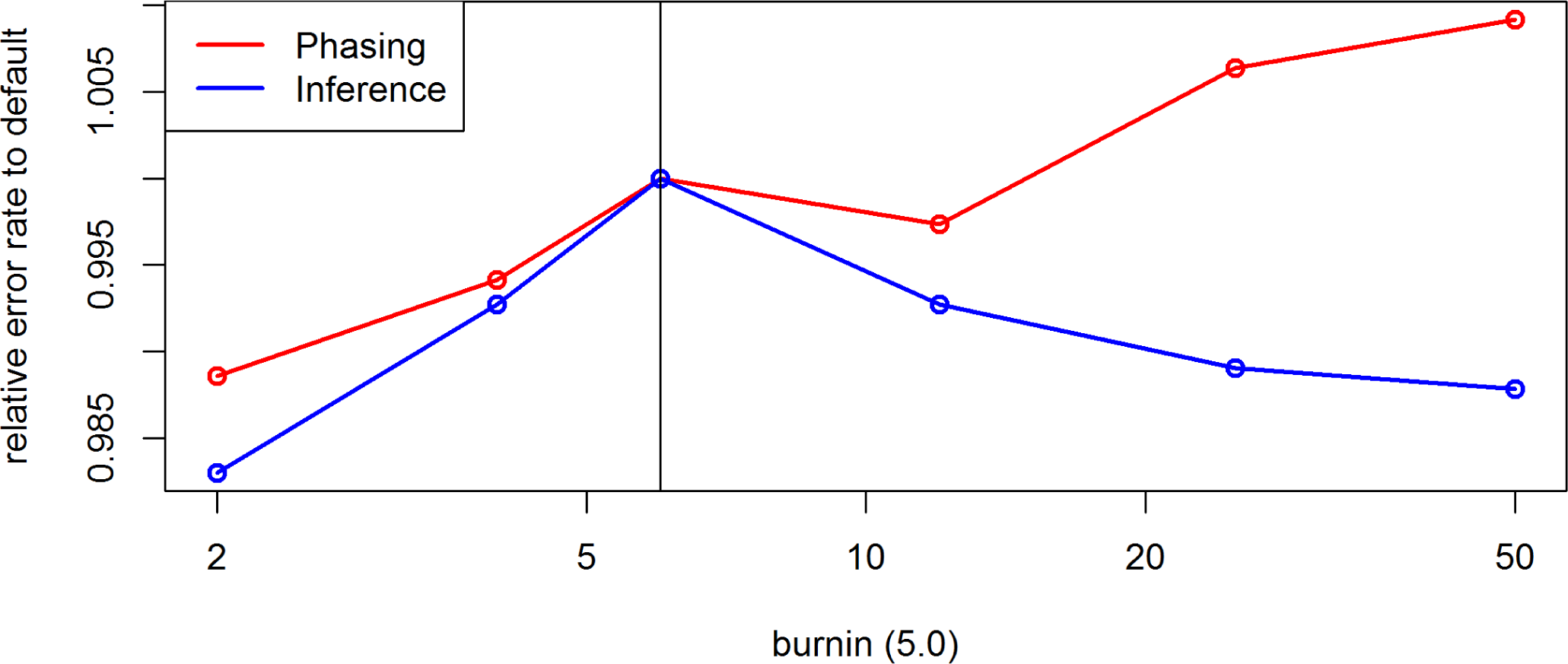
Effect of the parameter *burnin* on the inference and phasing error rates for chromosome 10 of 250 Pseudo *S*_0_ generated based on the maize data in BEAGLE 5.0. Default settings are indicated by the vertical line.

**Figure S32.**
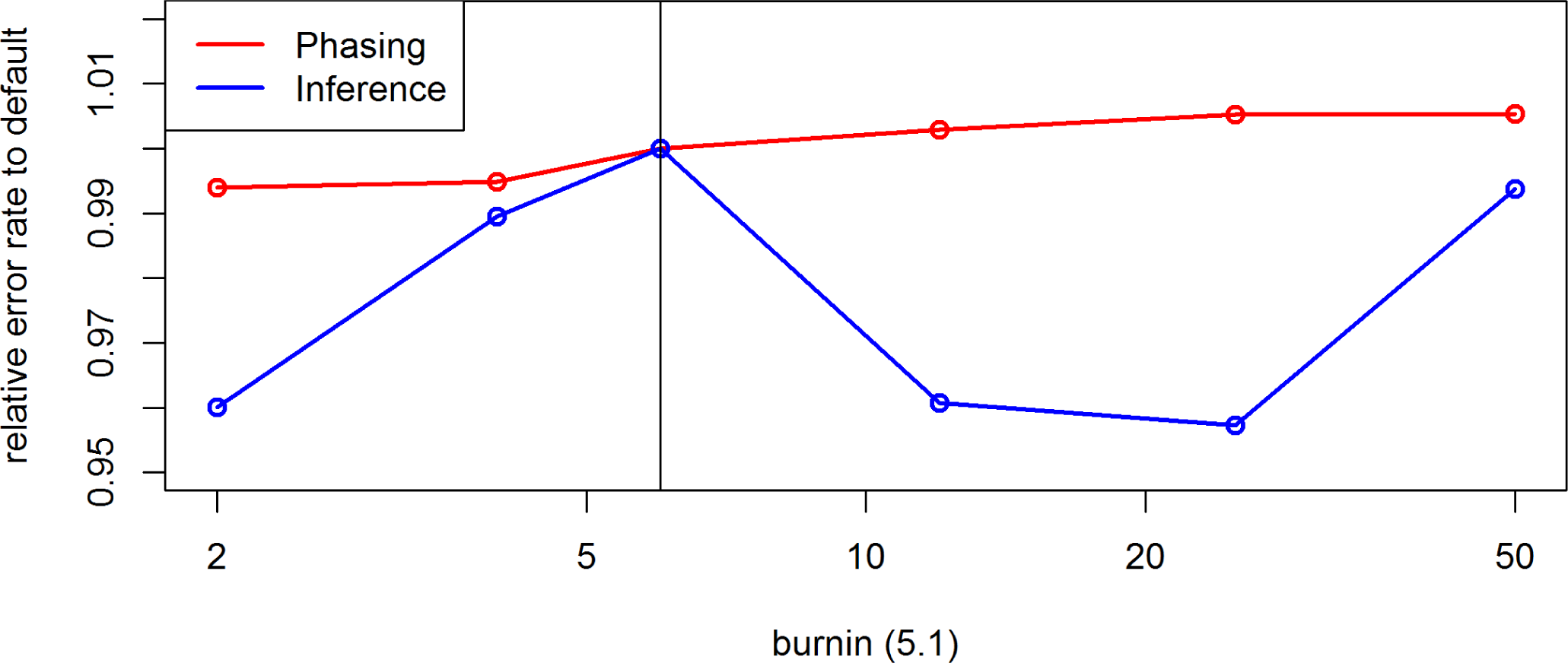
Effect of the parameter *burnin* on the inference and phasing error rates for chromosome 10 of 250 Pseudo *S*_0_ generated based on the maize data in BEAGLE 5.1. Default settings are indicated by the vertical line.

**Figure S33.**
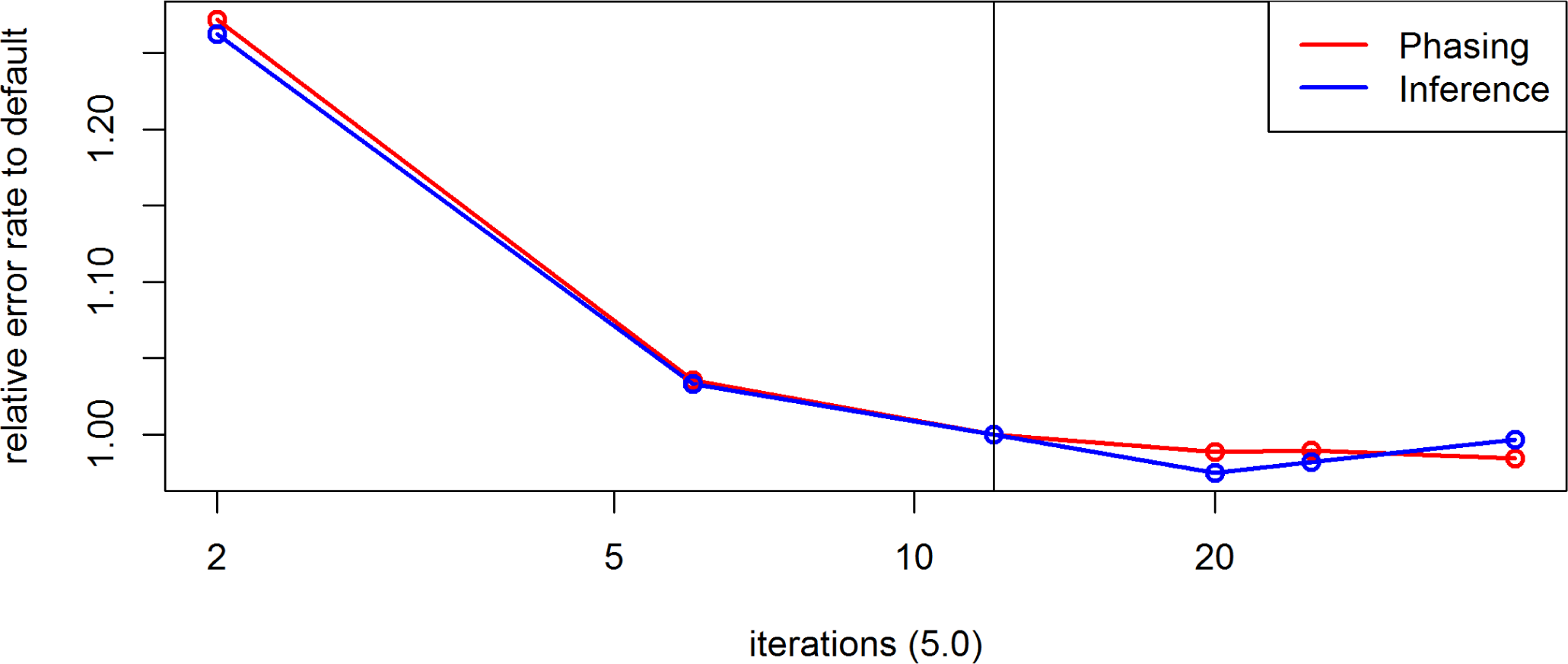
Effect of the parameter *iterations* on the inference and phasing error rates for chromosome 10 of 250 Pseudo *S*_0_ generated based on the maize data in BEAGLE 5.0. Default settings are indicated by the vertical line.

**Figure S34.**
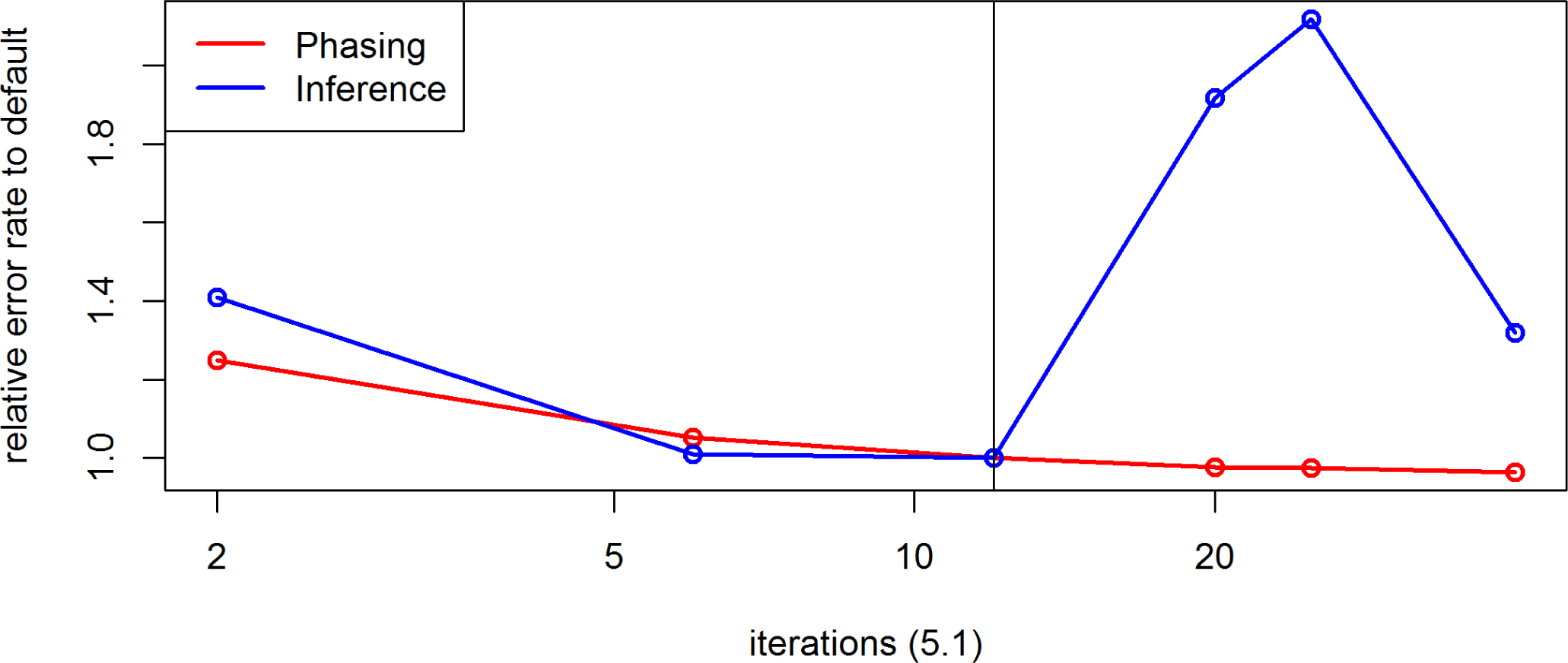
Effect of the parameter *iterations* on the inference and phasing error rates for chromosome 10 of 250 Pseudo *S*_0_ generated based on the maize data in BEAGLE 5.1. Default settings are indicated by the vertical line.

**Figure S35.**
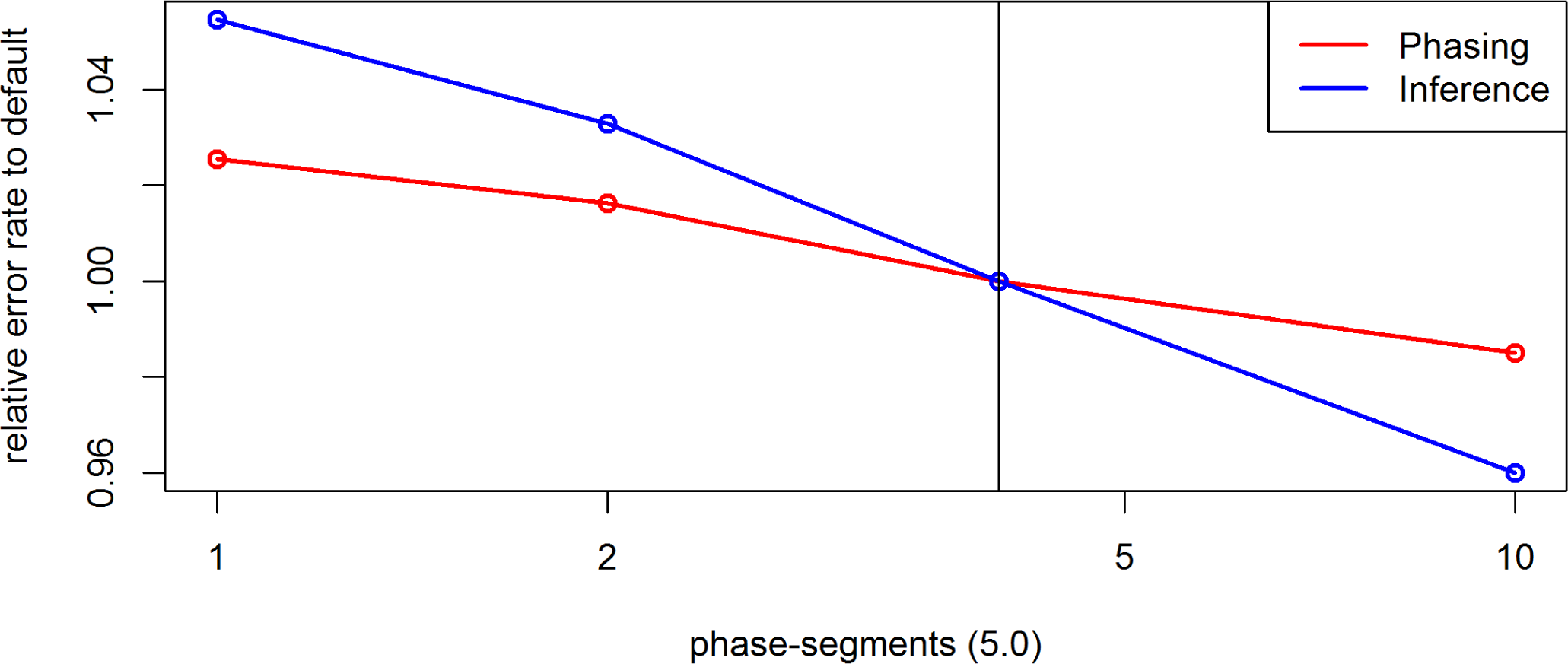
Effect of the parameter *phase-segment* on the inference and phasing error rates for chromosome 10 of 250 Pseudo *S*_0_ generated based on the maize data in BEAGLE 5.0. Default settings are indicated by the vertical line.

**Figure S36.**
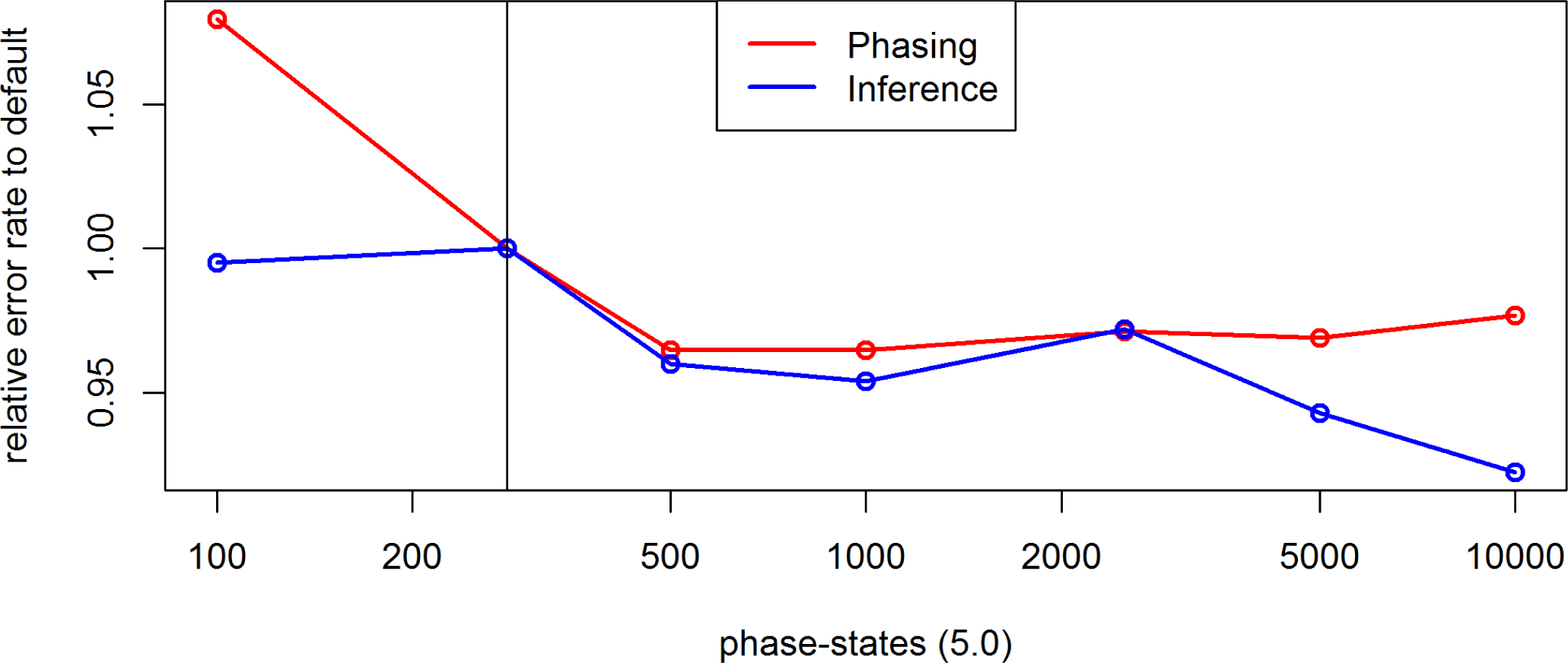
Effect of the parameter *phase-states* on the inference and phasing error rates for chromosome 10 of 250 Pseudo *S*_0_ generated based on the maize data in BEAGLE 5.0. Default settings are indicated by the vertical line.

**Figure S37.**
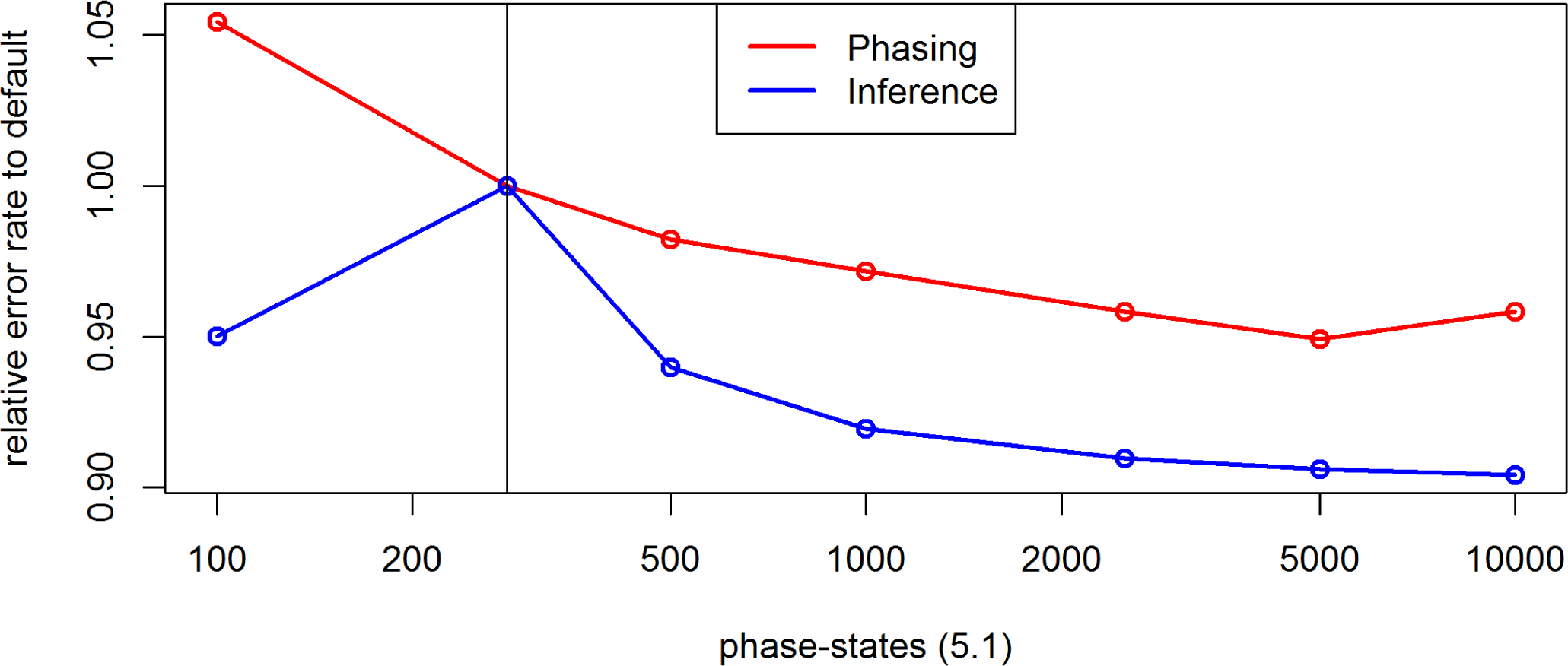
Effect of the parameter *phase-states* on the inference and phasing error rates for chromosome 10 of 250 Pseudo *S*_0_ generated based on the maize data in BEAGLE 5.1. Default settings are indicated by the vertical line.

**Figure S38.**
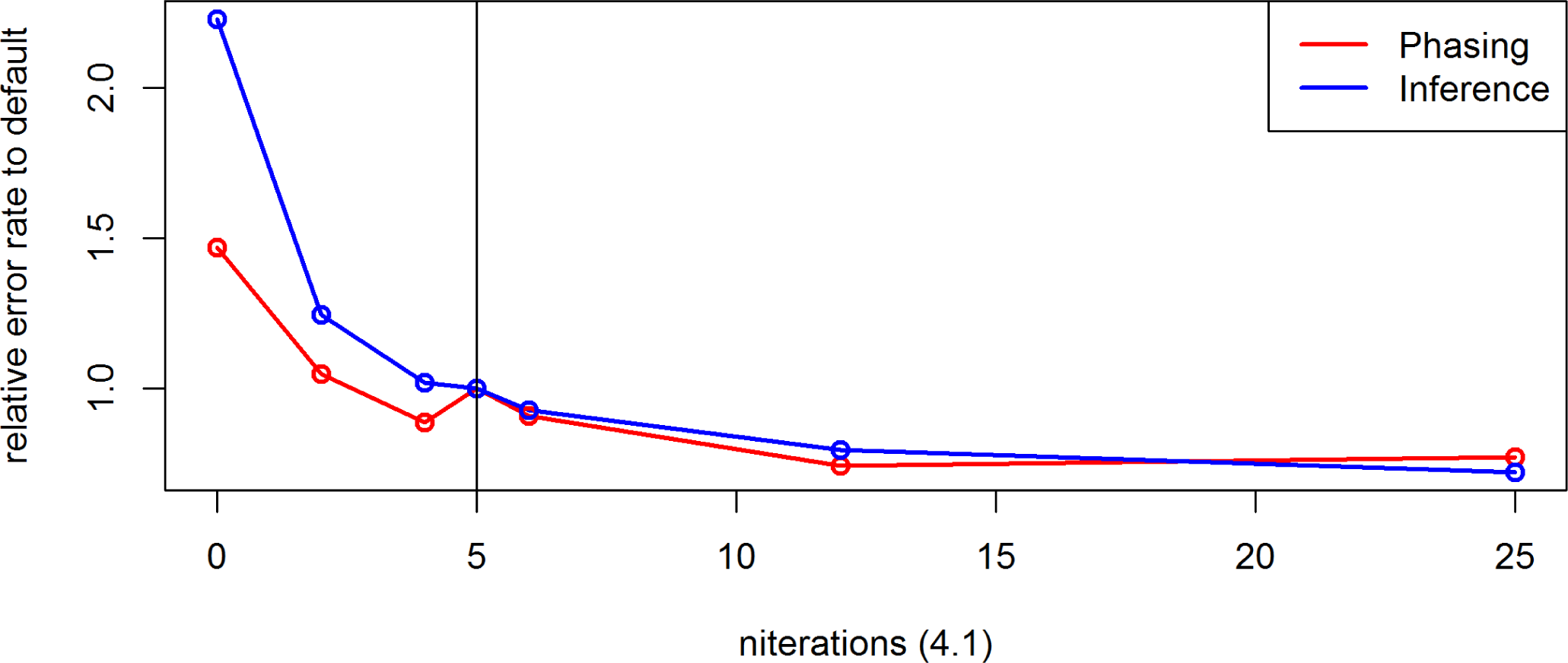
Effect of the parameter *niterations* on the inference and phasing error rates for chromosome 10 of 250 Pseudo *S*_0_ generated based on the maize data in BEAGLE 4.1. Default settings are indicated by the vertical line.

**Figure S39.**
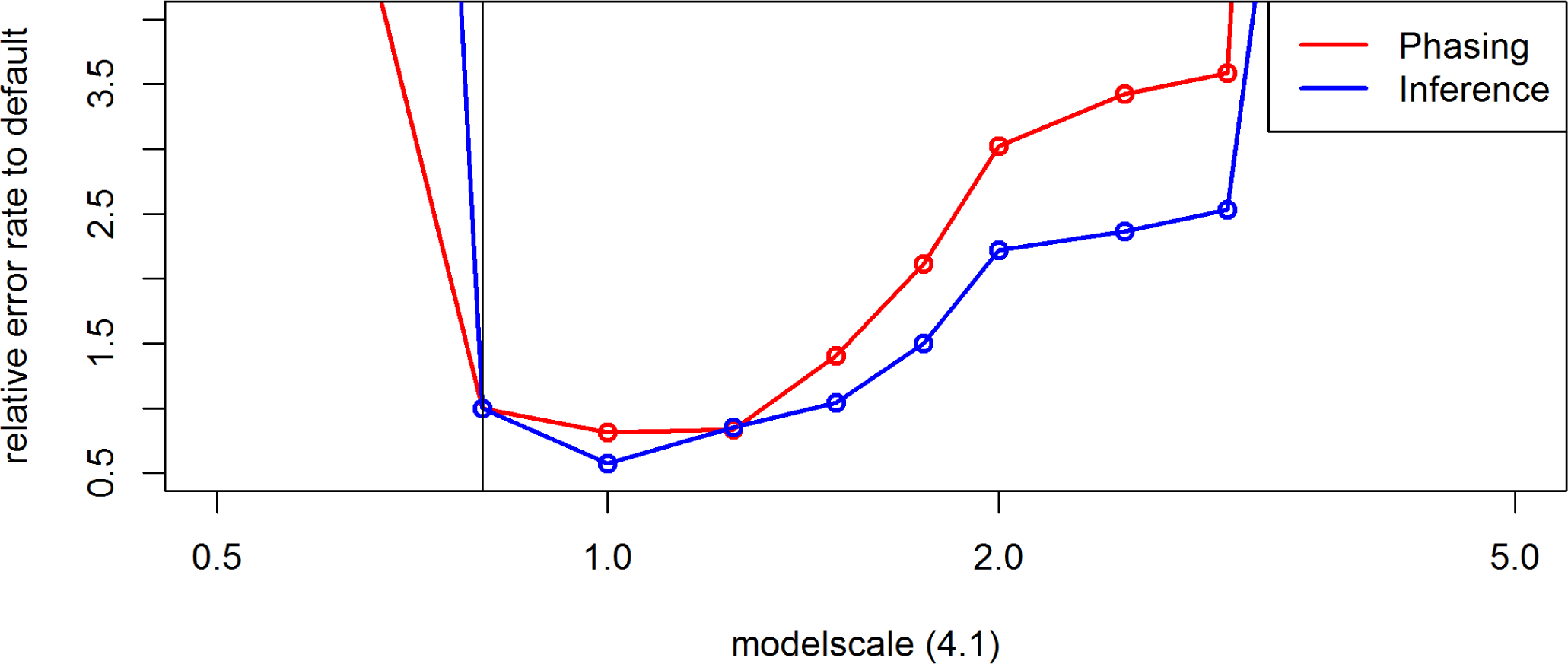
Effect of the parameter *modelscale* on the inference and phasing error rates for chromosome 10 of 250 Pseudo *S*_0_ generated based on the maize data in BEAGLE 4.1. Default settings are indicated by the vertical line.

**Figure S40.**
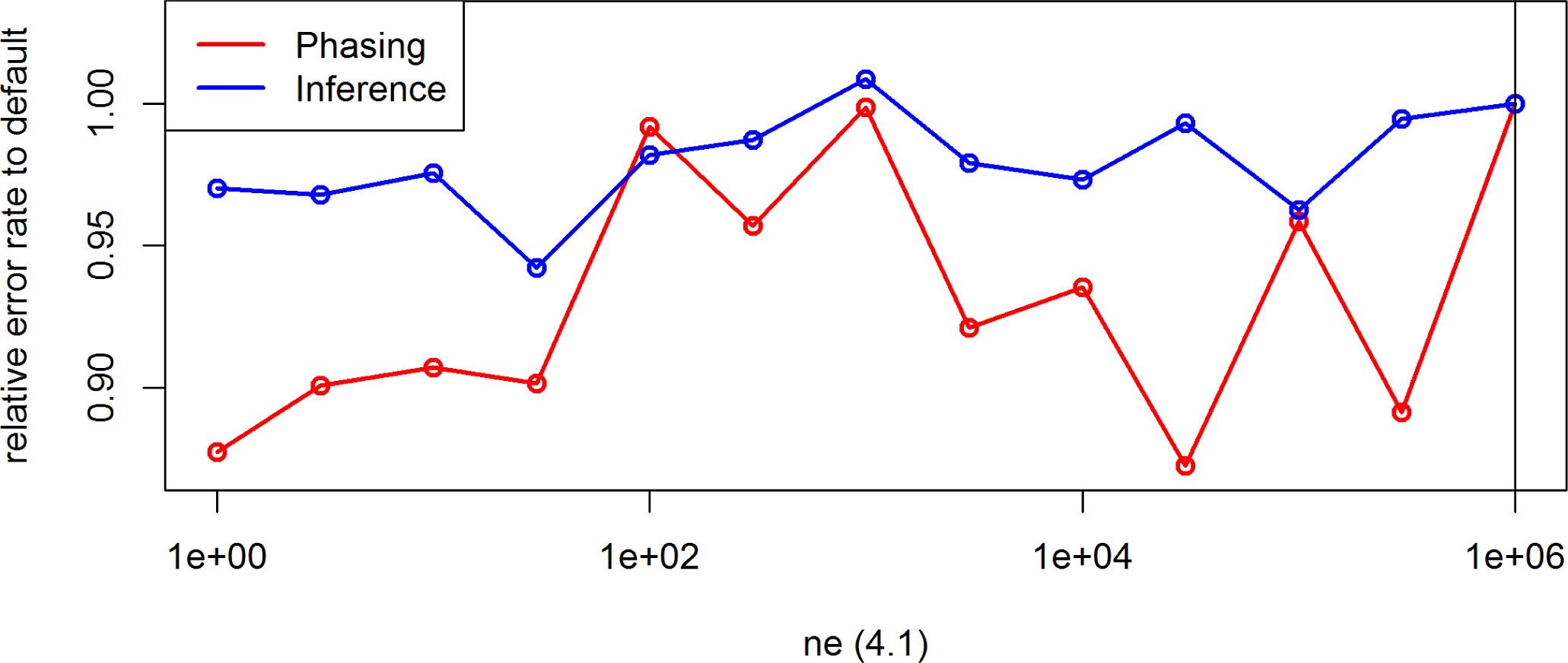
Effect of the parameter *ne* on the inference and phasing error rates for chromosome 10 of 250 Pseudo *S*_0_ generated based on the maize data in BEAGLE 4.1. Default settings are indicated by the vertical line.

**Figure S41.**
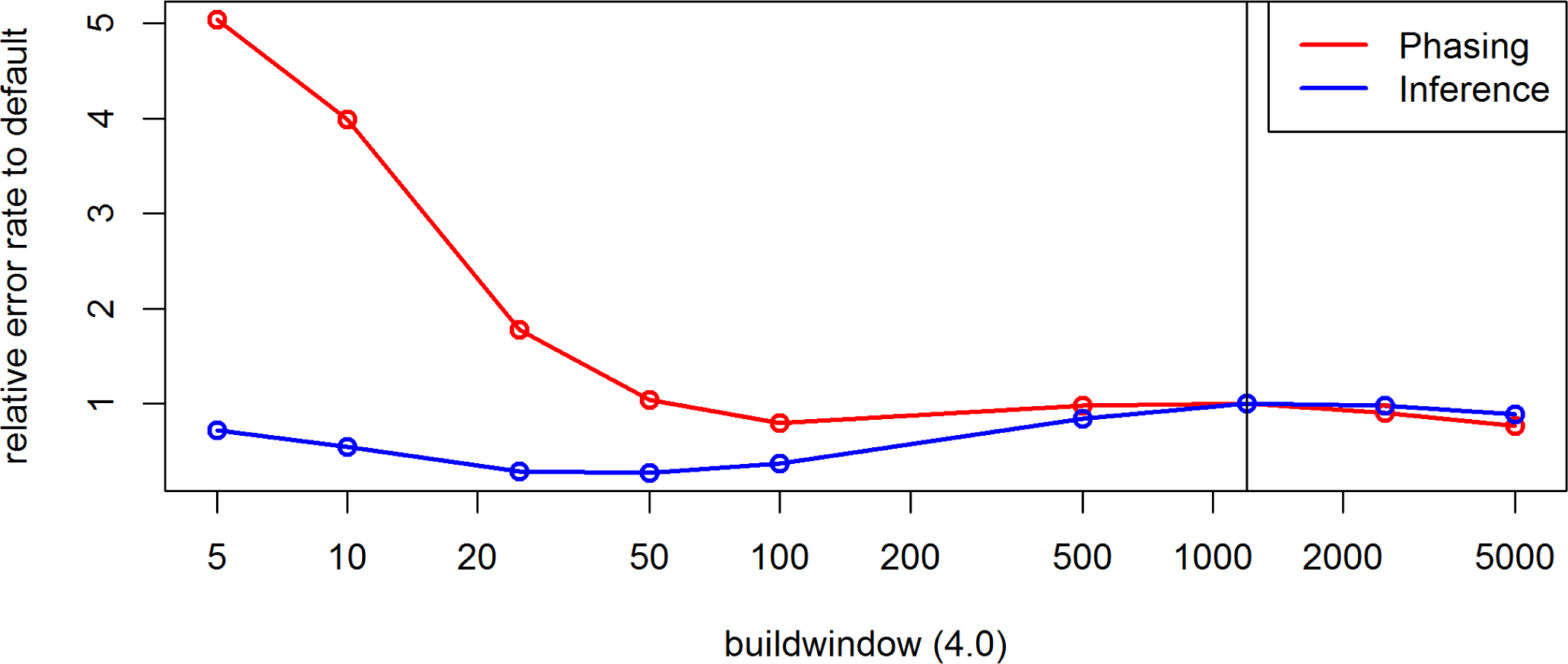
Effect of the parameter *buildwindow* on the inference and phasing error rates for chromosome 10 of 250 Pseudo *S*_0_ generated based on the maize data in BEAGLE 4.0. Default settings are indicated by the vertical line.

**Figure S42.**
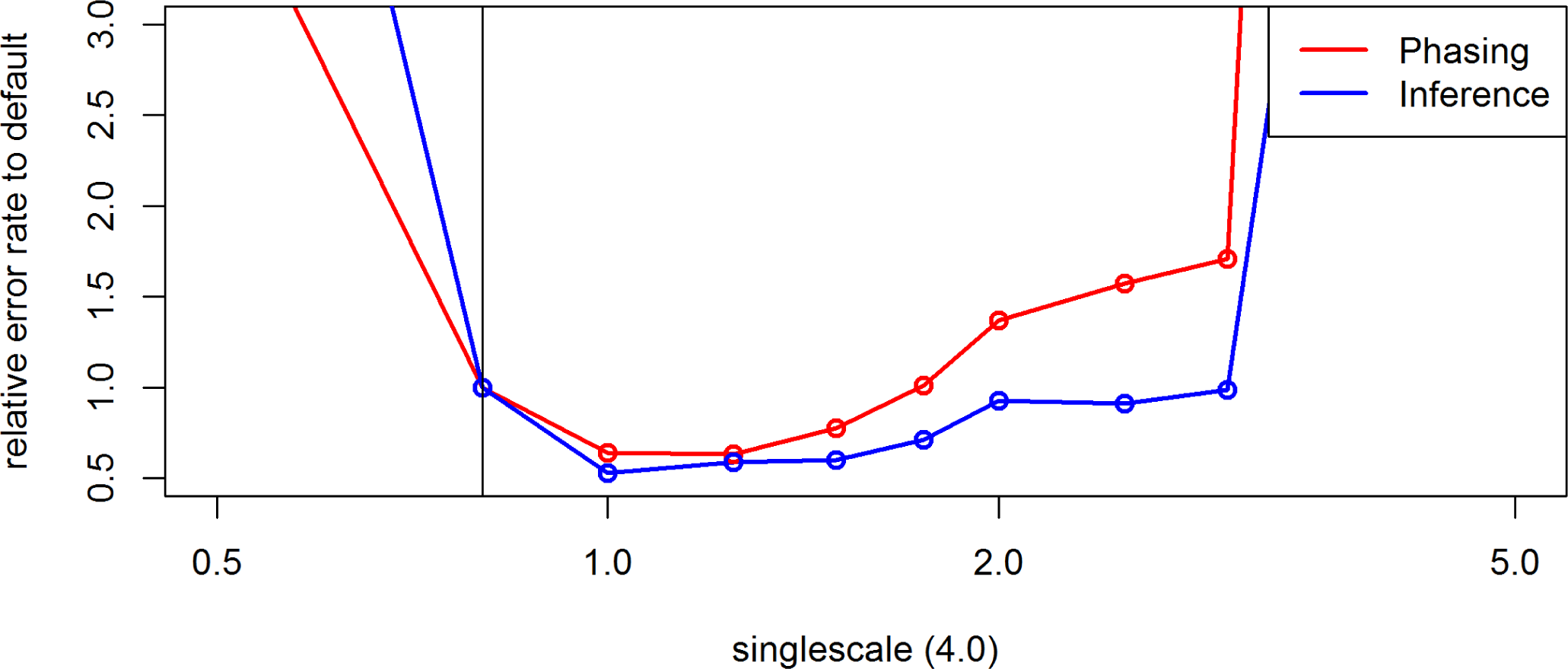
Effect of the parameter *singlescale* on the inference and phasing error rates for chromosome 10 of 250 Pseudo *S*_0_ generated based on the maize data in BEAGLE 4.0. Default settings are indicated by the vertical line.

**Figure S43.**
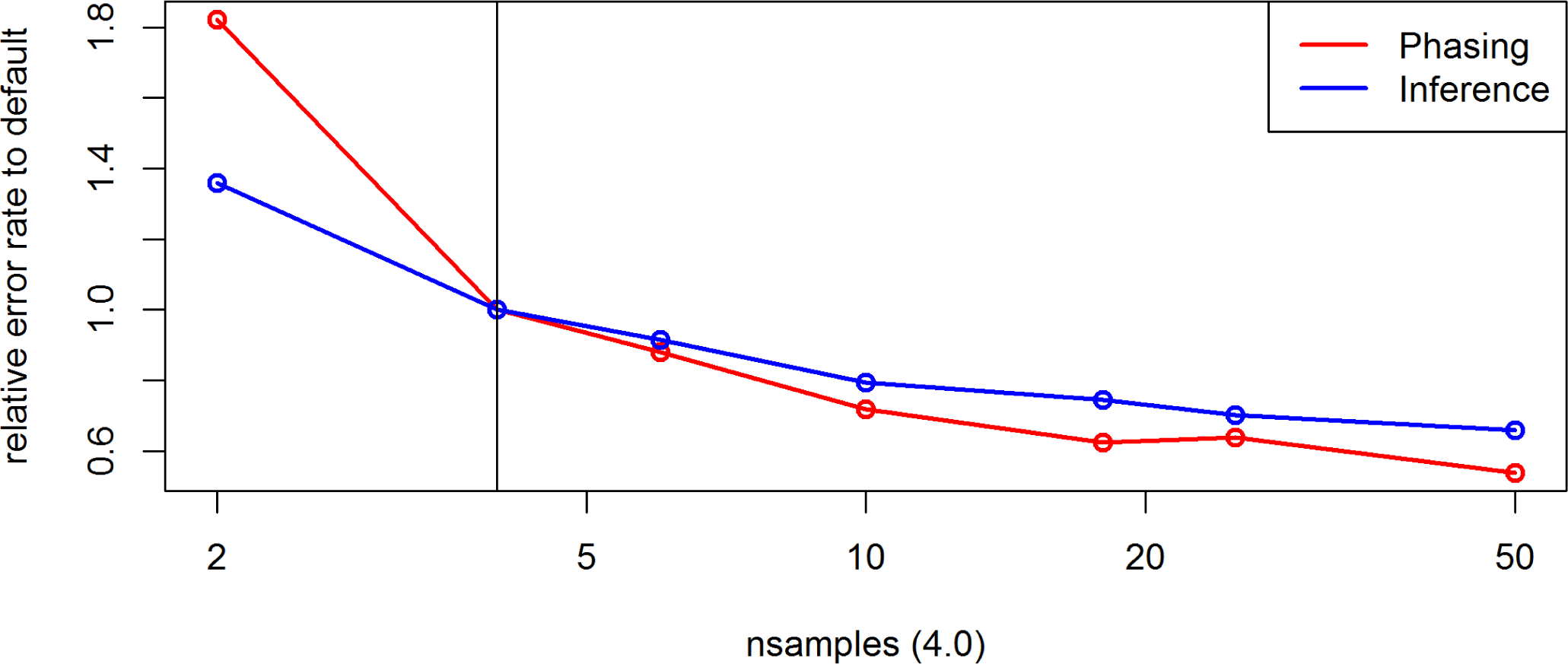
Effect of the parameter *nsamples* on the inference and phasing error rates for chromosome 10 of 250 Pseudo *S*_0_ generated based on the maize data in BEAGLE 4.0. Default settings are indicated by the vertical line.

**Figure S44.**
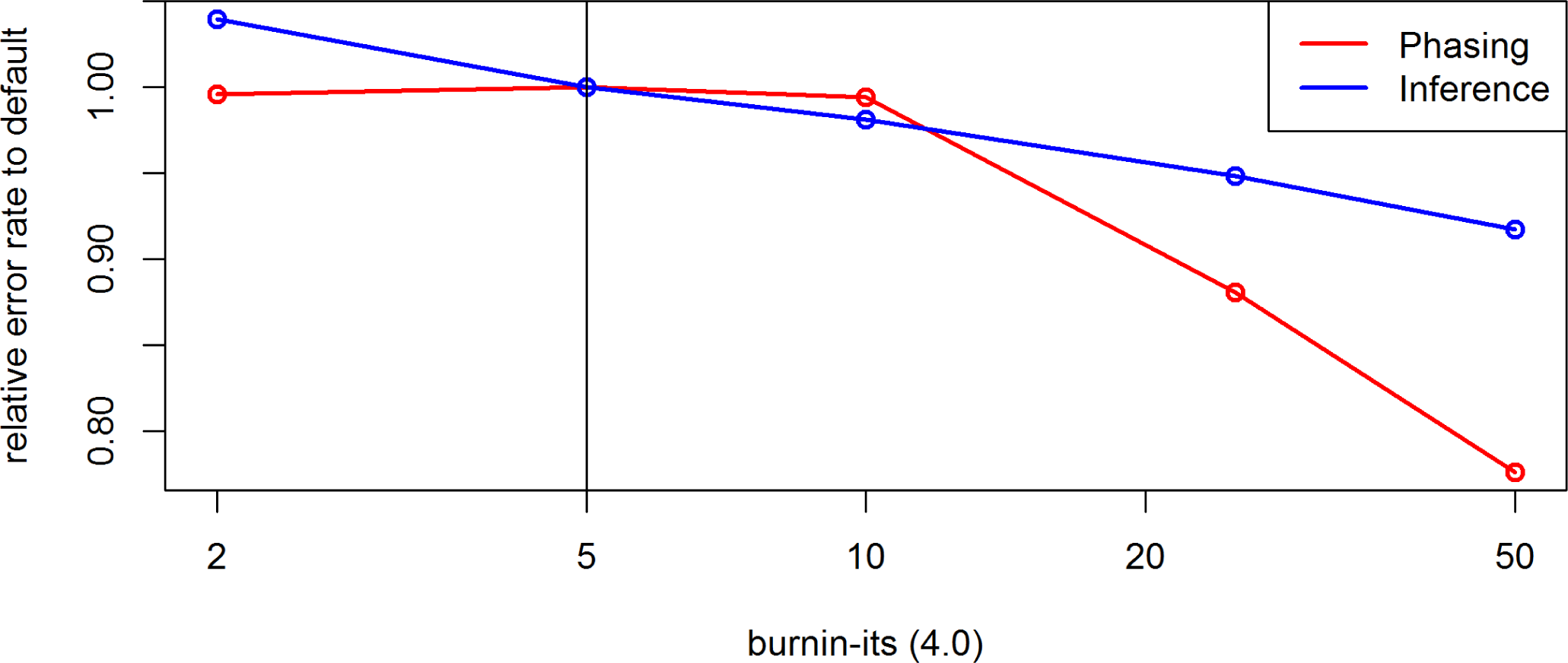
Effect of the parameter *burnin-its* on the inference and phasing error rates for chromosome 10 of 250 Pseudo *S*_0_ generated based on the maize data in BEAGLE 4.0. Default settings are indicated by the vertical line.

**Figure S45.**
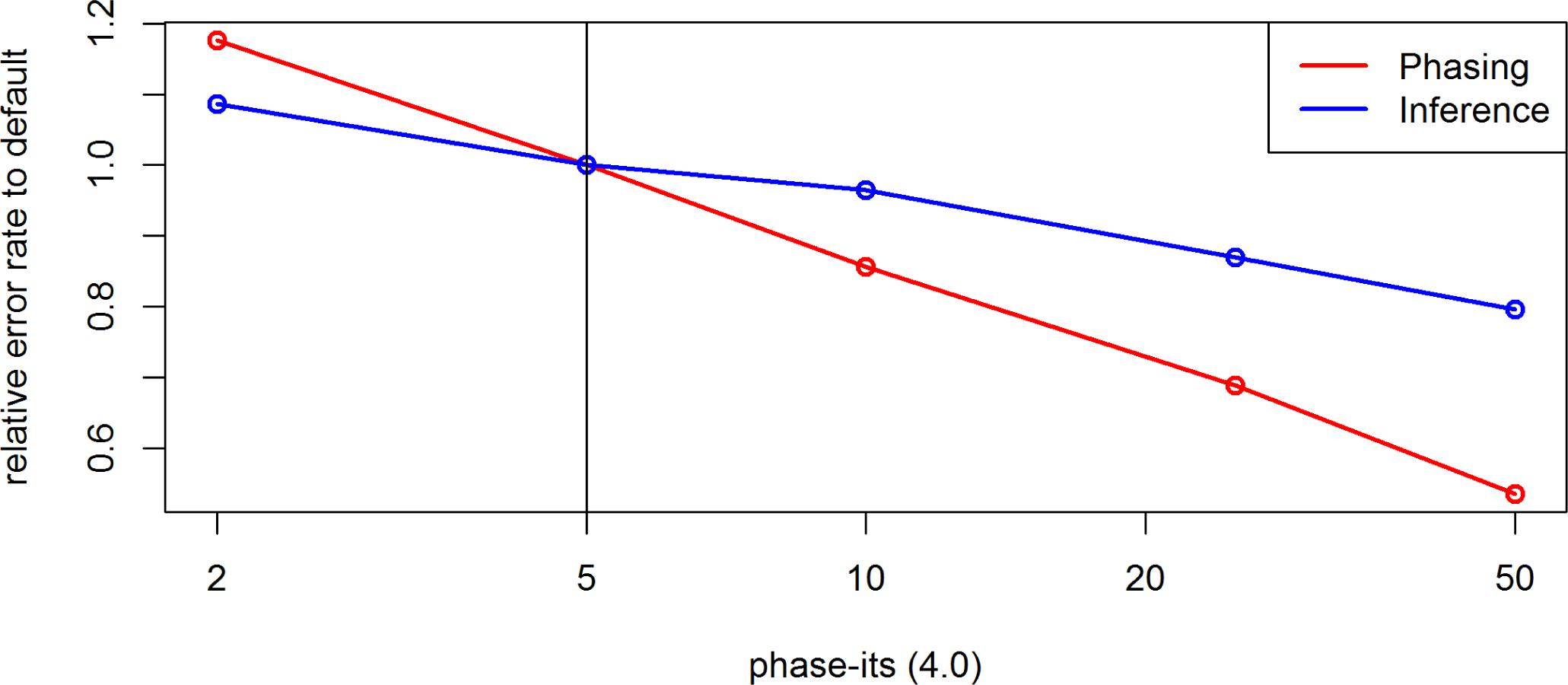
Effect of the parameter *phase-its* on the inference and phasing error rates for chromosome 10 of 250 Pseudo *S*_0_ generated based on the maize data in BEAGLE 4.0. Default settings are indicated by the vertical line.

**Figure S46.**
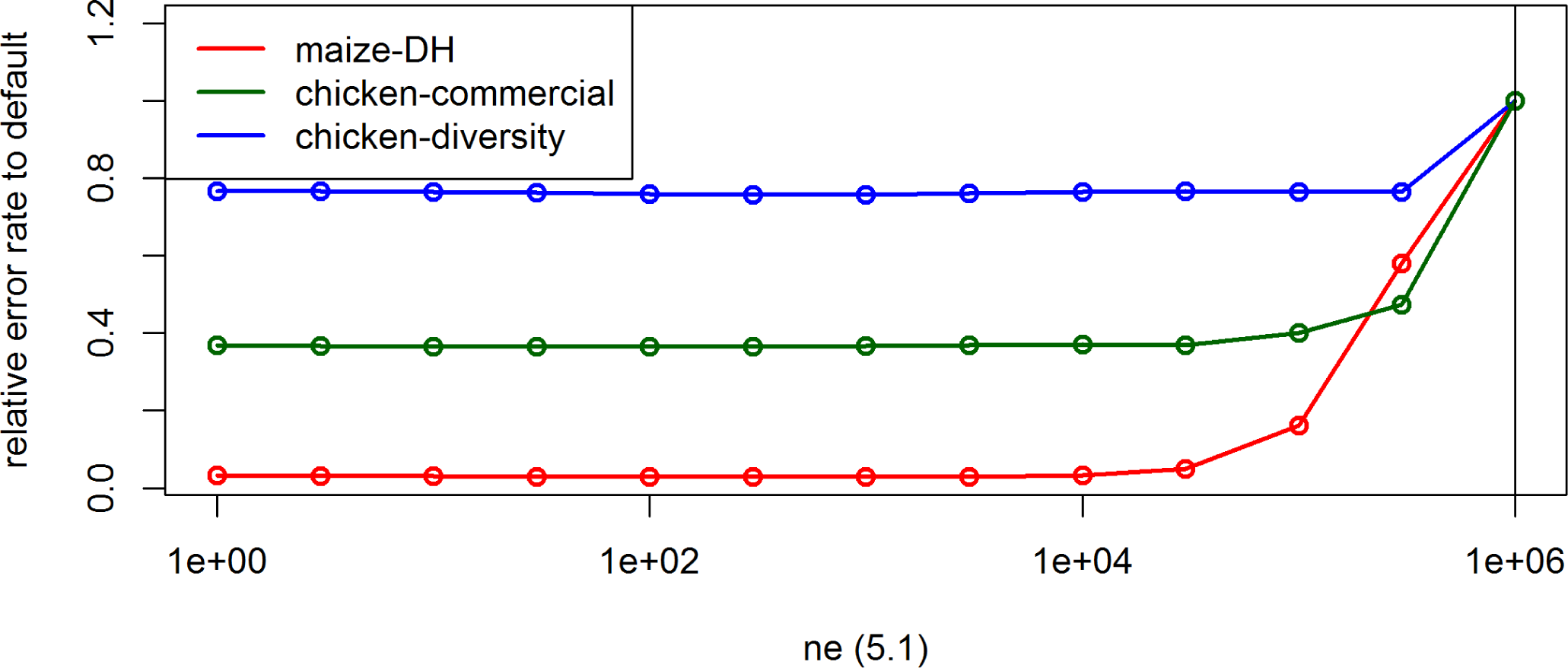
Effect of the parameter *ne* on the UM imputation error rate for the maize data, the commercial chicken line and the chicken diversity panel in BEAGLE 5.1. Default settings are indicated by the vertical line.

**Figure S47.**
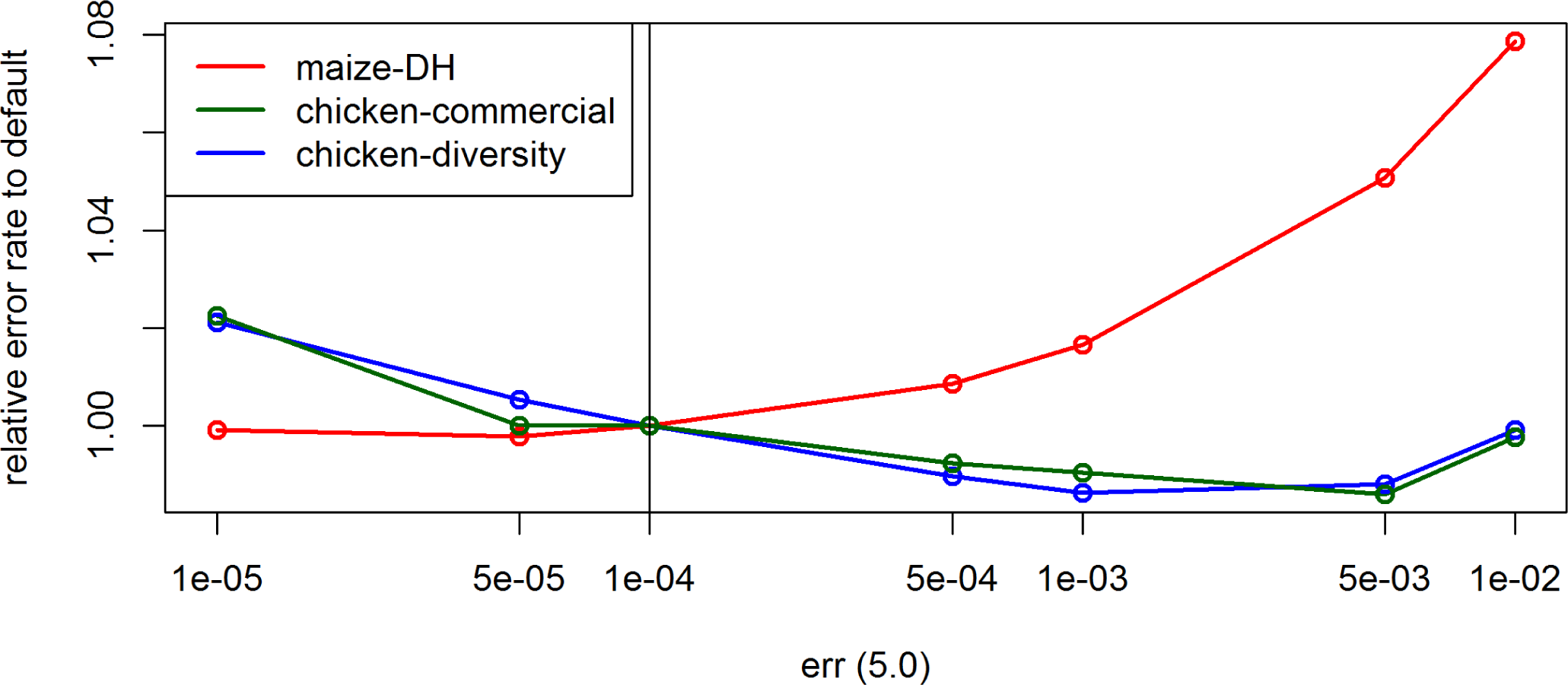
Effect of the parameter *err* on the UM imputation error rate for the maize data, the commercial chicken line and the chicken diversity panel in BEAGLE 5.0. Default settings are indicated by the vertical line.

**Figure S48.**
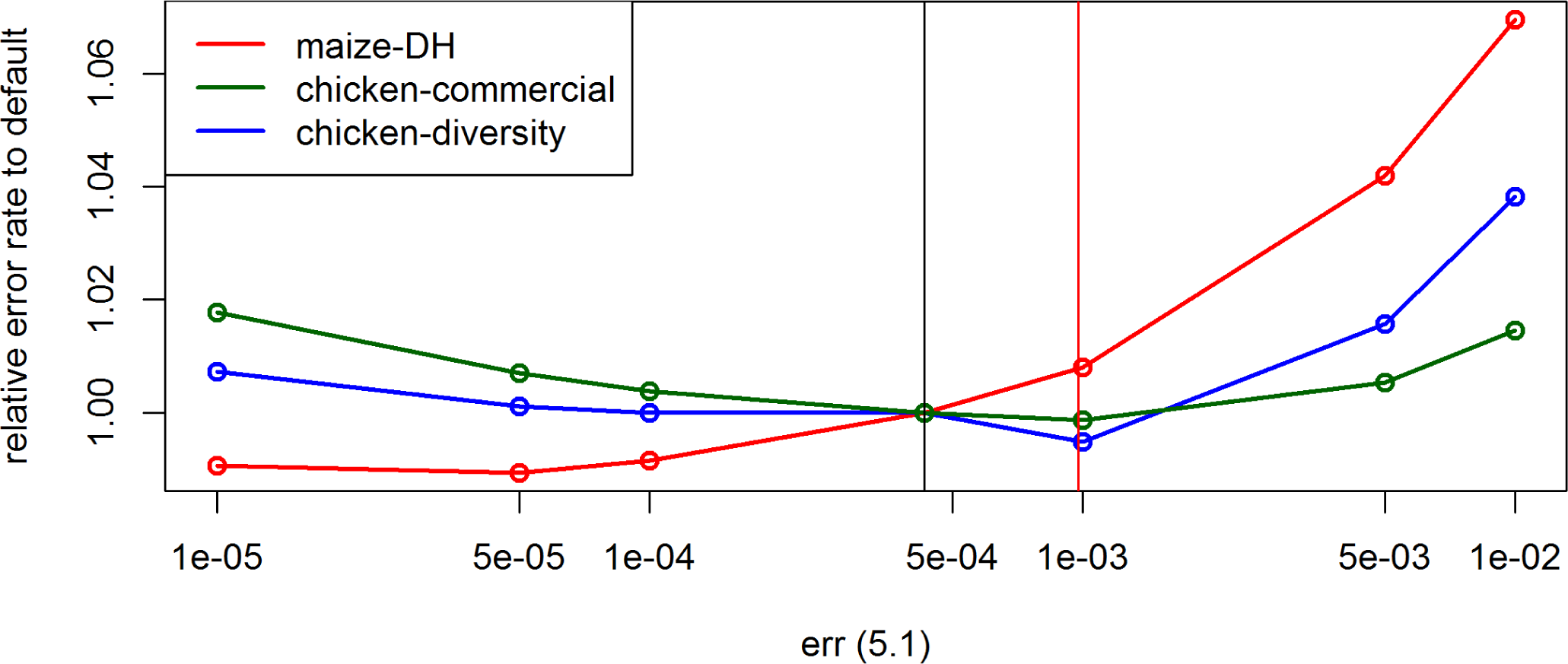
Effect of the parameter *err* on the UM imputation error rate for the maize data, the commercial chicken line and the chicken diversity panel in BEAGLE 5.1. Default settings are indicated by the vertical lines (black for chicken, red for maize).

**Figure S49.**
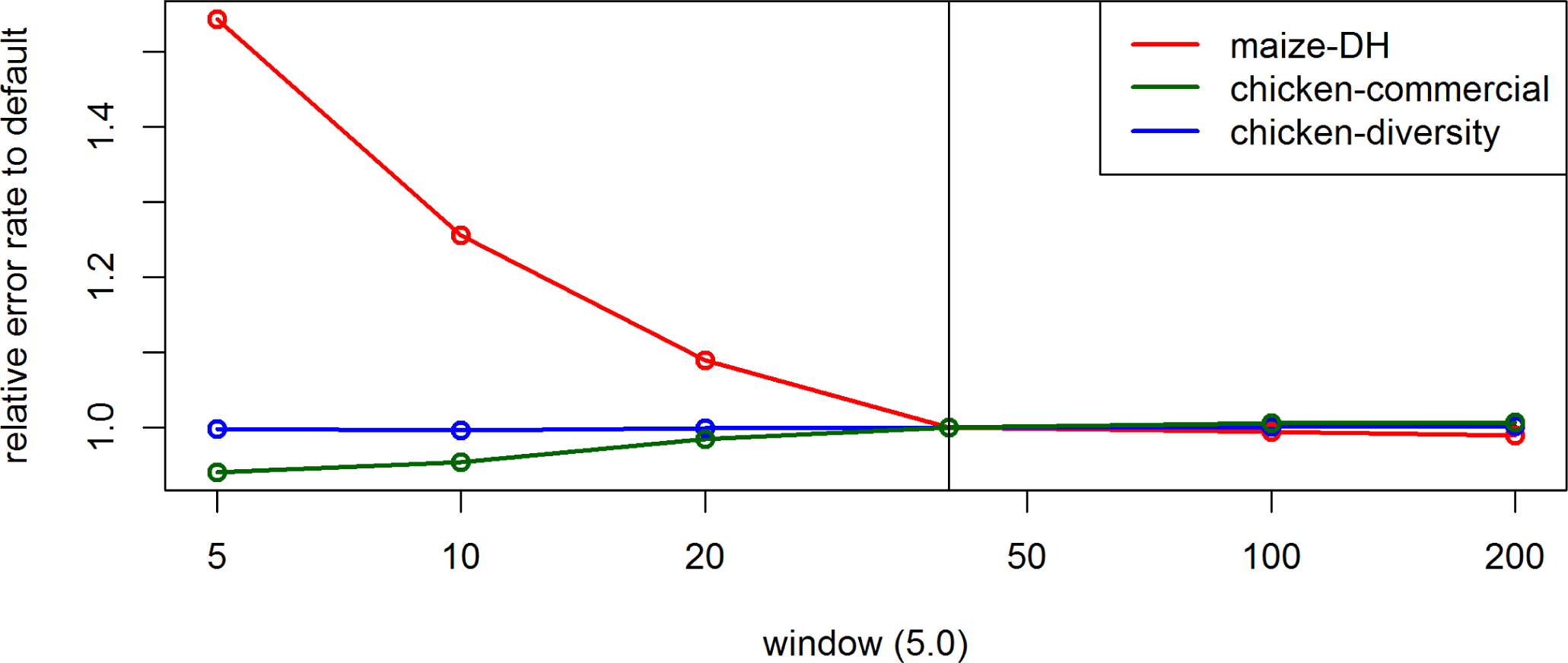
Effect of the parameter *window* on the UM imputation error rate for the maize data, the commercial chicken line and the chicken diversity panel in BEAGLE 5.0. Default settings are indicated by the vertical line.

**Figure S50.**
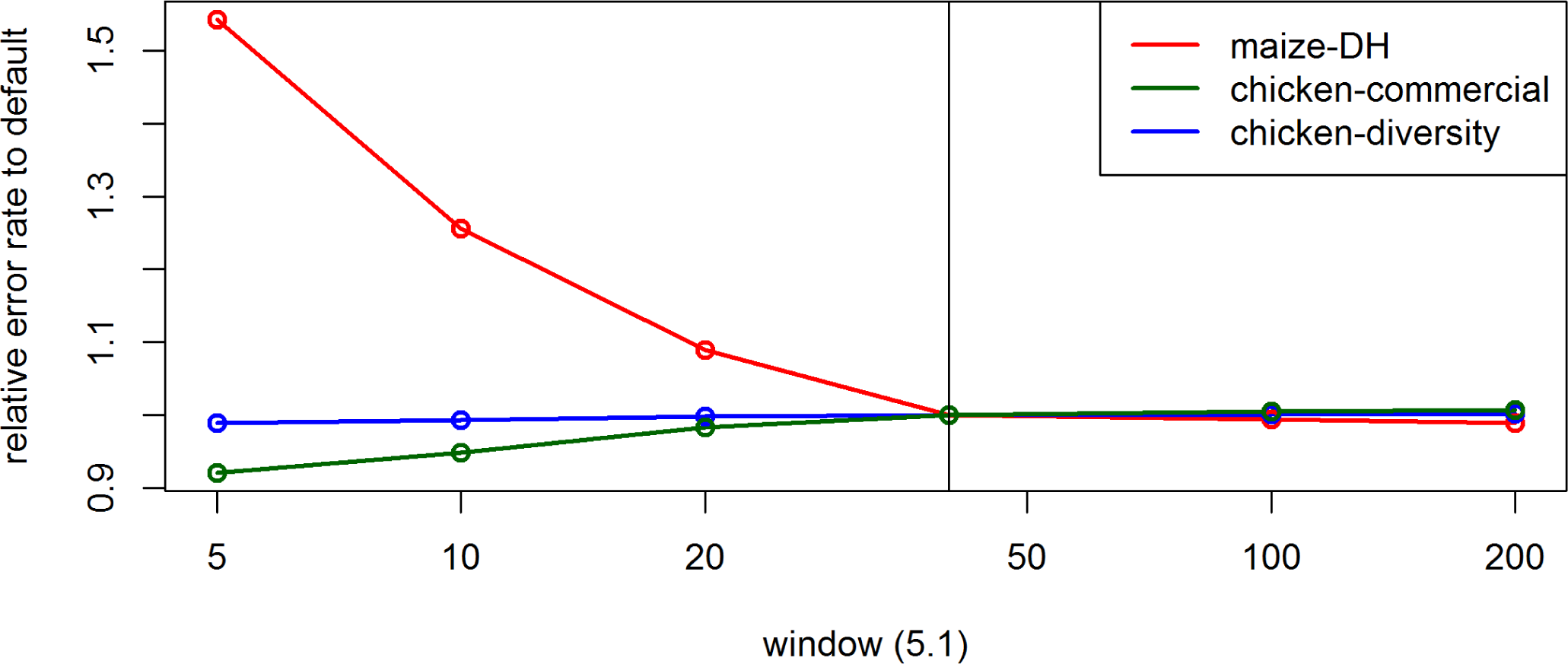
Effect of the parameter *window* on the UM imputation error rate for the maize data, the commercial chicken line and the chicken diversity panel in BEAGLE 5.1. Default settings are indicated by the vertical line.

**Figure S51.**
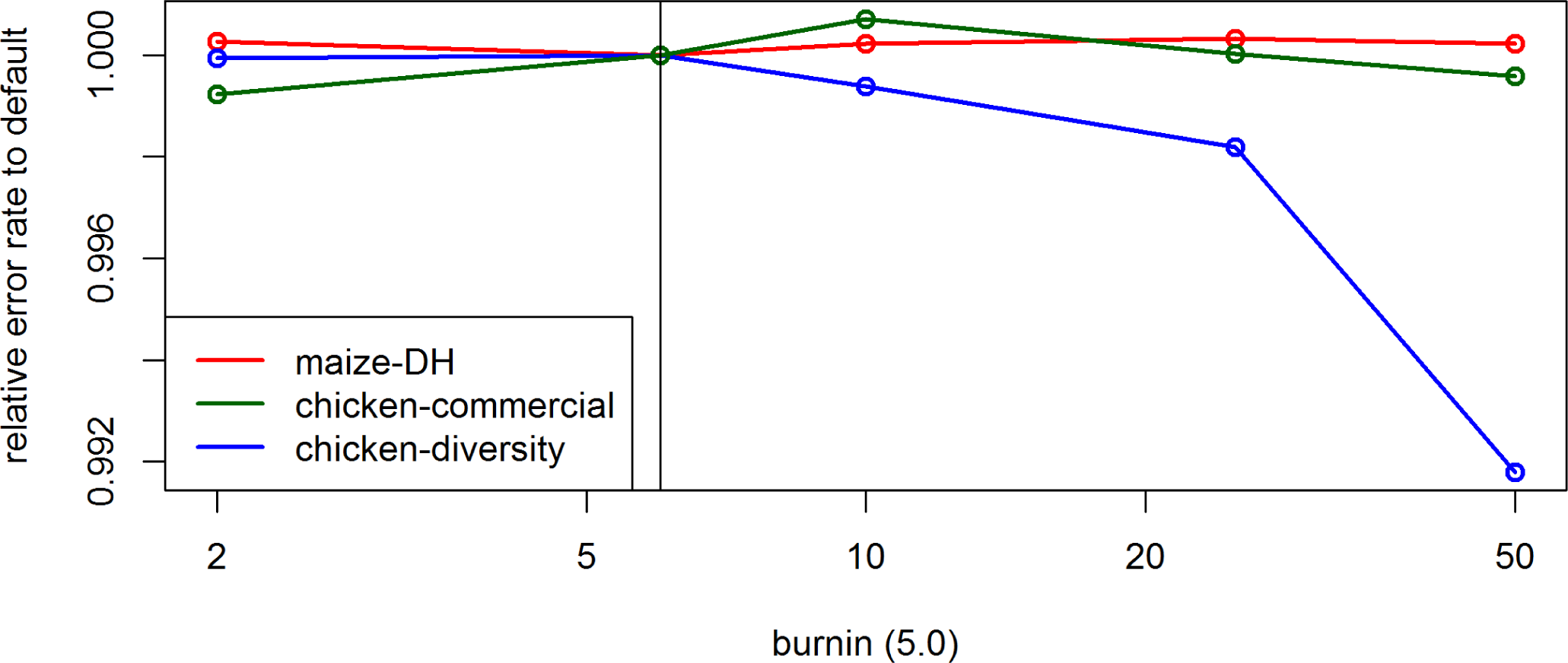
Effect of the parameter *burnin* on the UM imputation error rate for the maize data, the commercial chicken line and the chicken diversity panel in BEAGLE 5.0. Default settings are indicated by the vertical line.

**Figure S52.**
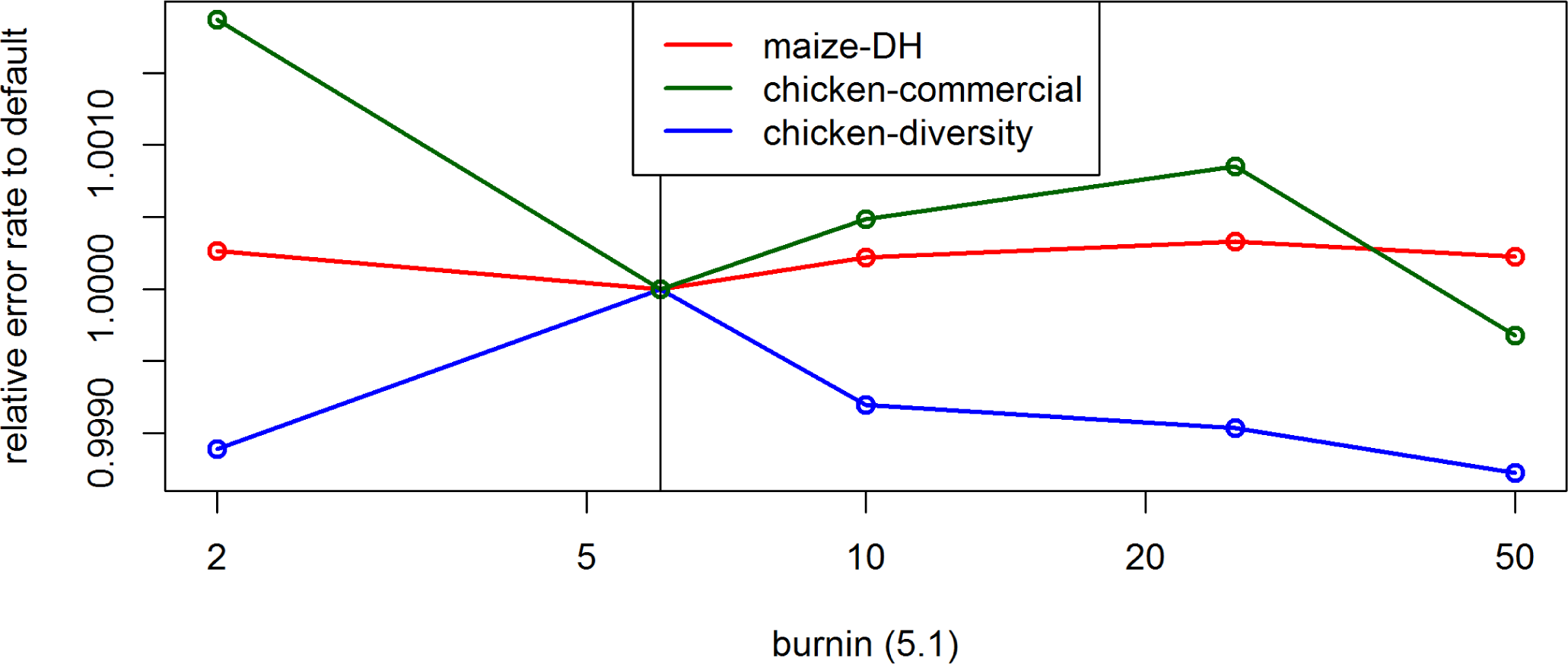
Effect of the parameter *burnin* on the UM imputation error rate for the maize data, the commercial chicken line and the chicken diversity panel in BEAGLE 5.1. Default settings are indicated by the vertical line.

**Figure S53.**
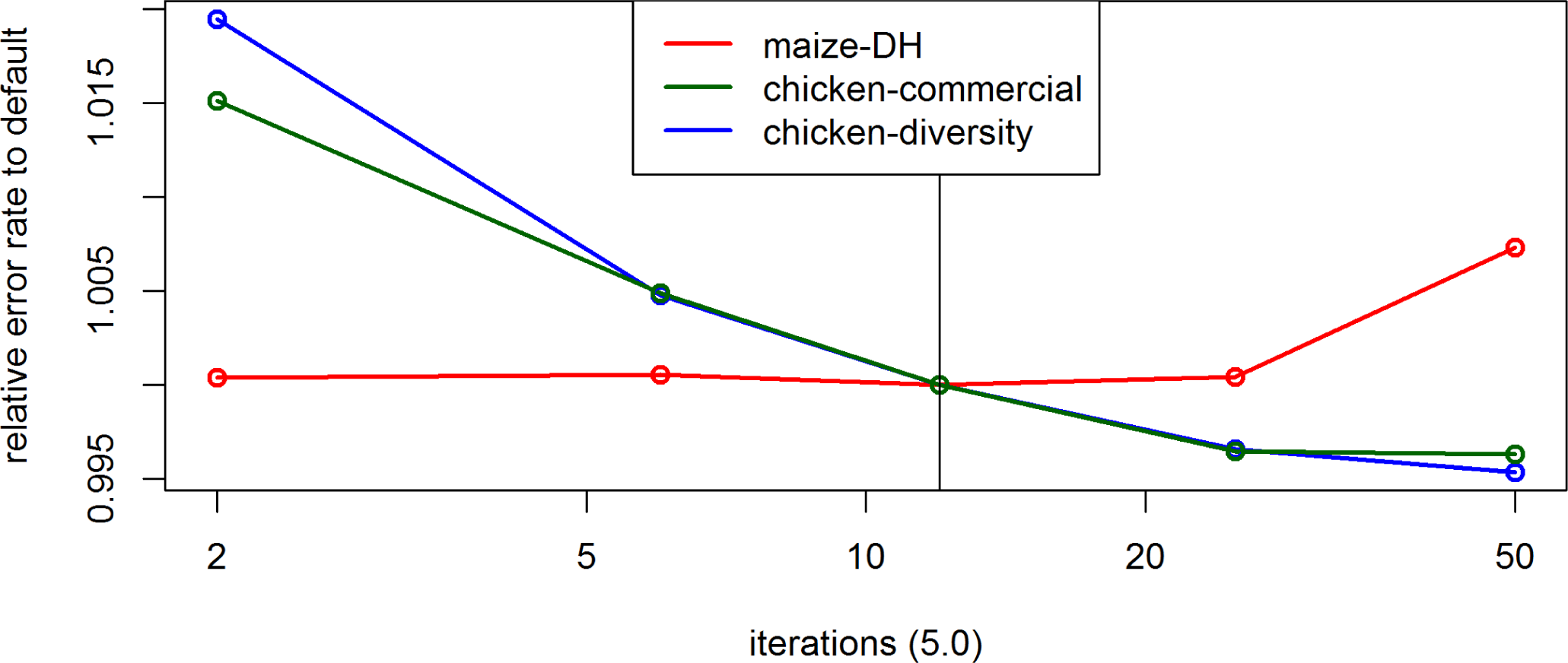
Effect of the parameter *iterations* on the UM imputation error rate for the maize data, the commercial chicken line and the chicken diversity panel in BEAGLE 5.0. Default settings are indicated by the vertical line.

**Figure S54.**
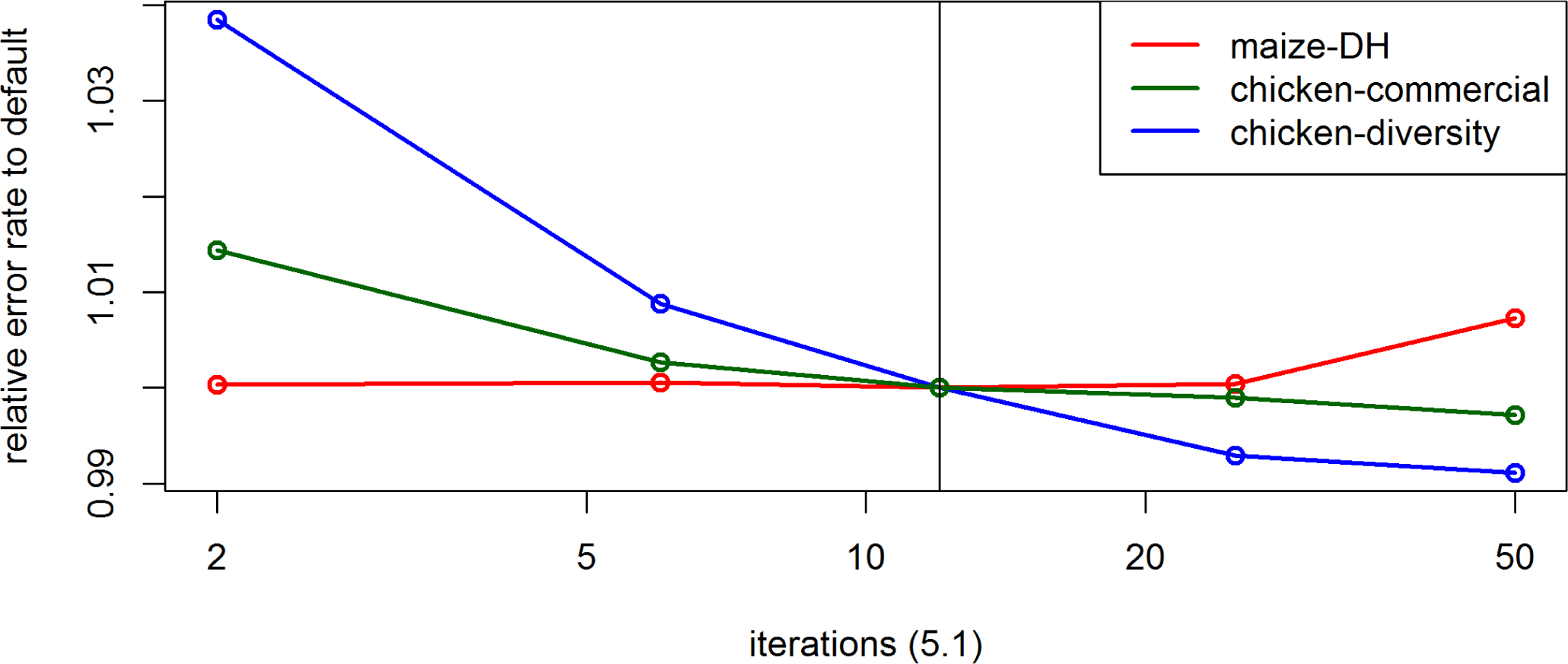
Effect of the parameter *iterations* on the UM imputation error rate for the maize data, the commercial chicken line and the chicken diversity panel in BEAGLE 5.1. Default settings are indicated by the vertical line.

**Figure S55.**
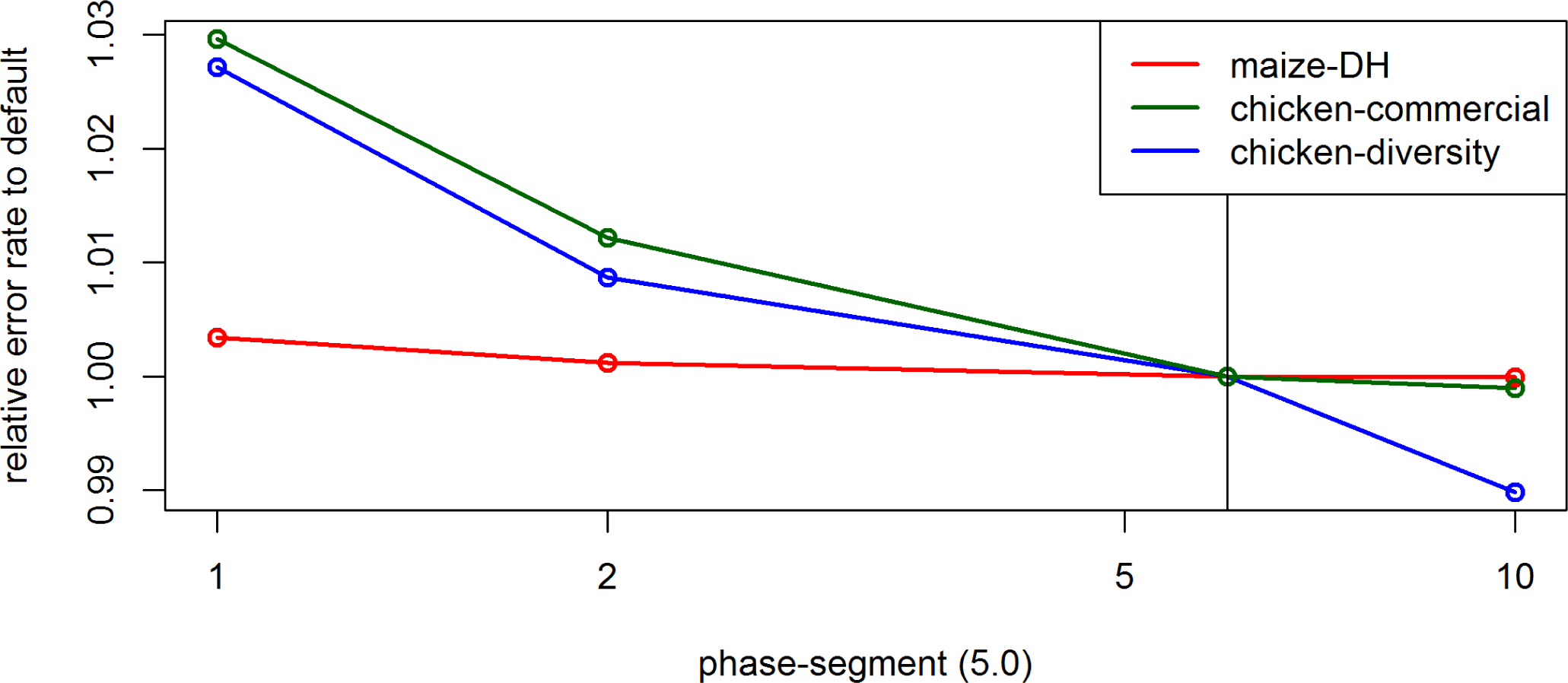
Effect of the parameter *phase-segment* on the UM imputation error rate for the maize data, the commercial chicken line and the chicken diversity panel in BEAGLE 5.0. Default settings are indicated by the vertical line.

**Figure S56.**
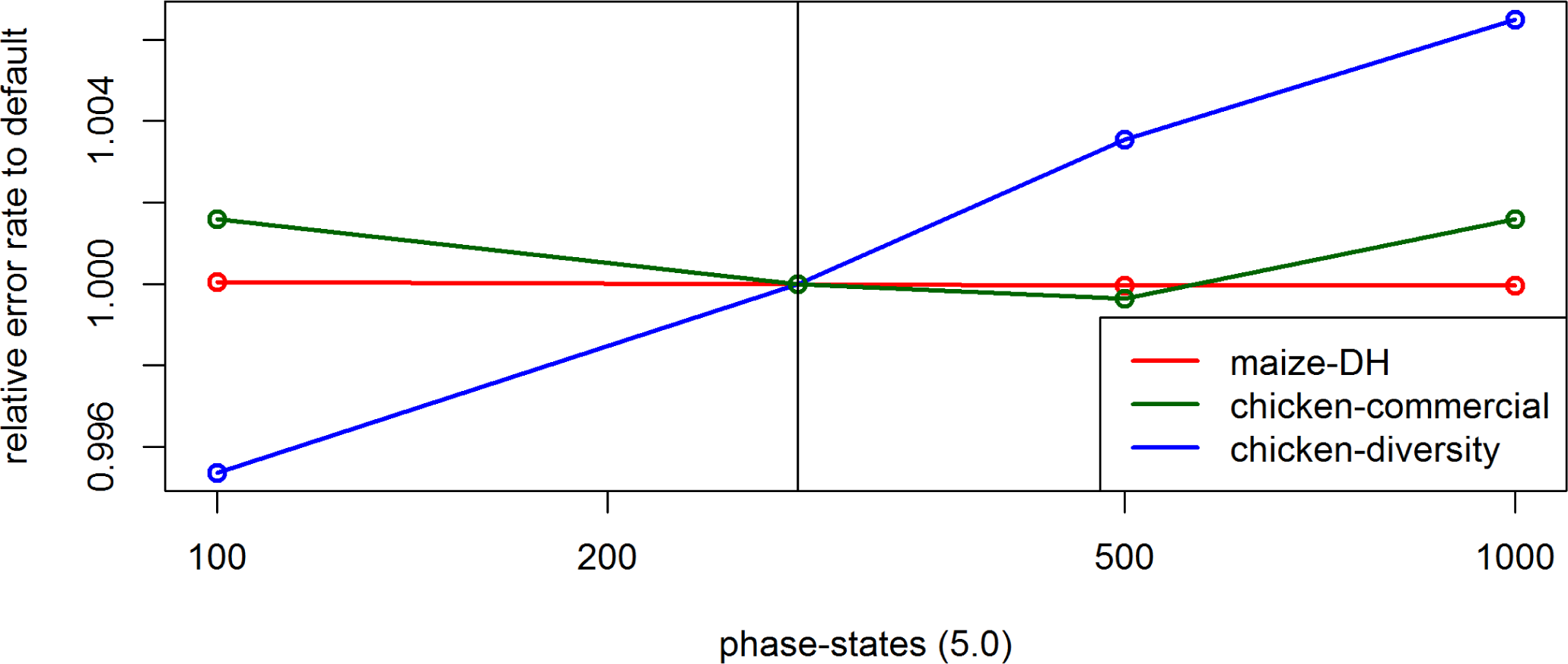
Effect of the parameter *phase-states* on the UM imputation error rate for the maize data, the commercial chicken line and the chicken diversity panel in BEAGLE 5.0. Default settings are indicated by the vertical line.

**Figure S57.**
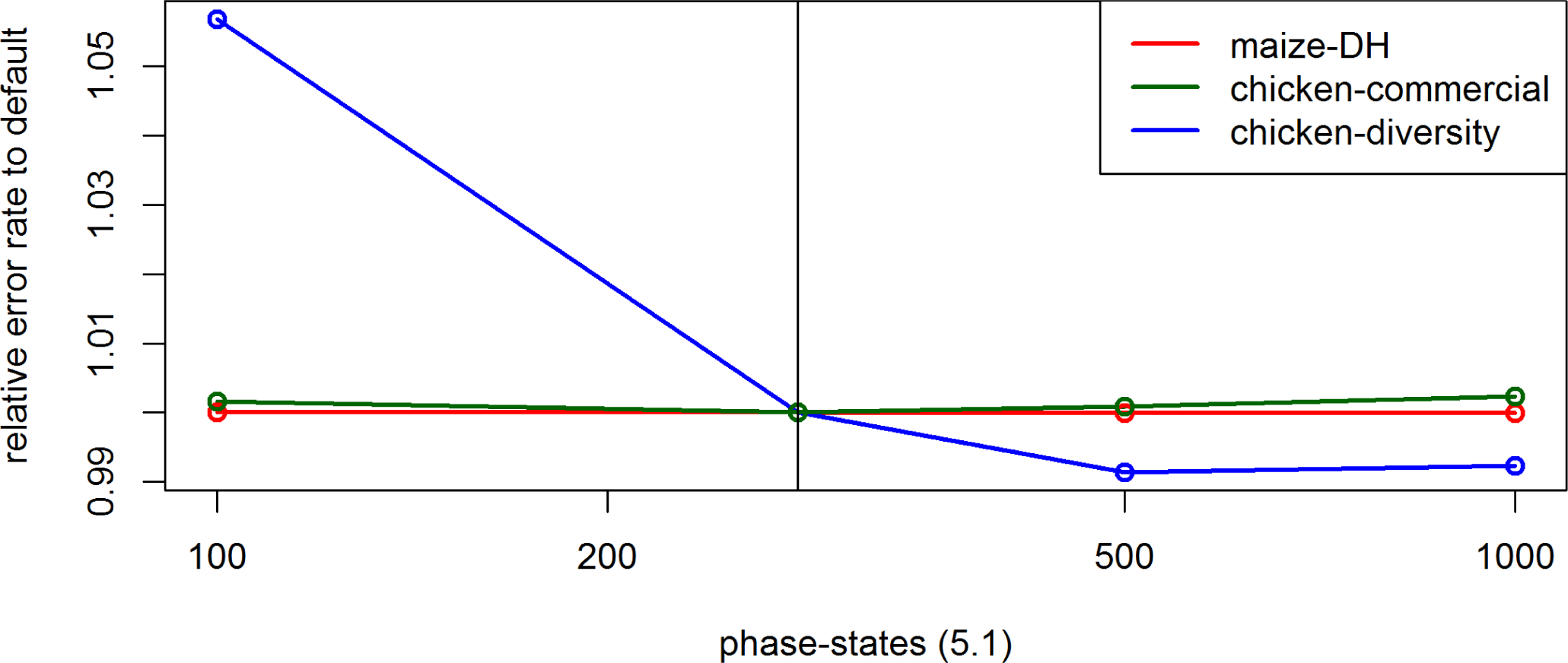
Effect of the parameter *phase-states* on the UM imputation error rate for the maize data, the commercial chicken line and the chicken diversity panel in BEAGLE 5.1. Default settings are indicated by the vertical line.

**Figure S58.**
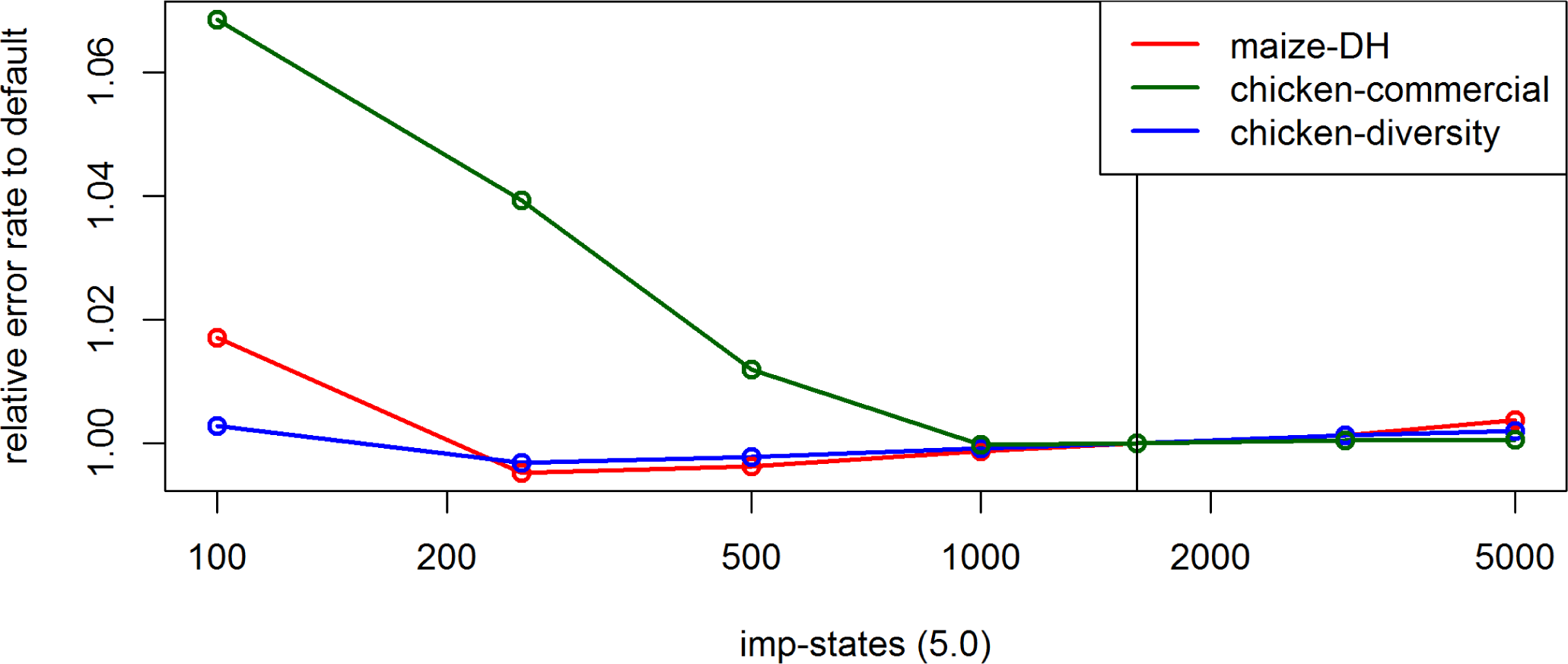
Effect of the parameter *imp-states* on the UM imputation error rate for the maize data, the commercial chicken line and the chicken diversity panel in BEAGLE 5.0. Default settings are indicated by the vertical line.

**Figure S59.**
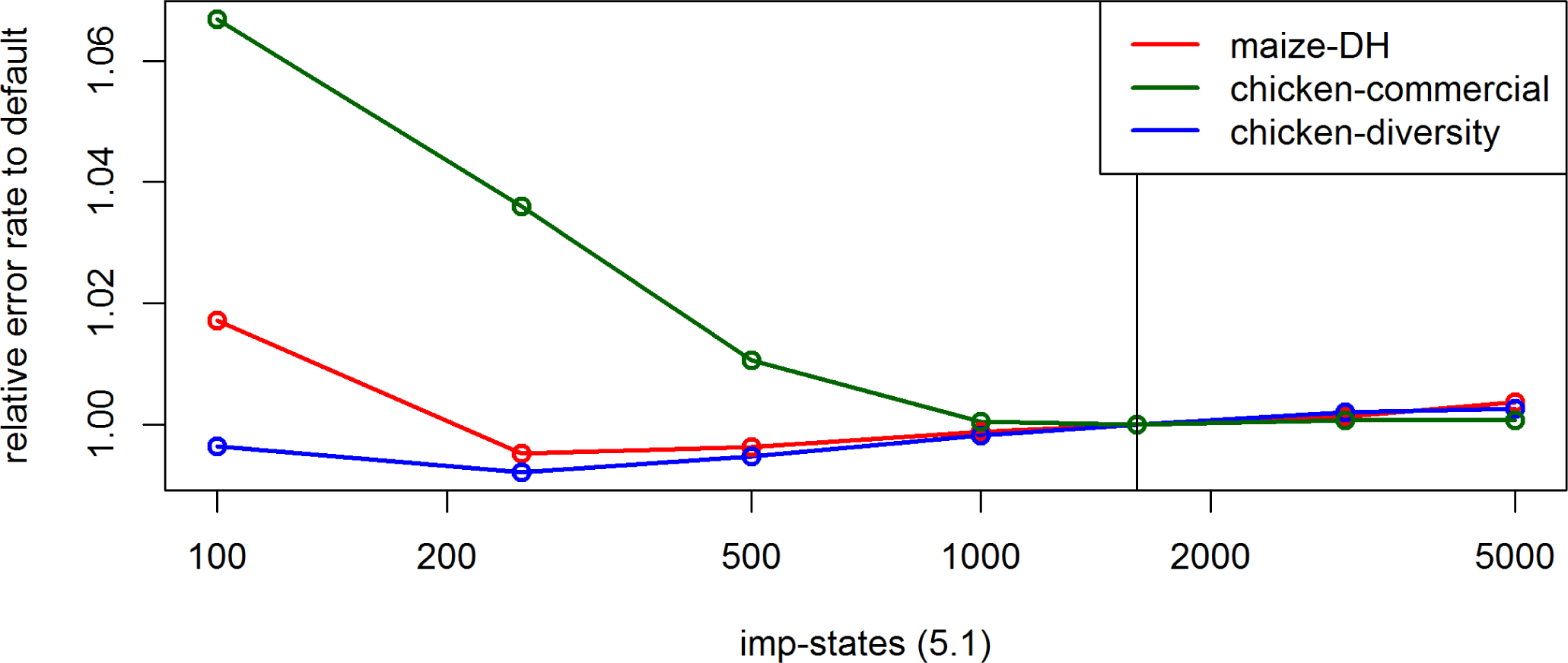
Effect of the parameter *imp-states* on the UM imputation error rate for the maize data, the commercial chicken line and the chicken diversity panel in BEAGLE 5.1. Default settings are indicated by the vertical line.

**Figure S60.**
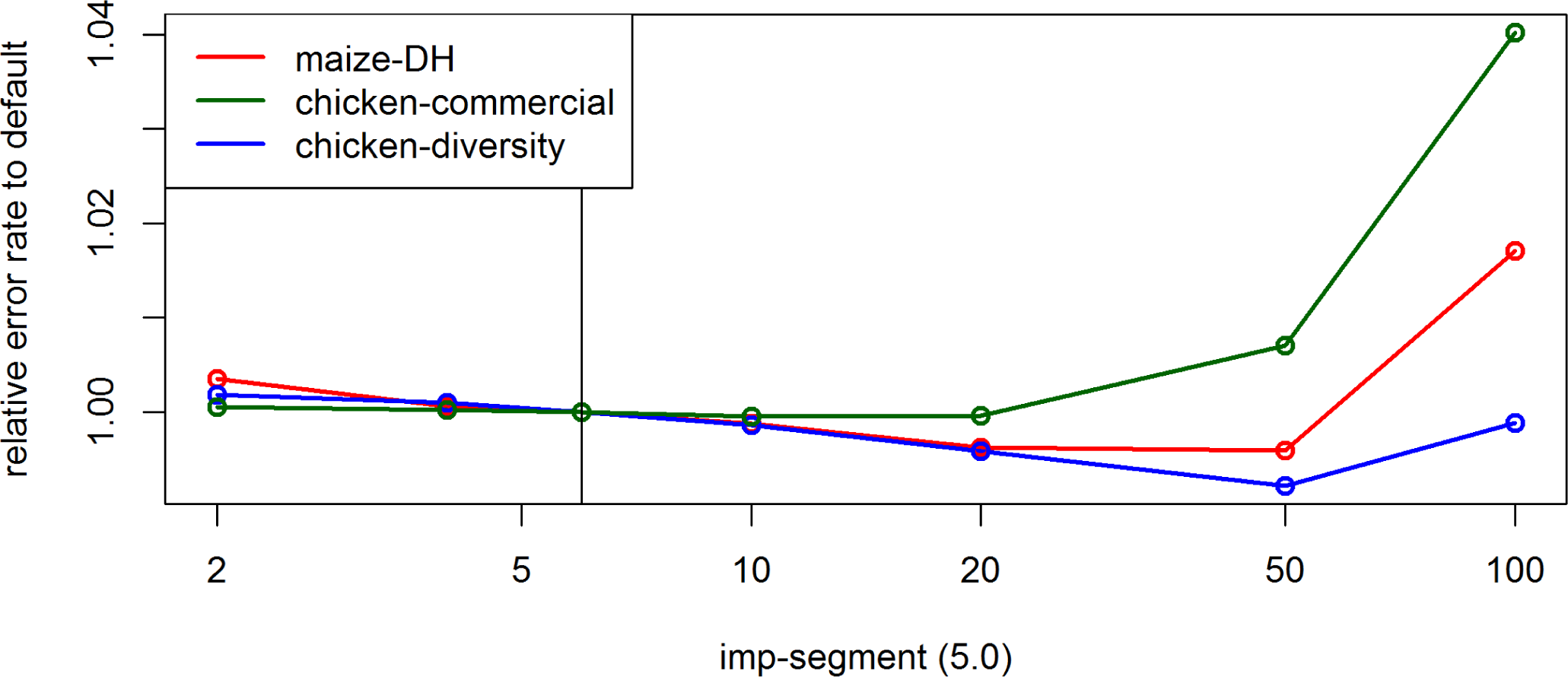
Effect of the parameter *imp-segment* on the UM imputation error rate for the maize data, the commercial chicken line and the chicken diversity panel in BEAGLE 5.0. Default settings are indicated by the vertical line.

**Figure S61.**
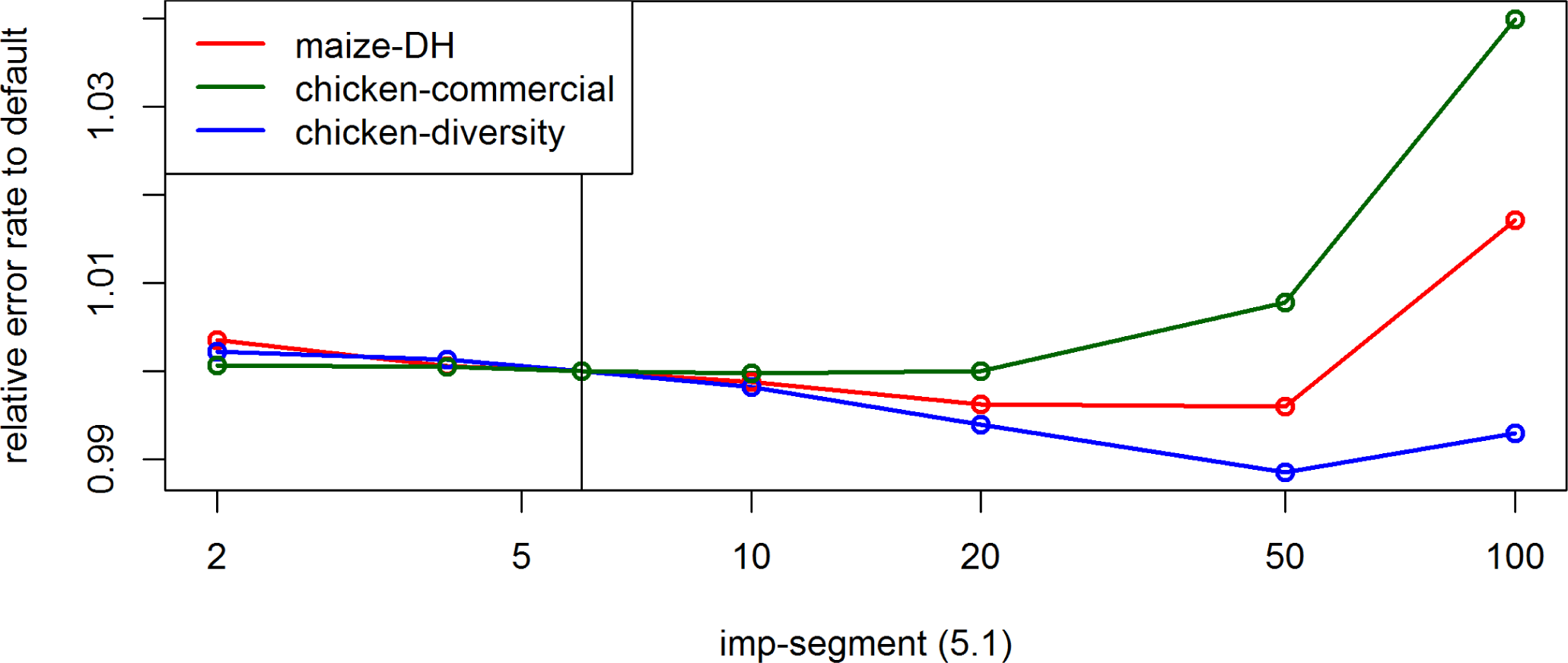
Effect of the parameter *imp-segment* on the UM imputation error rate for the maize data, the commercial chicken line and the chicken diversity panel in BEAGLE 5.1. Default settings are indicated by the vertical line.

**Figure S62.**
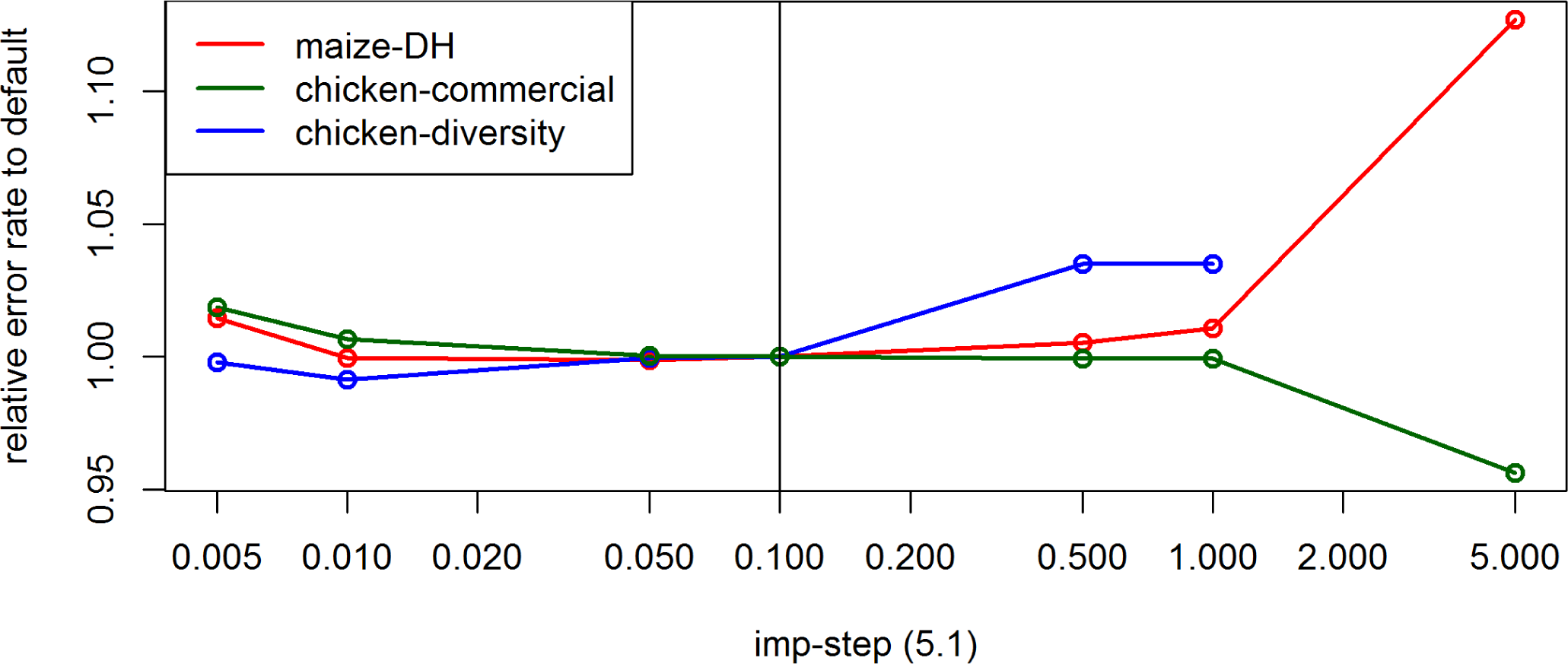
Effect of the parameter *imp-step* on the UM imputation error rate for the maize data, the commercial chicken line and the chicken diversity panel in BEAGLE 5.1. Default settings are indicated by the vertical line.

**Figure S63.**
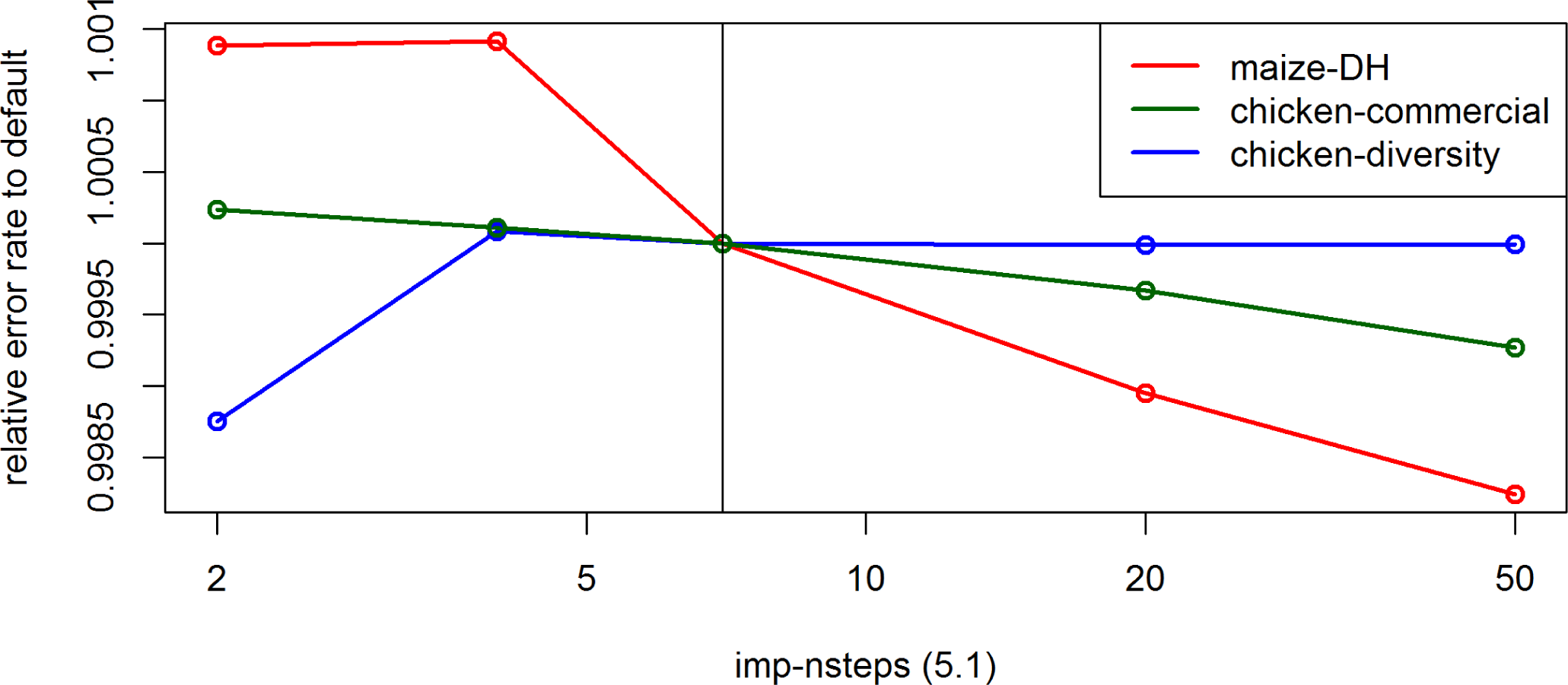
Effect of the parameter *imp-nsteps* on the UM imputation error rate for the maize data, the commercial chicken line and the chicken diversity panel in BEAGLE 5.1. Default settings are indicated by the vertical line.

**Figure S64.**
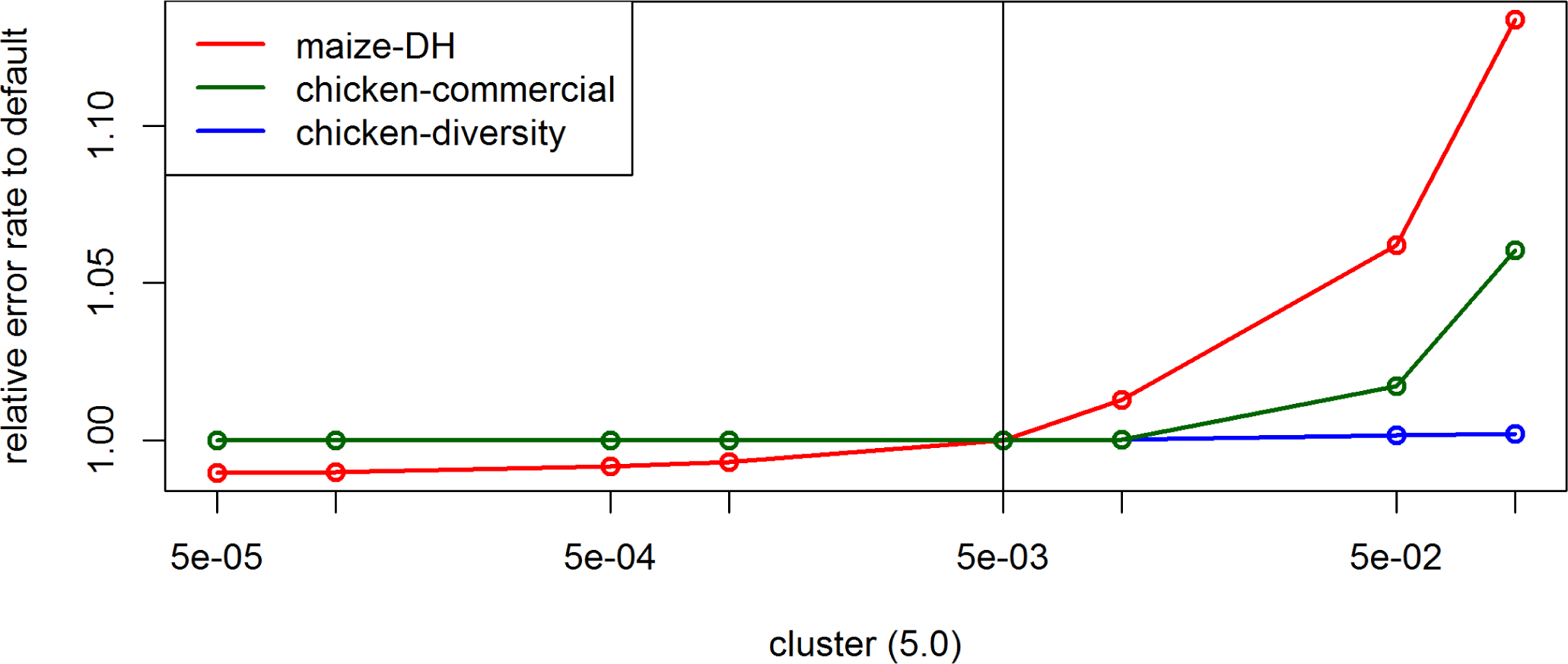
Effect of the parameter *cluster* on the UM imputation error rate for the maize data, the commercial chicken line and the chicken diversity panel in BEAGLE 5.0. Default settings are indicated by the vertical line.

**Figure S65.**
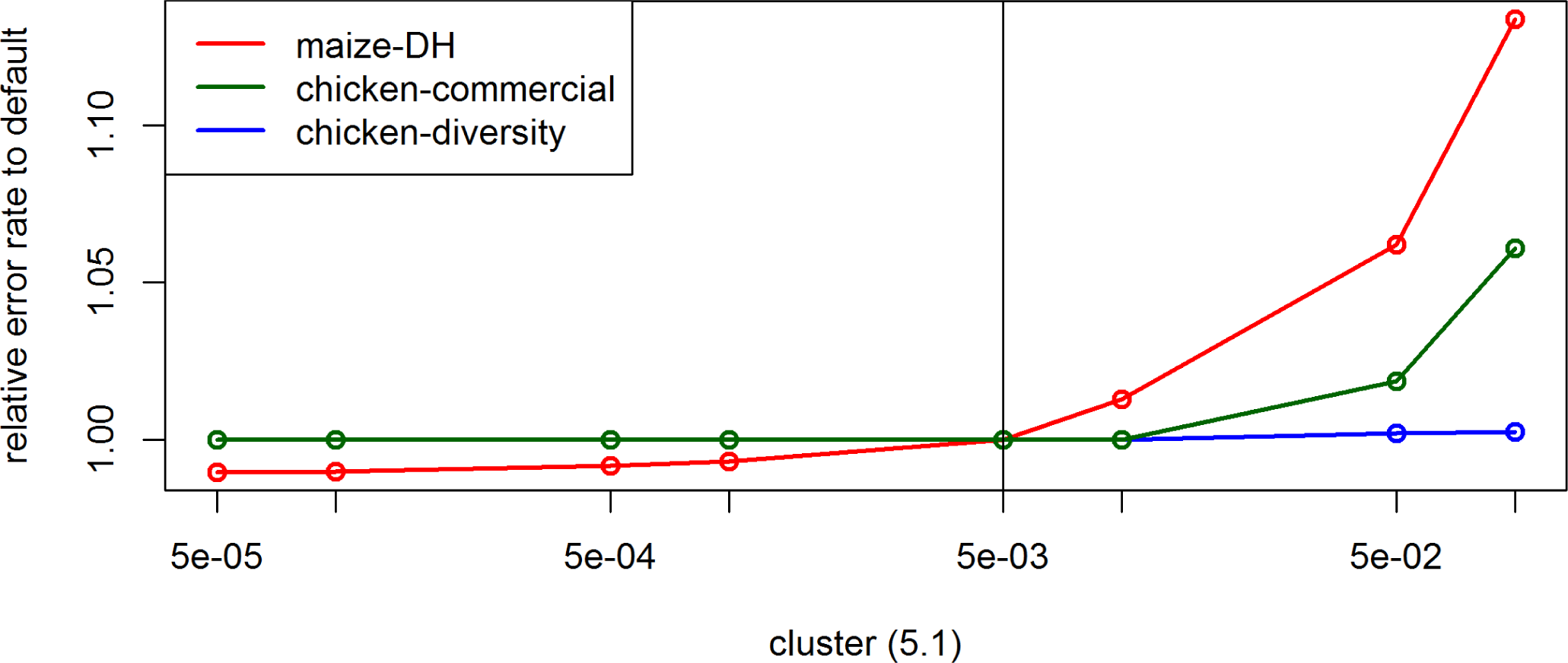
Effect of the parameter *cluster* on the UM imputation error rate for the maize data, the commercial chicken line and the chicken diversity panel in BEAGLE 5.1. Default settings are indicated by the vertical line.

**Figure S66.**
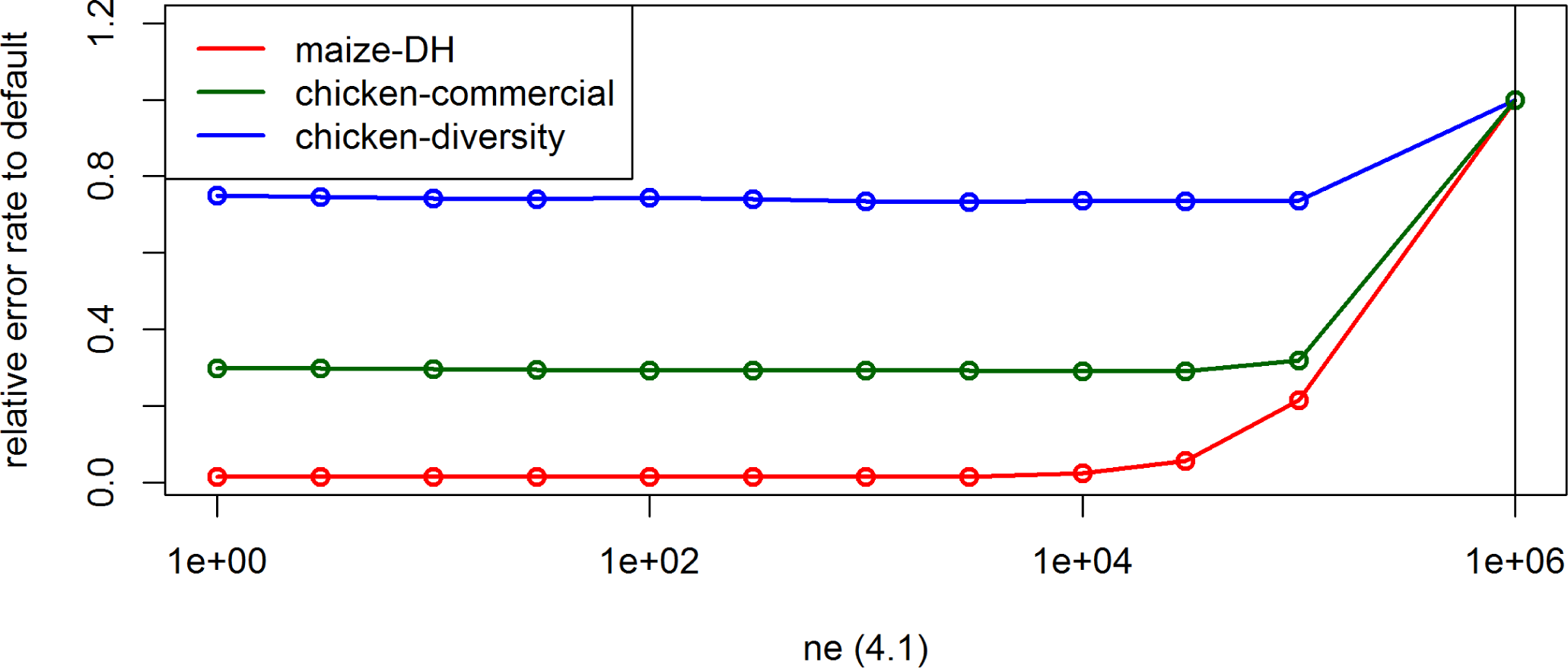
Effect of the parameter *ne* on the UM imputation error rate for the maize data, the commercial chicken line and the chicken diversity panel in BEAGLE 4.1. Default settings are indicated by the vertical line.

**Figure S67.**
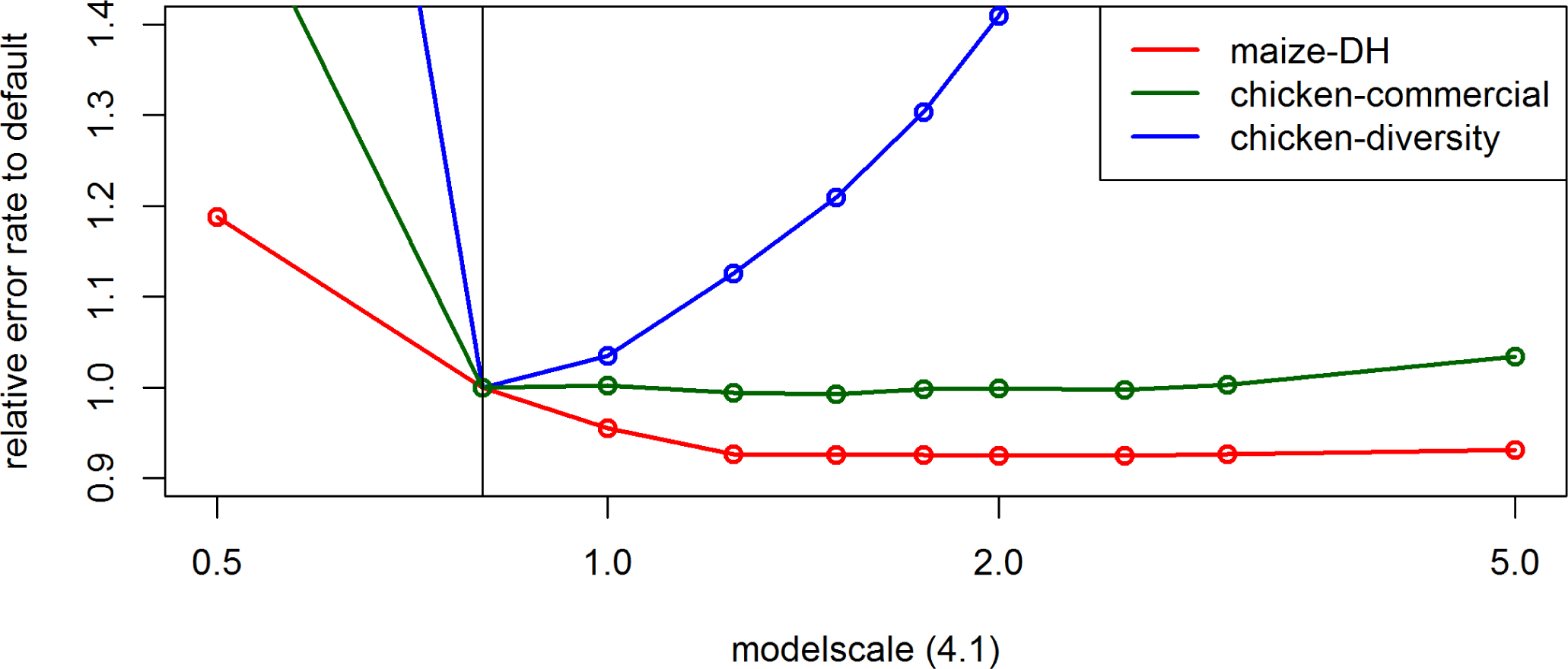
Effect of the parameter *modelscale* on the UM imputation error rate for the maize data, the commercial chicken line and the chicken diversity panel in BEAGLE 4.1. Default settings are indicated by the vertical line.

**Figure S68.**
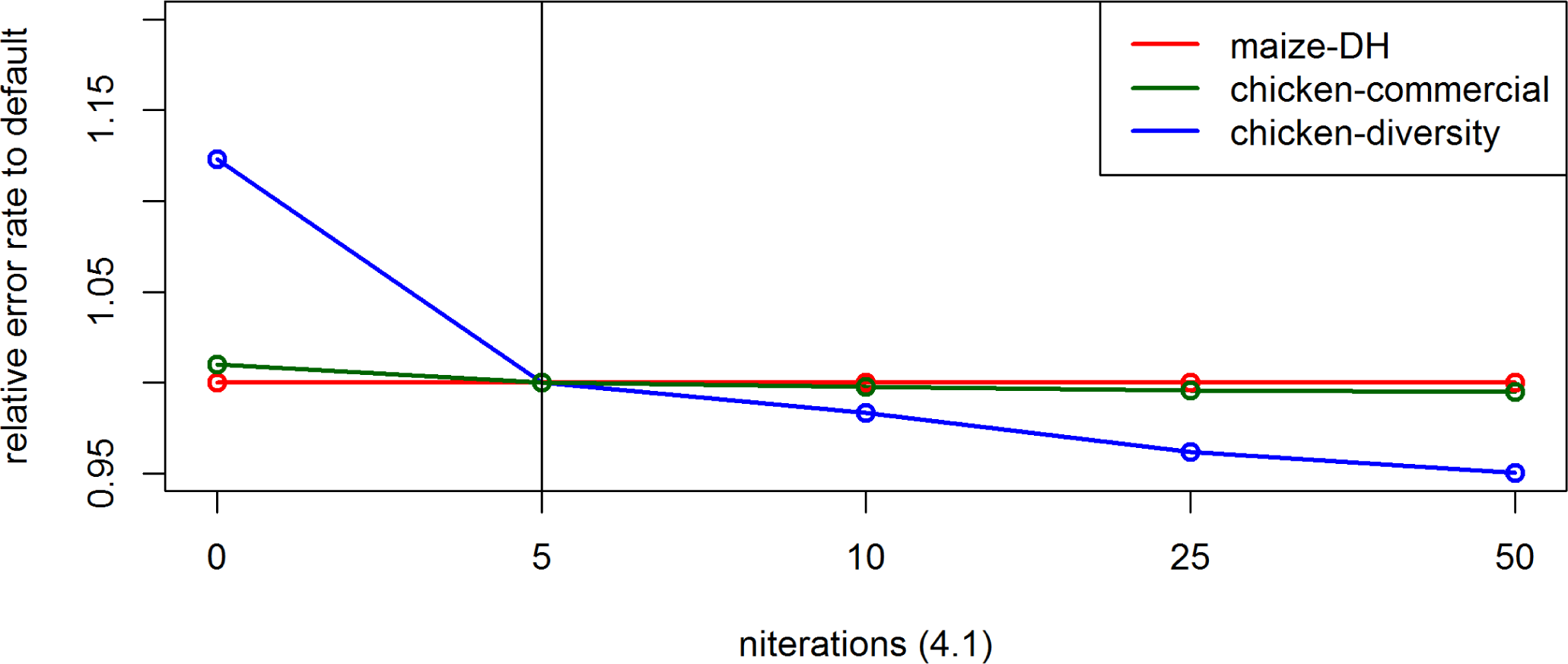
Effect of the parameter *niterations* on the UM imputation error rate for the maize data, the commercial chicken line and the chicken diversity panel in BEAGLE 4.1. Default settings are indicated by the vertical line.

**Figure S69.**
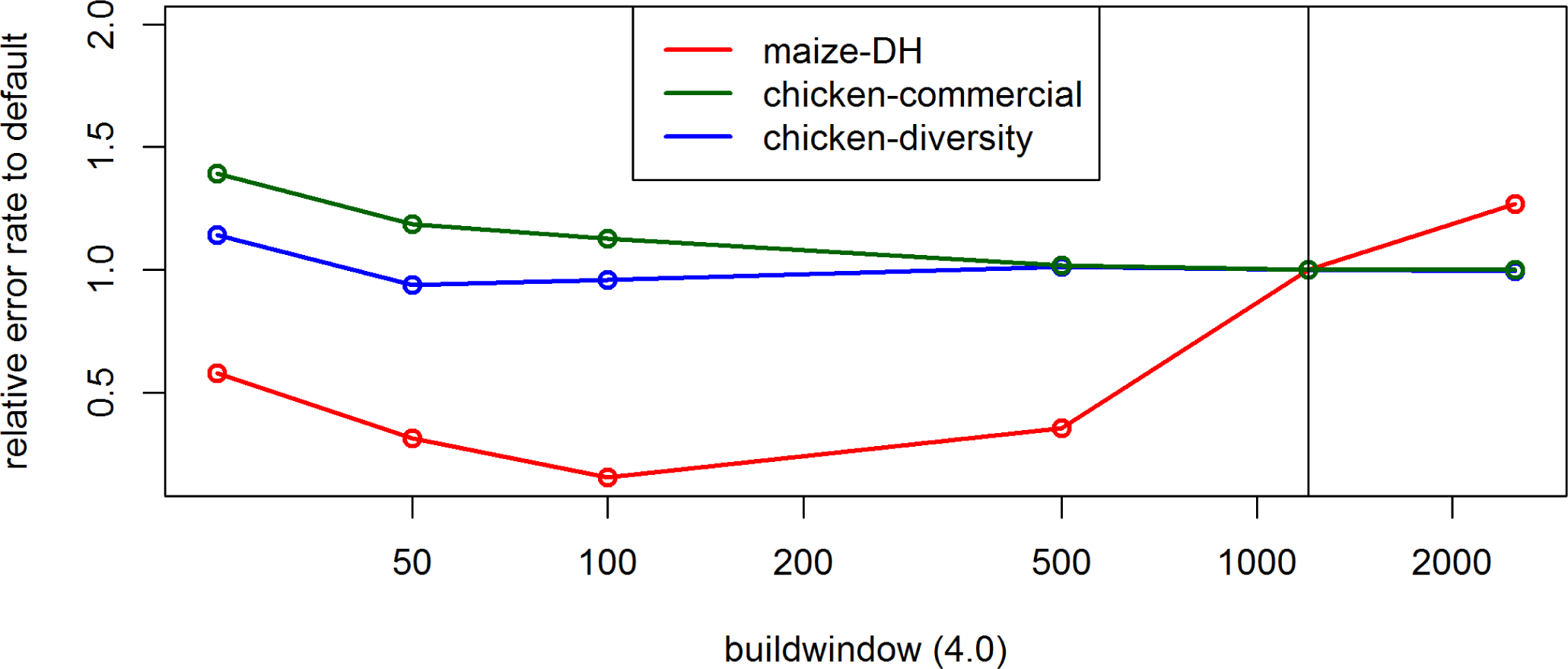
Effect of the parameter *buildwindow* on the UM imputation error rate for the maize data, the commercial chicken line and the chicken diversity panel in BEAGLE 4.0. Default settings are indicated by the vertical line.

**Figure S70.**
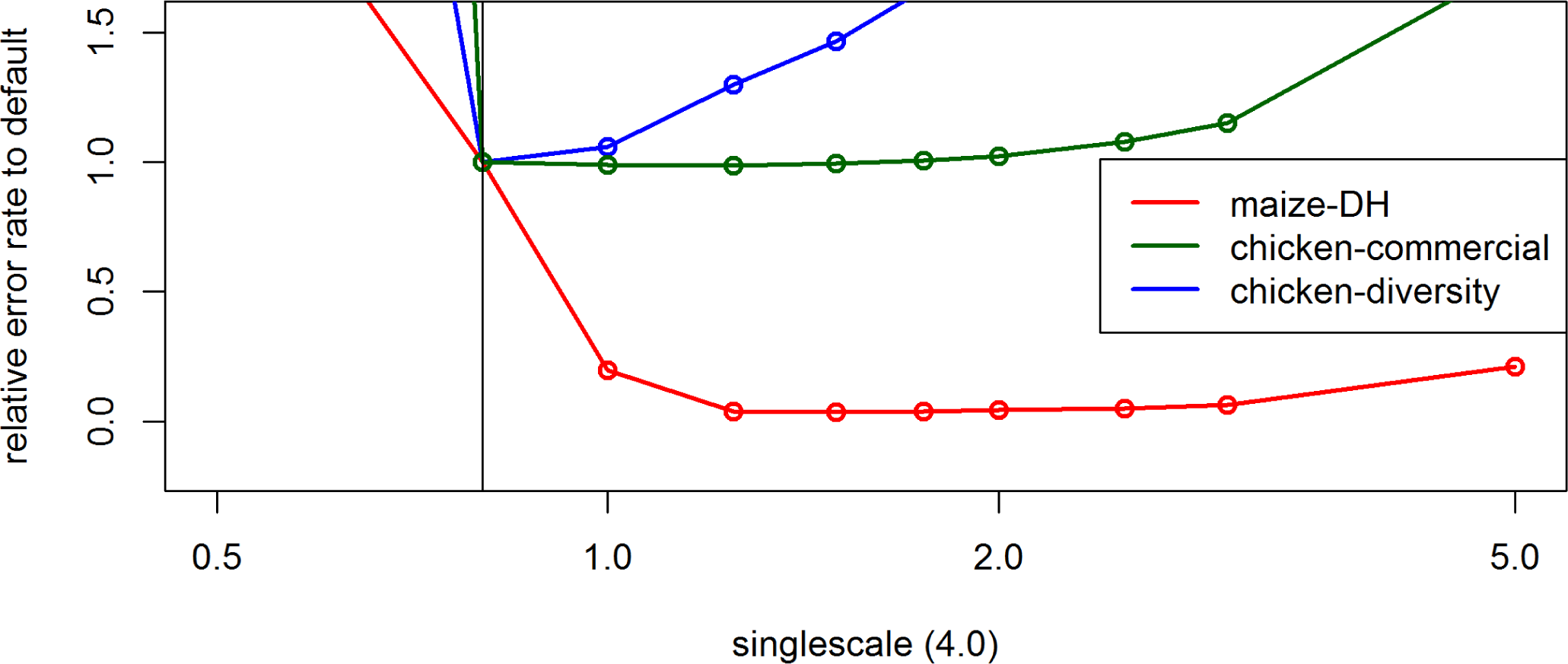
Effect of the parameter *singlescale* on the UM imputation error rate for the maize data, the commercial chicken line and the chicken diversity panel in BEAGLE 4.0. Default settings are indicated by the vertical line.

**Figure S71.**
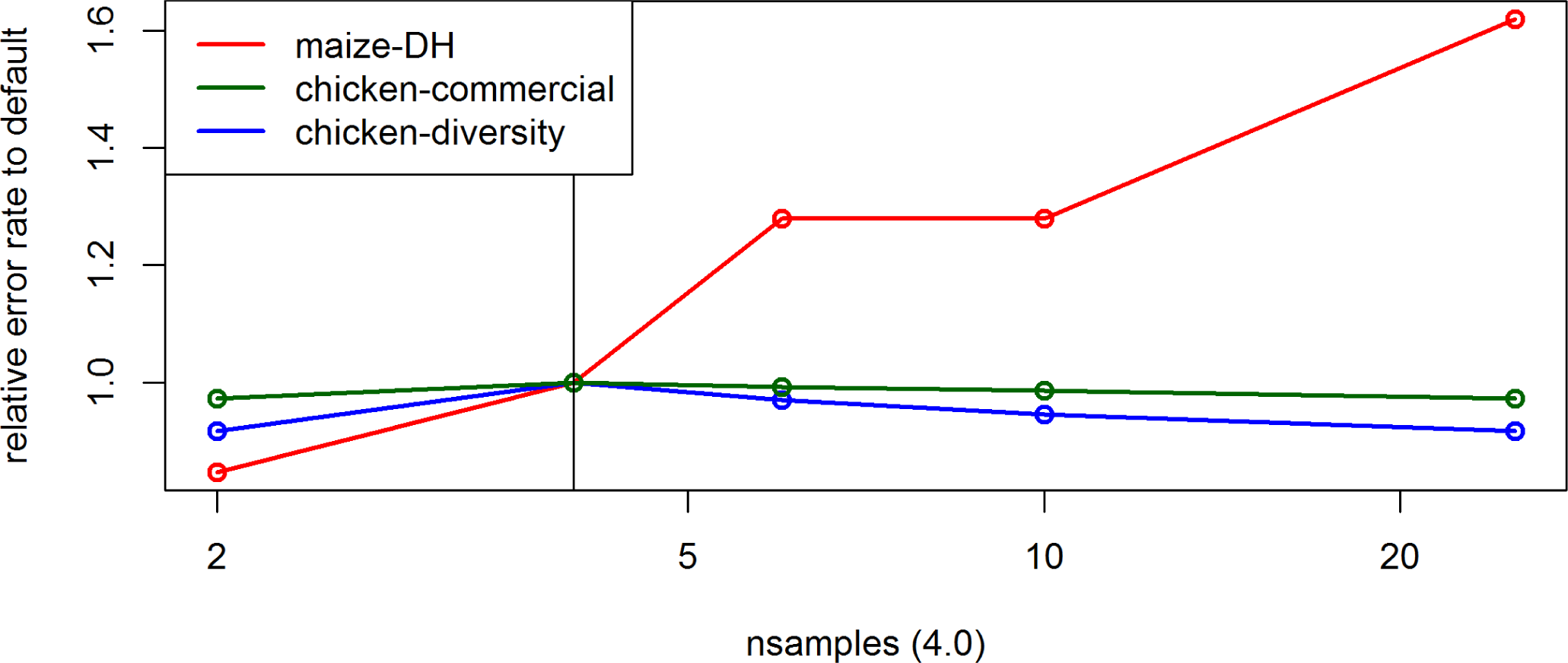
Effect of the parameter *nsamples* on the UM imputation error rate for the maize data, the commercial chicken line and the chicken diversity panel in BEAGLE 4.0. Default settings are indicated by the vertical line.

**Figure S72.**
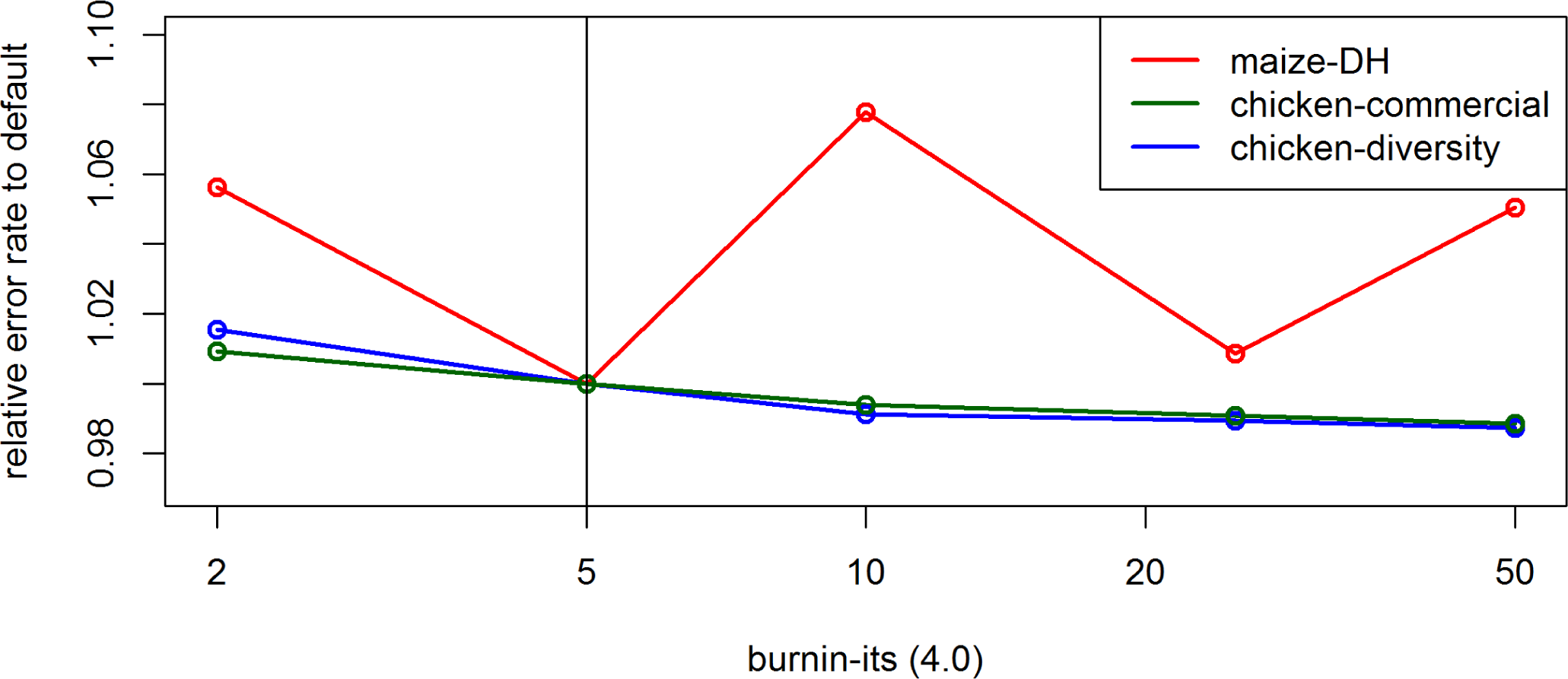
Effect of the parameter *burnin-its* on the UM imputation error rate for the maize data, the commercial chicken line and the chicken diversity panel in BEAGLE 4.0. Default settings are indicated by the vertical line.

**Figure S73.**
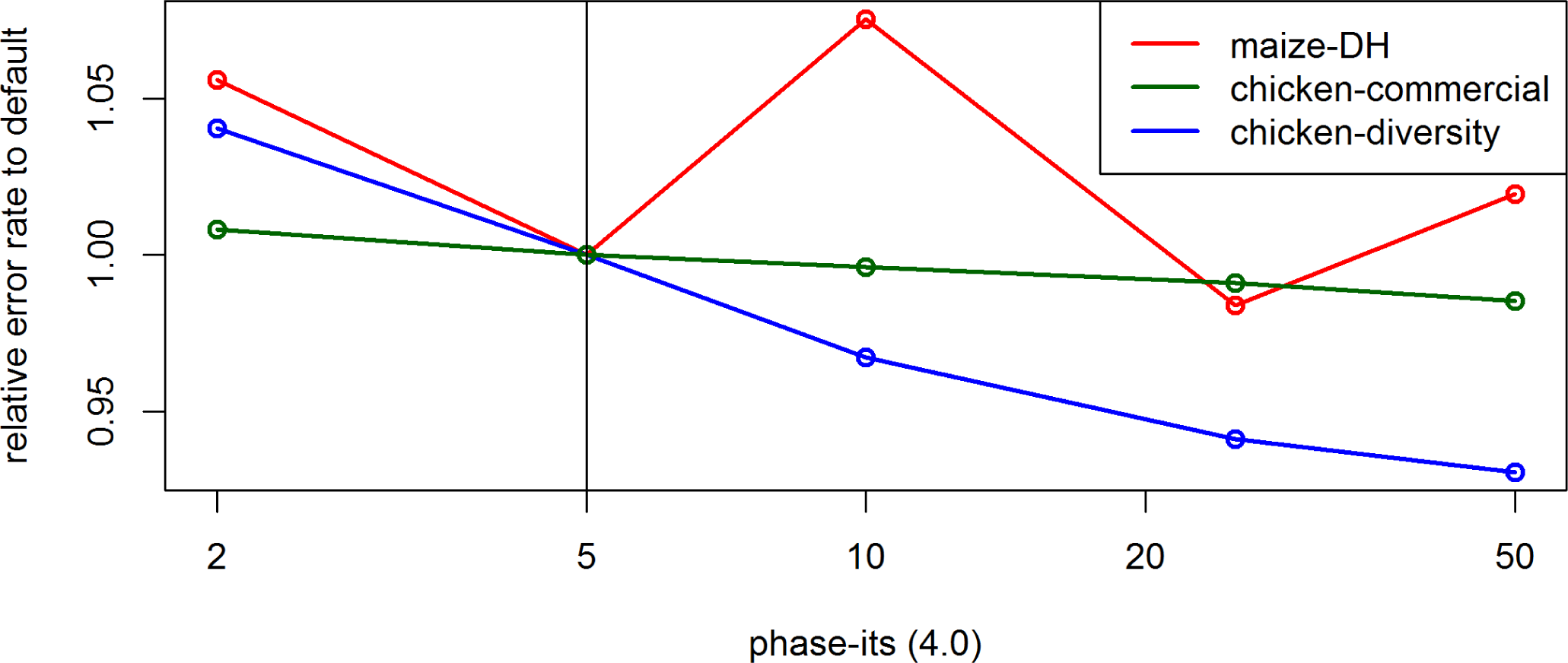
Effect of the parameter *phase-its* on the UM imputation error rate for the maize data, the commercial chicken line and the chicken diversity panel in BEAGLE 4.0. Default settings are indicated by the vertical line.

**Figure S74.**
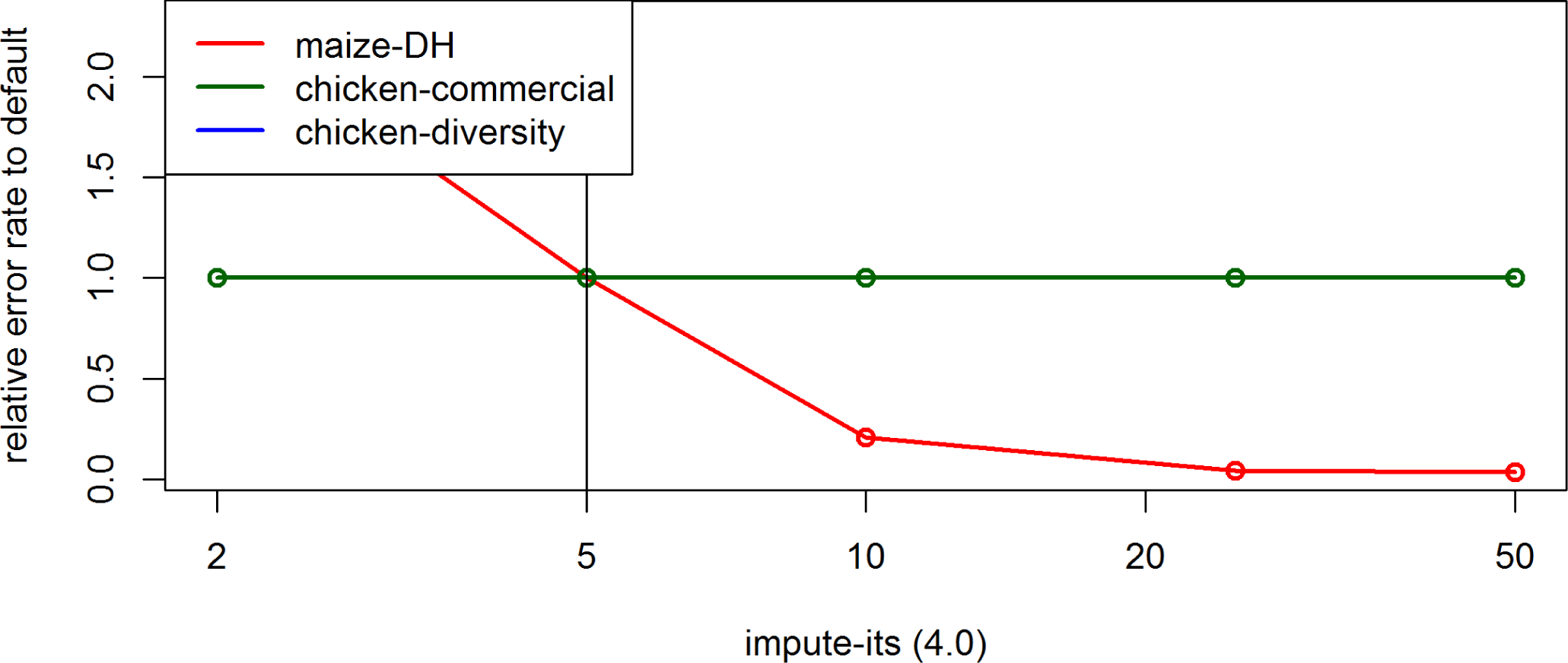
Effect of the parameter *impute-its* on the UM imputation error rate for the maize data, the commercial chicken line and the chicken diversity panel in BEAGLE 4.0. Default settings are indicated by the vertical line.

